# Early postnatal CA3 hyperexcitability drives hippocampal development and epileptogenesis in SCN2A developmental and epileptic encephalopathy

**DOI:** 10.1101/2025.06.29.661458

**Authors:** Yana Reva, Katharina Ulrich, Xian Xin, Yuanyuan Liu, Michaela Barboni, Daniil Kirianov, Nima Mojtahedi, Sarah Schoeb, Hanna Oelßner, Daria Savitska, Mohamad Samehni, Francesca Xompero, Eva Schulze, Heinz Beck, Fabio Morellini, Birgit Engeland, Malte Stockebrand, Stephan Lawrence Marguet, Konstantin Khodosevich, Olga Garaschuk, Tony Kelly, Walid Fazeli, Holger Lerche, Dirk Isbrandt

**Author notes:** Equally contributing authors (Yana Reva, Katharina Ulrich). Correspondence should be addressed to: Prof. Dr. Dirk Isbrandt, DZNE Research Group Experimental Neurophysiology Institute for Molecular and Behavioral Neuroscience University Hospital Cologne, Kerpener Str. 62, 50937 Köln.

## Abstract

Developmental and epileptic encephalopathies caused by pathogenic variants in *SCN2A* (*SCN2A*-DEE), encoding the voltage-gated sodium channel Na_v_1.2, present with early-life seizures, developmental delay, and increased mortality. Using a novel *Scn2a* p.A263V gain-of-function (GOF) mouse model, we demonstrate gene-dose and background-dependent phenotypes ranging from self-limited neonatal seizures to chronic epilepsy with high mortality. In vivo electrophysiology revealed hippocampal seizures as early as postnatal day 2.5, with CA3-driven gamma oscillations preceding seizure onset. CA3 and CA1 pyramidal neurons exhibited transient hyperexcitability during early postnatal development, resolving by P24-30. Single-cell RNA sequencing uncovered gene dose-dependent accelerated maturation of hippocampal networks, peaking at P7, alongside widespread transcriptional changes in excitatory and inhibitory neurons. In adulthood, persistent hippocampal network alterations emerged, marked by reduced mid-gamma oscillations and theta-gamma coupling. Our findings establish hippocampal CA3 hyperexcitability as an early driver of epileptogenesis in *SCN2A*-DEE and highlight it as a potential therapeutic target to mitigate disease progression.

## Introduction

Developmental and epileptic encephalopathies (DEEs) are a genetically heterogeneous group of diseases with early onset, limited treatment options, and significant lifelong impact, including developmental delay, intellectual disability (ID), behavioral abnormalities, and pharmacoresistant epileptic seizures. Approximately 25% of DEE-associated genes encode ion channels^1^, including *SCN2A*, a channel gene also associated with neuropsychiatric disorders^2^. *SCN2A* encodes the alpha subunit of Na_v_1.2, a sodium channel crucial for the initiation, propagation, and backpropagation of neuronal action potentials^3–6^. Na_v_1.2 is the dominant axonal sodium channel isoform during human and mouse brain development^7^. Inherited or de novo pathogenic *SCN2A* variants are strongly associated with neurodevelopmental disorders. Loss-of-function (LOF) variants are linked to autism and ID, with or without epilepsy^8–12^, whereas gain-of-function (GOF) mutations are associated with neonatal or early-infantile onset epilepsy syndromes. These syndromes range from mild, self-limited familial neonatal-infantile epilepsy (SeLFNIE) to severe *SCN2A*-DEE^11^.

As in humans, GOF and LOF variants in mouse *Scn2a* lead to neurodevelopmental disorders, with *Scn2a* GOF mutations causing epilepsy in both species^13,14^. In early adulthood, mice haploinsufficient for Scn2a or harboring LOF mutations show ASD-like phenotypes^15,16^ and develop non-convulsive absence-like seizures with spike-wave discharges (SWDs) later in life^15,17^.

Despite that it is established that Na_v_1.2 GOF mutations cause generalized behavioral seizures and sudden unexpected death (SUDEP)^18^, the networks implicated in epileptogenesis in the developing brain are unknown. Na_v_1.2 plays multiple, evolving roles in mediating neuronal and network excitability across development. In the first two weeks of postnatal mouse brain development, Na_v_1.2 is highly expressed in the axons of glutamatergic pyramidal neurons, where it is a key factor in somatic action potential (AP) generation^3,19^. As development progresses, Na_v_1.2 channels become confined to the proximal axon initial segment (AIS), while Na_v_1.6 expression increases at the distal AIS^19^. At this stage, Na_v_1.6 governs AP initiation, while Na_v_1.2 channels promote AP invasion of the soma from the AIS^3^. In addition, Na_v_1.2 is involved in dendritic excitability and AP back-propagation in mature cortical pyramidal neurons^5^. Beyond excitatory neurons, Na_v_1.2 is also expressed in caudal ganglionic eminence (CGE)-derived disinhibitory interneurons, suggesting a role in inhibitory neuronal circuit control^20^.

The dominant involvement of Na_v_1.2 during development and its functional replacement at the AIS by Na_v_1.6 suggest that the characterization of cellular and network dysfunction associated with Na_v_1.2 GOF during early postnatal development may be vital to understanding epileptogenesis and associated behavioral comorbidities. We therefore engineered *SCN2A* GOF mouse models based on one of the most recurrent human pathogenic *SCN2A* variants, i.e., p.Ala263Val (p.A263V). This variant is associated with a broad clinical spectrum of variable severity, including SeLFNIE and childhood-onset episodic ataxia^20–23^, severe ID, early infantile epileptic encephalopathy (EIEE, Ohtahara syndrome), intractable neonatal-onset epileptic encephalopathy, and increased mortality^11^. Using mice heterozygous or homozygous for the *Scn2a*(p.A263V) variant on two different genetic backgrounds, we recapitulated the clinical variability observed in *SCN2A*-linked DEE. This approach revealed a gene dose-dependent phenotypic spectrum, ranging from transient neonatal epilepsy without persistent sequelae to a chronic epilepsy phenotype with high mortality. Our multidisciplinary experimental strategy, encompassing the characterization of neonatal and infantile intrinsic neuronal properties, spontaneous in situ and in vivo network activities, and developmental transcriptomic profiling of hippocampal maturation, identified hippocampal CA3 hyperexcitability as an early driver of epileptogenesis in *SCN2A*-DEE.

## Results

### *Scn2a* p.A263V mutation elicits gene dose-dependent seizures, mortality, and developmental changes

We generated a genetic mouse model of Na_v_1.2 channelopathy by introducing the human *SCN2A*(p.A263V) pathogenic sequence variant into the *Scn2a* locus through homologous recombination in embryonic stem cells^21^. When expressed in a heterologous expression system, Na_v_1.2(p.A263V) subunits mediated an increased tetrodotoxin-sensitive persistent sodium current, confirming the gain-of-function (GOF) nature of this pathogenic sequence variant^22^. To achieve a comprehensive phenotypic characterization, we established the model on the C57BL/6J (*B6.Scn2a*) and CD1 (*CD1.Scn2a*) backgrounds. Throughout this study, we refer to *Scn2a^A263V/+^* mice as *+/mut* and *Scn2a^A263V/A263V^*mice as *mut/mut*.

To investigate whether the p.A263V variant recapitulates the epilepsy phenotype observed in patients, we conducted chronic telemetric video electrocorticogram (ECoG) recordings from the sensorimotor cortical region of at least six-week-old animals. Both *CD1.Scn2a* and *B6.Scn2a mut/mut* mice exhibited spontaneous generalized tonic-clonic seizures (Fig. 1a and Extended Data Fig. 1a). Seizures typically occurred during sleep transitions, beginning with a tonic phase followed by amplitude suppression and a generalized tonic-clonic phase. However, not all seizures exhibited each phase (Fig. 1b, Supplementary Video 1). Each seizure event was followed by postictal depression. Analysis of pooled data from *B6.Scn2a* and *CD1.Scn2a mut/mut* mice revealed a frequency of 0.7 [0.3 2.1] (median [Q1 Q3]) seizures per day, with episodes lasting 72.2 [29.3 89.6] s (Fig. 1c), and 82% of mice experienced seizures within the recording time frame. Individual seizure phases lasted approximately 22.0-26.3 seconds each (Fig. 1d), while postictal depression persisted for 178.4 [129.6 233.2] s (Fig. 1e).

**Fig. 1.**
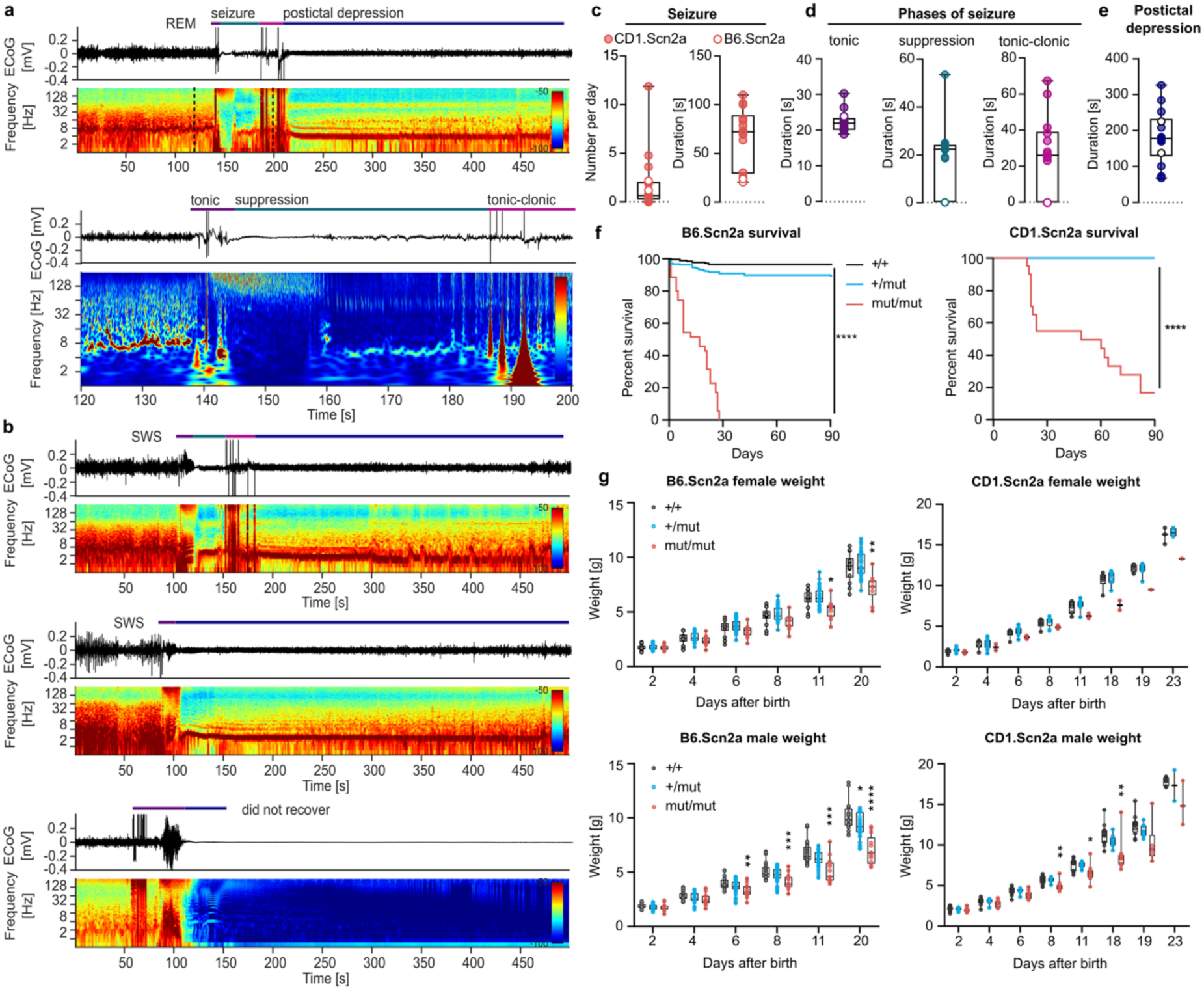
| Spontaneous seizures, increased mortality, and decreased body weight in *Scn2a* mice harboring p.A263V mutation. **a,** Representative spontaneous seizure in an adult *CD1.Scn2a mut/mut* mouse. Top: Raw electrocorticogram (ECoG) trace and spectrogram of a seizure episode, including onset from REM sleep and subsequent postictal depression (dashed lines indicate magnified area). Bottom: Magnified view of the seizure showing progression from the tonic phase through amplitude suppression to the tonic-clonic phase. **b,** Seizure variations in *CD1.Scn2a mut/mut* mice. Top: Seizure following slow-wave sleep. Middle: Seizure with tonic phase only. Bottom: Fatal seizure episode. All seizures were followed by postictal depression. **c-e,** Seizure characteristics in *B6.Scn2a* and *CD1.Scn2a mut/mut* mice (*n =* 17, *B6.Scn2a* = 2 (open circles), *CD1.Scn2a* = 15 (closed circles)). **c,** Left: Average seizures per day. Right: Mean seizure duration. **d,** Duration of seizure phases: Tonic (left), amplitude suppression (middle), tonic-clonic (right). **e,** Mean postictal depression duration. **f,** Survival curves for *B6.Scn2a* (left, *n =* 544, *+/+* = 266, *+/mut* = 242, *mut/mut* = 36) and *CD1.Scn2a* (right, *n =* 130, *+/+* = 31, *+/mut* = 79, *mut/mut* = 20) lines. Log-rank (Mantel-Cox) test, ****P < 0.0001. **g,** Body weight trajectories in *B6.Scn2a* mice: Females (top left, *n =* 57, *+/+* = 14, *+/mut* = 32, *mut/mut* = 11), males (bottom left, *n =* 75, *+/+* = 23, *+/mut* = 39, *mut/mut* = 13), and *CD1.Scn2a* mice: Females (top right, *n =* 17, *+/+* = 6, *+/mut* = 9, *mut/mut* = 2), males (bottom right, *n =* 34, +/+ = 13, *+/mut* = 10, *mut/mut* = 11). Mixed-effect analysis with Dunnett’s multiple comparison test, \**P* < 0.05, \*\**P* < 0.01, \*\*\**P* < 0.001.

In addition to seizures, *mut/mut* mice displayed other signs of aberrant excitability. In *B6.Scn2a mut/mut* mice, we observed frequent polymorphic spikes, spike-and-wave discharges, and spike trains during interictal periods. Some *mut/mut* animals also showed low-voltage background activity with a burst-suppression ECoG pattern as previously reported for mice with GOF mutations in *Scn2a*^13^. While *CD1.Scn2a +/mut* mice showed no apparent ECoG abnormalities, *B6.Scn2a +/mut* animals exhibited occasional spikes and short spike-wave discharges (Extended Data Fig. 1b).

During our ECoG recordings, 18% of *mut/mut* mice invariably died during the postictal depression phase following a seizure event (Fig. 1b, bottom). Subsequent analysis of overall survival revealed increased mortality in *mut/mut* mice but not in *+/mut* mice (Fig. 1f). The effect was line-dependent, that is, *B6.Scn2a mut/mut* mice showed 100% mortality by postnatal day 30 (P30), while 55% of *CD1.Scn2a mut/mut* mice survived beyond this point. The *mut/mut* mice that spontaneously died were often found in a characteristic body posture (forelimb, hindlimb, and tail extension), suggesting that they had experienced a behavioral seizure immediately followed by death.

The mortality phenotype was accompanied by altered body weight development. Both *+/mut* and *mut/mut* newborn mice had normal weights at birth. Later, the body weight trajectories of *mut/mut* mice progressively deviated from those of *+/mut* or *+/+* mice (Fig. 1g). This effect was persistent in *B6.Scn2a mut/mut* mice of both sexes and transient (P8-P19) only in male *CD1.Scn2a mut/mut* mice. Altogether, our *Scn2a* p.A263V mouse models recapitulate key features of severe chronic epilepsy observed in patients, including spontaneous seizures and increased mortality. They exhibit an apparent gene dose effect, with *+/mut* mice presenting milder phenotypes than their *mut/mut* counterparts and showing background-dependent variations in phenotype severity.

**Extended Data Fig. 1.**
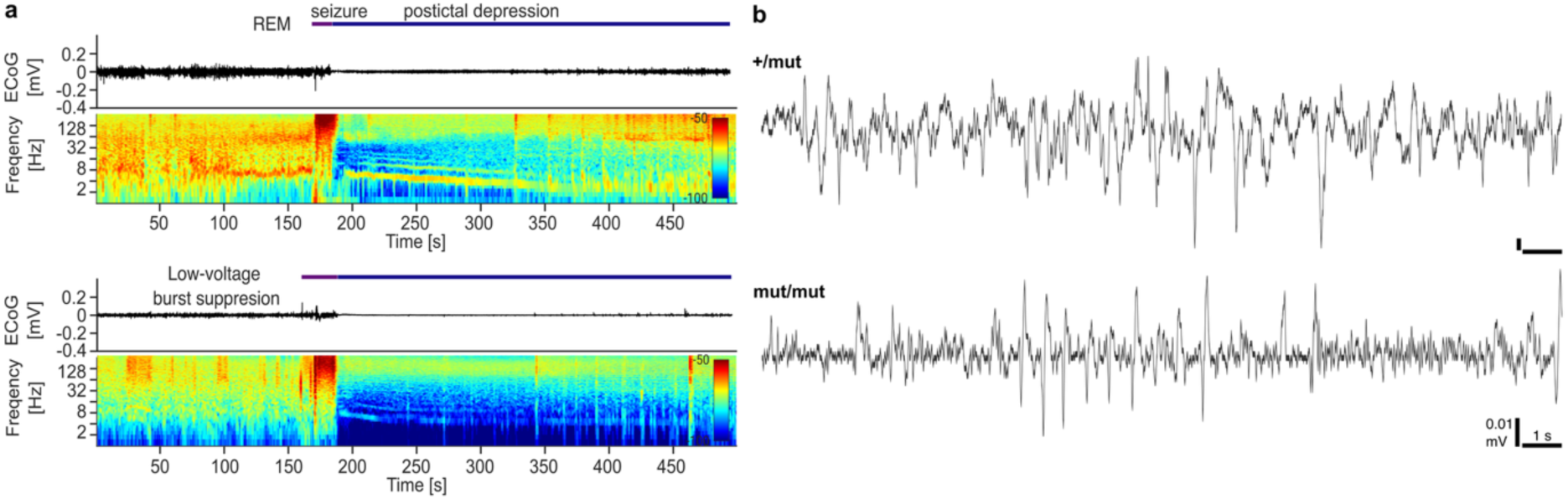
| Spontaneous seizures in *B6.Scn2a* mice harboring p.A263V mutation. **a**, Spontaneous seizure in an adult *B6.Scn2a mut/mut* mouse. Top: Raw ECoG trace and spectrogram of a seizure episode, including onset from REM sleep and subsequent postictal depression. Bottom: Low-voltage burst suppression ECoG interrupted by a short spike-and-wave seizure, followed by postictal depression. **b,** Representative interictal spike trains in *+/mut* (top) and *mut/mut* (bottom) mice.

### Severe seizure phenotype in *Scn2a* p.A263V mice does not induce major brain morphological changes

Chronic epilepsy is often associated with hippocampal alterations, such as neuronal loss, reactive astrogliosis, and inflammation^23–25^. We analyzed brain size and various hippocampal markers in *CD1.Scn2a* mice to investigate potential structural changes. When comparing the brain size using bregma-lambda distance measurements, no significant differences were observed between genotypes across multiple developmental time points (P2-3, P6-7, P13-14, and adults) (Fig. 2a-b). To evaluate potential astrogliosis, we examined glial fibrillary acidic protein (GFAP) immunoreactivity at P21. Quantification of GFAP immunoreactivity showed comparable levels across genotypes in all hippocampal regions (Fig. 2c-d), indicating no substantial astrogliosis. We then assessed perineuronal nets (PNNs) using Wisteria floribunda agglutinin (WFA) staining. While the overall staining intensity was similar across genotypes (Fig. 2e-f), WFA-positive cell counts were reduced in *mut/mut* mice across all hippocampal regions (Fig. 2g). Adult *CD1.Scn2a* mice showed no differences in WFA staining (Extended Data Fig. 2a-b), suggesting age-dependent effects. Given the observed alterations in PNNs, we next investigated parvalbumin (PV) immunoreactivity in adult *CD1.Scn2a* mice because PV-positive interneurons are often surrounded by these extracellular matrix structures. We found decreased PV staining intensities in *mut/mut* animals throughout the hippocampus, especially in CA1, CA2, and CA3, with no change in PV-positive cell counts (Extended Data Fig. 2c-e). To further investigate potential epilepsy-related changes in adult *CD1.Scn2a* mice, we examined neuropeptide Y (NPY) immunoreactivity, a marker known to be upregulated in epilepsy^24^. NPY staining intensity was increased in the dentate gyrus of *mut/mut* mice (Extended Data Fig. 2f-g).

**Fig. 2.**
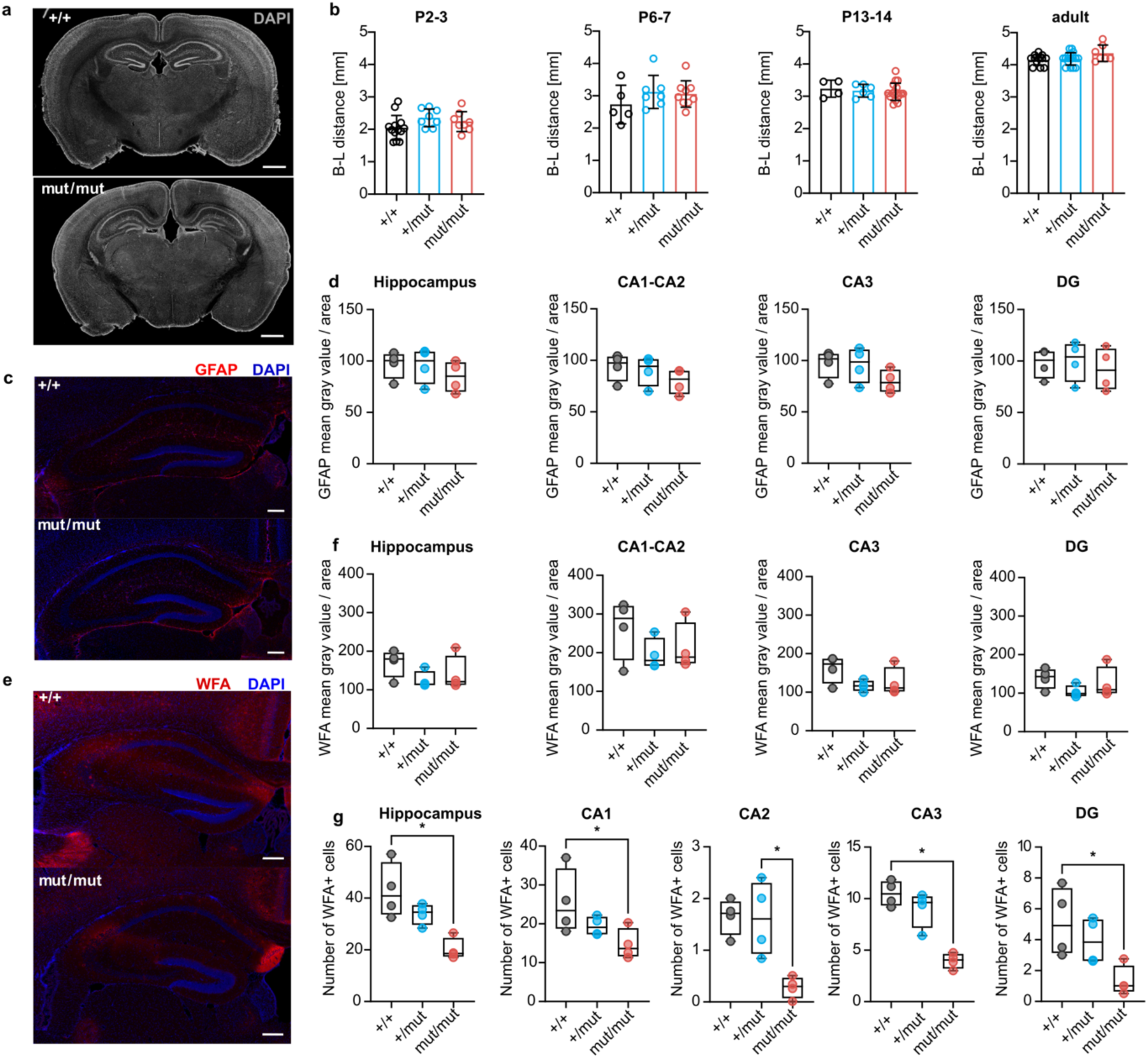
| No gross morphological changes in *CD1.Scn2a* p.A263V mouse brains at P21. **a-b**, Brain size comparison in *Scn2a* mice. **a**, Representative coronal brain slices of *Scn2a +/+* and *mut/mut* mice. **b**, Bregma-lambda distance at P2-3, P6-7, P13-14, and in adults, showing comparable brain size. Scale bar 1 mm. One-way ANOVA with Tukey’s post-hoc test (\**P* < 0.05, \*\**P* < 0.01, \*\*\**P* < 0.001) **c-d**, GFAP immunoreactivity in the hippocampus. **c**, Representative GFAP immunostaining of hippocampal coronal sections of *+/+* and *mut/mut* mice. Scale bar 200 µm. **d**, Quantification of GFAP staining intensity in the whole hippocampus, CA1-CA2, CA3, and dentate gyrus (DG). **e-g**, Perineuronal nets in the hippocampus. **e**, Representative WFA staining in hippocampal sections of *+/+* and *mut/mut* mice, scale bar 200 µm. **f**, Quantification of WFA staining intensity in the whole hippocampus, CA1-CA2, CA3, and DG. **g**, WFA-positive cell counts in the whole hippocampus, CA1, CA2, CA3, and DG of *+/+*, *+/mut*, and *mut/mut* mice. For all the experiments: *n =* 4 per genotype (*+/+*, *+/mut*, *mut/mut*). Unless indicated otherwise, Kruskal-Wallis test with Dunn’s correction for multiple comparisons was used for all analyses (\**P* < 0.05, \*\**P* < 0.01, \*\*\**P* < 0.001).

**Extended Data Fig. 2.**
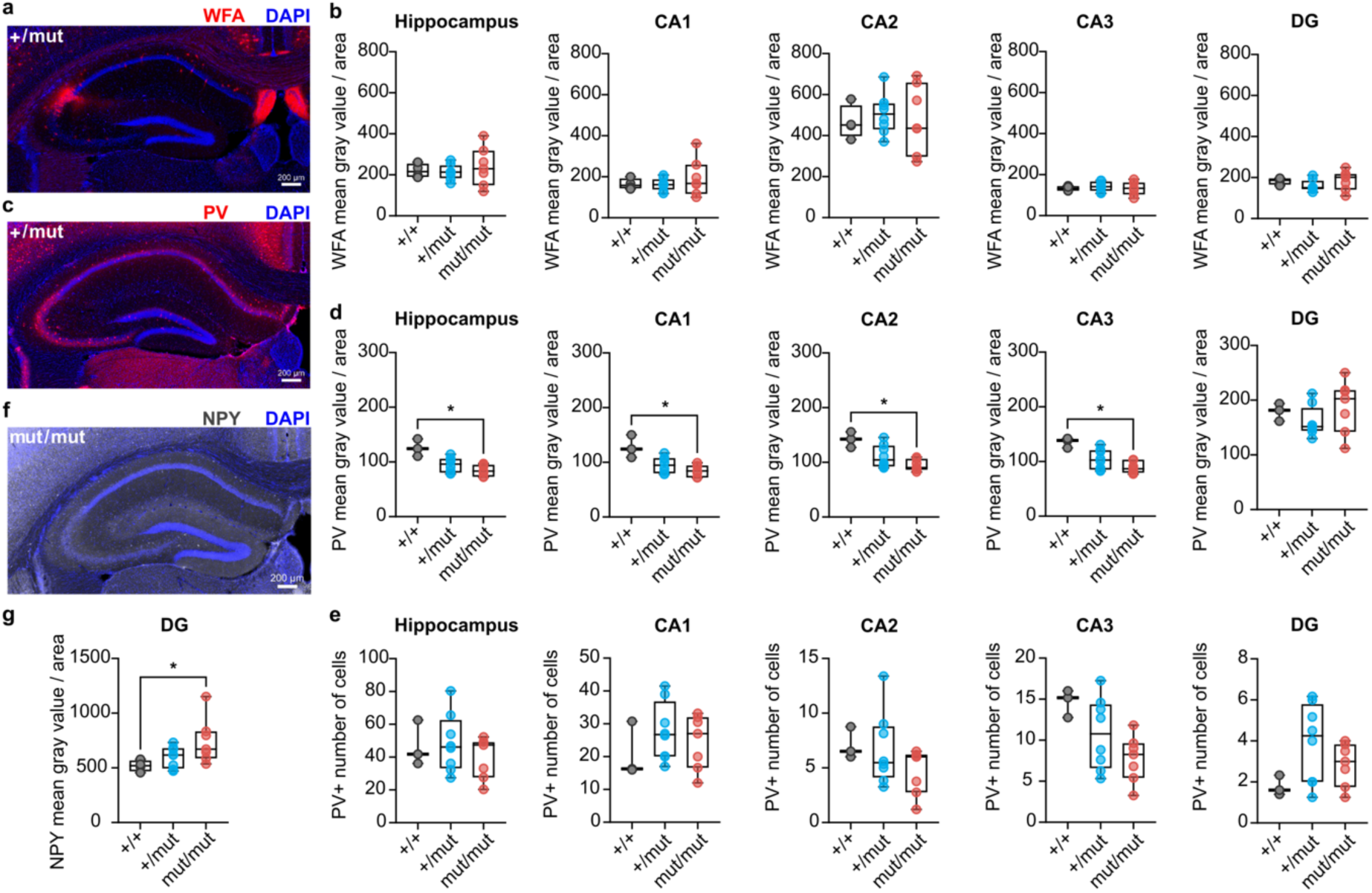
| No gross morphological changes in *CD1.Scn2a* p.A263V adult mouse brains. **a-b**, Perineuronal nets in the hippocampus. **a**, Representative WFA immunostaining in hippocampal coronal sections of a *+/mut* mouse. **b,** Quantification of WFA staining intensity in the whole hippocampus, CA1, CA2, CA3, and DG. **c-e**, PV hippocampal immunoreactivity. **c,** Representative PV staining in hippocampal sections of a *+/mut* mice. **d**, Quantification of PV staining intensity in the whole hippocampus, CA1, CA2, CA3, and DG. **e**, PV-positive cell counts in the whole hippocampus, CA1, CA2, CA3, and DG of *+/+*, *+/mut*, and *mut/mut* mice. **f-g**. NPY hippocampal immunoreactivity. **f,** Representative NPY staining in hippocampal sections of a *mut/mut* mouse. Scale bar 200 µm. **g**, Quantification of NPY hippocampal staining intensity. For all experiments: *n =* 19 (*+/+* = 4, *+/mut* = 8, *mut/mut* = 7), except for PV staining (*n =* 18 (*+/+* = 3, *+/mut* = 8, *mut/mut* = 7)); scale bar 200 µm. Kruskal-Wallis test with Dunn’s correction for multiple comparisons was used for all analyses (\**P* < 0.05, \*\**P* < 0.01, \*\*\**P* < 0.001).

### *Scn2a* p.A263V mutation leads to hyperactive behavior and strain-specific memory impairments

Epilepsy is frequently accompanied by a spectrum of cognitive and behavioral comorbidities. To characterize the behavioral phenotype of our *Scn2a* p.A263V mouse models, we conducted a series of tests assessing locomotor activity, motor skills, and various forms of memory in adult *B6.Scn2a* and *CD1.Scn2a* mice. In the open field test, *B6.Scn2a mut/mut* mice exhibited significant hyperactivity compared to their *+/+* and *+/mut* littermates as shown by the increased average distance moved per 5-minute time bin and a larger total distance moved for the entire test duration, particularly in males (Fig. 3a). *CD1.Scn2a* mice had a milder phenotype, with only *mut/mut* males displaying slightly increased total distance moved (Extended Data Fig. 3.1a). Both *B6.Scn2a* and *CD1.Scn2a* mice had largely normal motor function. Rotarod testing revealed no impairment in *Scn2a* p.A263V mutant mice (Extended Data Fig. 4a, c). CatWalk gait analysis showed subtle differences in *CD1.Scn2a* mice, that is, decreased average speed and base of support (BOS) of hind paws in *mut/mut* females compared to *+/mut*, and decreased BOS of front and hind paws in *mut/mut* males compared to *+/+* (Extended Data Fig. 4d). We next assessed various aspects of memory function. In the Y-maze test, *B6.Scn2a mut/mut* mice of both sexes showed impaired working memory, as indicated by a reduced percentage of alternation (Fig. 3b). In contrast, *CD1.Scn2a* mice displayed no significant differences in alternation, although *mut/mut* males transitioned less frequently between the arms (Extended Data Fig. 3.1b). In the contextual fear conditioning test, *B6.Scn2a mut/mut* and *+/mut* males displayed reduced freezing compared to *+/+* animals, indicating impaired fear memory (Fig. 3c). The contextual fear conditioning paradigm did not yield reliable results in *CD1.Scn2a* mice. Therefore, we, instead, used a passive avoidance test for this line. All *CD1.Scn2a* genotypes showed increased latency to enter the dark chamber during the test trial compared to the sample trial, suggesting successful acquisition of avoidance memory (Extended Data Fig. 3.1c). To assess spatial learning and memory, we performed one-trial spatial memory^26^ and water maze tests in *B6.Scn2a* mice. Male *B6.Scn2a mut/mut* mice had impaired performance, showing no preference for interacting with females and reduced exploration of the target area in both trials (Fig. 3d). Conversely, the performance of *mut/mut* and *+/mut* animals in the water maze was comparable to *+/+*, showing no significant impairment (Fig. 3e).

**Fig. 3.**
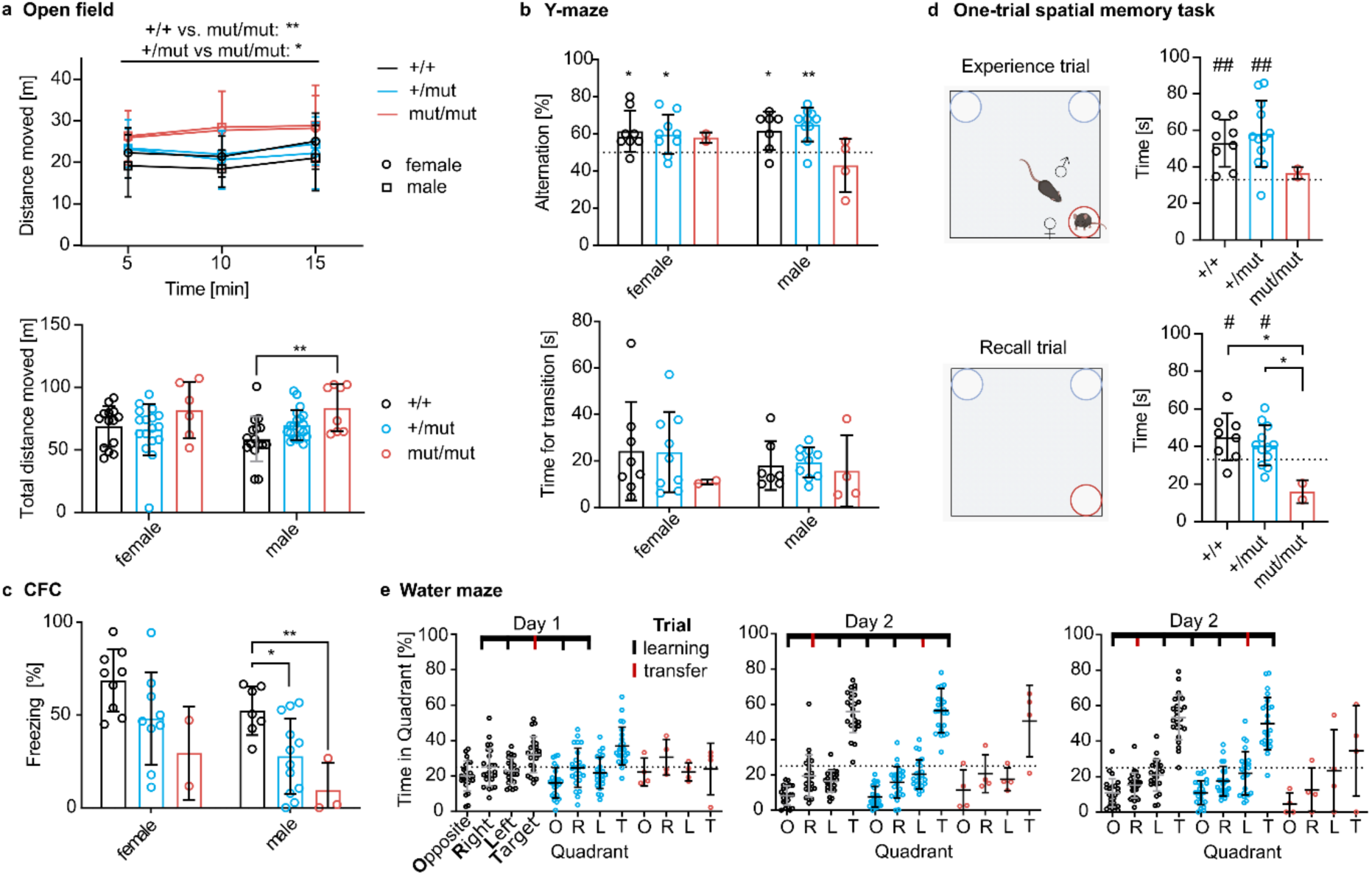
| Hyperactivity and impaired memory in adult *B6.Scn2a* p.A263V mice. **a**, Open field test. Top: Average distance moved per 5-minute time bin. Bottom: Total distance moved for the entire test duration (females, *n =* 37, *+/+* = 14, *+/mut* = 17, *mut/mut* = 6; males, *n =* 44, *+/+* = 16, *+/mut* = 20, *mut/mut* = 8). **b**, Y-maze test. Top: Percent of alternation (Wilcoxon-Signed-Rank Test, *P < 0.05, **P < 0.01). Bottom: Time for transition between the arms (females, *n =* 19, *+/+* = 8, *+/mut* = 9, *mut/mut* = 2; males, *n =* 21, +/+ = 7, *+/mut* = 10, *mut/mut* = 4). **c**, Contextual fear conditioning. Percentage of total freezing time (females, *n =* 20, *+/+* = 9, *+/mut* = 9, *mut/mut* = 2; males, *n =* 22, *+/+* = 7, *+/mut* = 12, *mut/mut* = 3). **d,** One-trial spatial memory task. Time spent interacting with females during the experience trial (top) and recall trial (bottom) compared to the chance level (males, *n =* 22, *+/+* = 8, *+/mut* = 12, *mut/mut* = 2; Wilcoxon-Signed-Rank Test, #P < 0.05) and between genotypes. **e**, Morris water maze. Time spent in each maze quadrant during the transfer trial, indicated in the scheme in red (*n =* 52, *+/+* = 24, *+/mut* = 24, *mut/mut* = 4, repeated measures ANOVA). Unless indicated otherwise, one-way ANOVA with Tukey’s post-hoc test was used for all analyses (\**P* < 0.05, \*\**P* < 0.01).

Taken together, our behavioral characterization revealed line-dependent effects of the *Scn2a* p.A263V mutation. *B6.Scn2a* mice, particularly *mut/mut* animals, were markedly hyperactive and had impaired working memory, spatial memory, and fear conditioning. By comparison, *CD1.Scn2a* mice showed milder hyperactivity and largely preserved cognitive function. While subtle gait differences were observed, motor skills, overall, remained intact in both lines.

**Extended Data Fig. 3.**
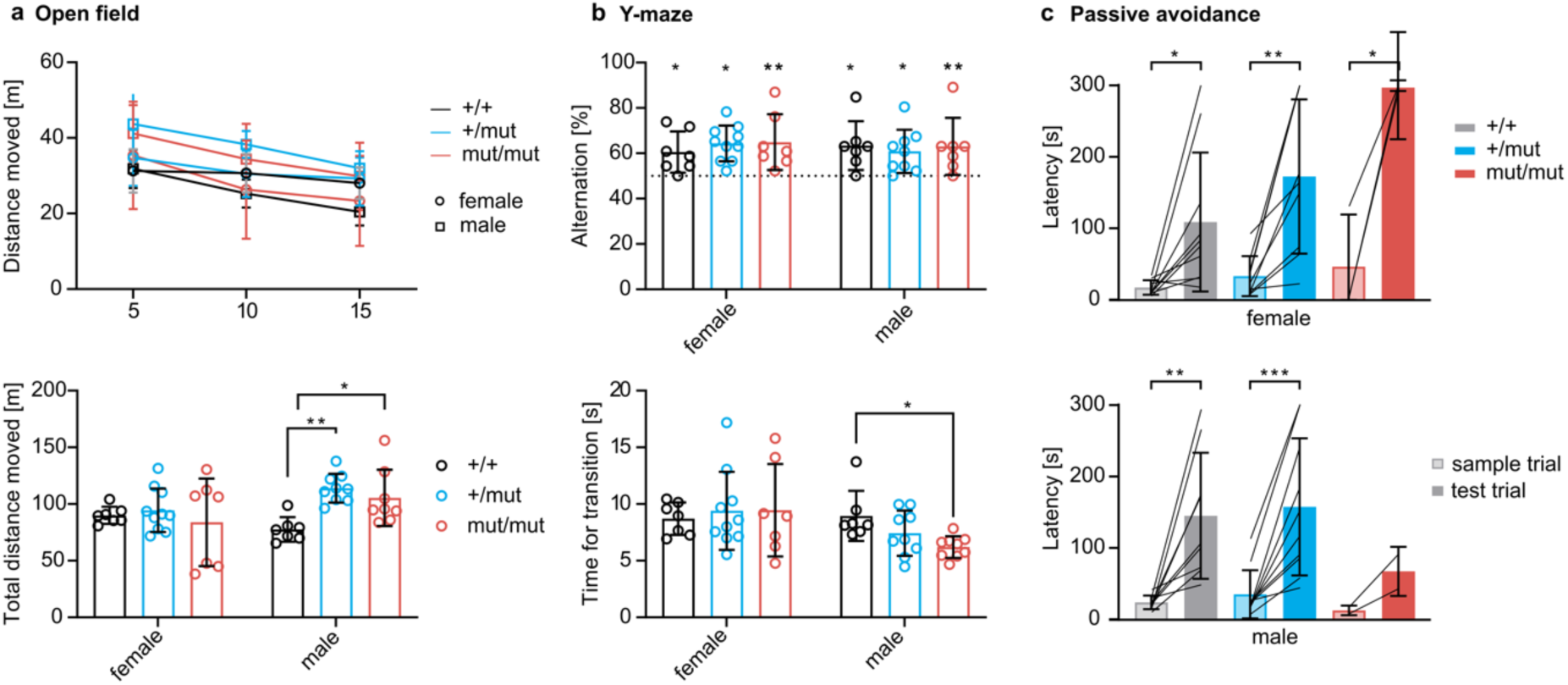
| Mild hyperactivity and unimpaired memory in adult CD1.Scn2a p.A263V mice. **a**, Open field test. Top: Average distance moved per 5-minute time bin. Bottom: Total distance moved for the entire test duration (females, *n =* 24, *+/+* = 7, *+/mut* = 10, *mut/mut* = 7; males, *n =* 24, *+/+* = 7, *+/mut* = 9, *mut/mut* = 8; one-way ANOVA with Tukey’s post-hoc test, *P < 0.05, **P < 0.01). **b**, Y-maze test. Top: Alternation percentage (Wilcoxon-Signed-Rank Test, *P < 0.05, **P < 0.01). Bottom: Time for transition between the arms (females, *n =* 24, *+/+* = 7, *+/mut* = 10, *mut/mut* = 7; males, *n =* 24, *+/+* = 7, *+/mut* = 9, *mut/mut* = 8; one-way ANOVA with Tukey’s post-hoc test, *P < 0.05). **c**, Passive avoidance test. Latency to enter the dark chamber during the test trial compared to the sample trial for females (top, *n =* 24, *+/+* = 10, *+/mut* = 10, *mut/mut* = 3) and males (bottom, *n =* 24, *+/+* = 9, *+/mut* = 9, *mut/mut* = 2; paired t-test, \**P* < 0.05, \*\**P* < 0.01, \*\*\**P* < 0.001).

**Extended data Fig. 4.**
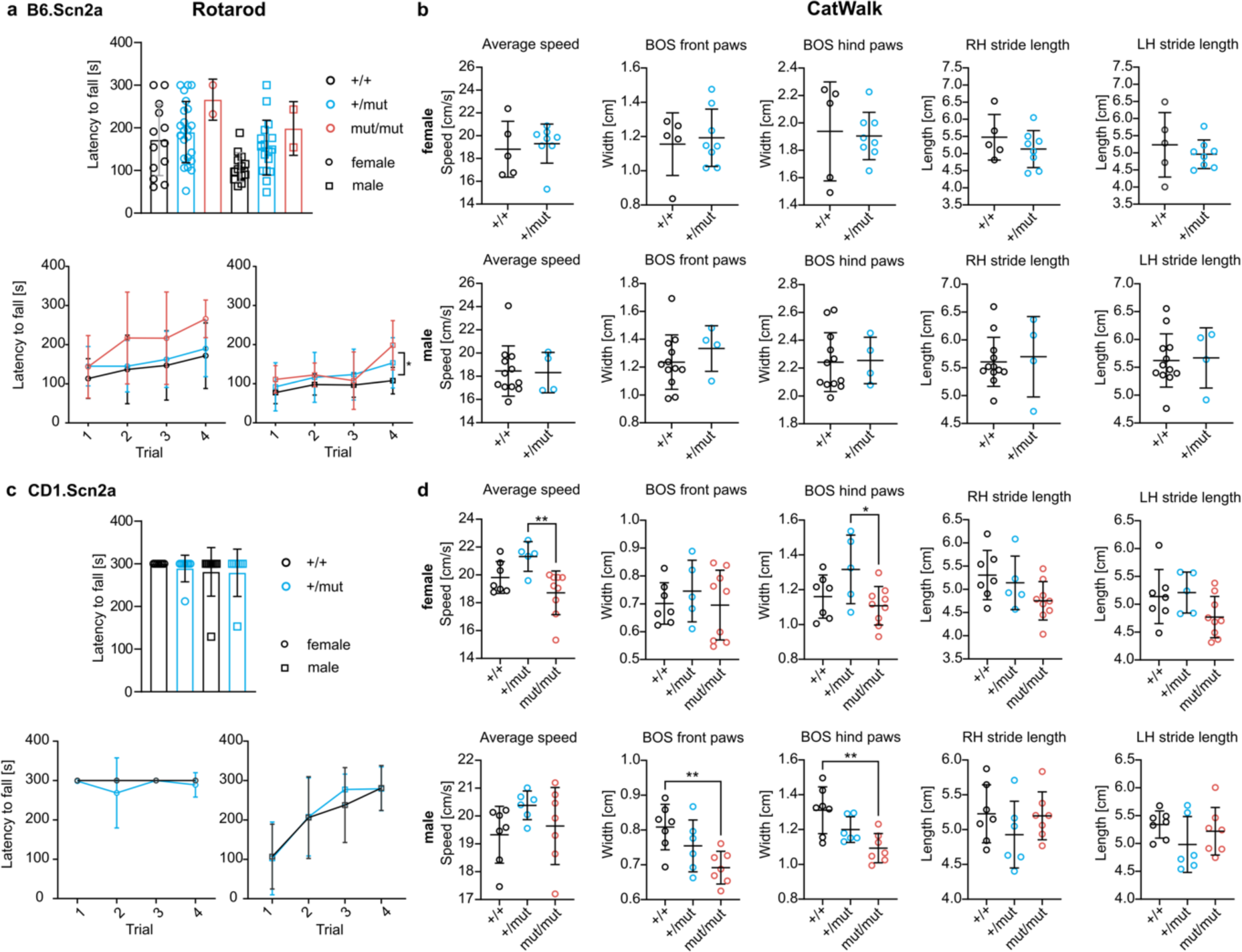
| Normal locomotor behavior in adult *B6.Scn2a* and *CD1.Scn2a* p.A263V mice. **a**, Rotarod test in *B6.Scn2a* mice. Top: Latency to fall across all trials. Bottom: Latency to fall per trial for females (right, *n =* 38, *+/+* = 13, *+/mut* = 23, *mut/mut* = 2) and males (left, *n =* 32, *+/+* = 13, *+/mut* = 17, *mut/mut* = 2). One-way ANOVA with Tukey’s post-hoc test, *P < 0.05. **b**, CatWalk gait analysis in *B6.Scn2a* mice. Left to right: Average speed, base of support (BOS) of front paws, BOS of hind paws, stride length for right hind (RH) paw, stride length for left hind (LH) paw in females (top, *n =* 13, *+/+* = 5, *+/mut* = 8) and males (bottom, *n =* 16, *+/+* = 12, *+/mut* = 4). Unpaired t-test. **c**, Rotarod test in CD1.Scn2a mice. Top: Latency to fall across all trials. Bottom: Latency to fall per trial for females (right, *n =* 15, *+/+* = 7, *+/mut* = 8) and males (left, *n =* 16, *+/+* = 9, *+/mut* = 7). Unpaired t-test. **d**, CatWalk gait analysis in *CD1.Scn2a*. Left to right: Average speed, base of support (BOS) of front paws, BOS of hind paws, stride length for right hind (RH) paw, stride length for left hind (LH) paw in females (top, *n =* 21, *+/+* = 7, *+/mut* = 5, *mut/mut* = 9) and males (bottom, *n =* 20, *+/+* = 7, *+/mut* = 6, *mut/mut* = 7). One-way ANOVA with Tukey’s post-hoc test, \*\**P* < 0.01.

### Cortical and hippocampal interictal activities show persistent changes in *Scn2a* p.A263V mice

Given the importance of interictal activity as a correlate of impaired cognitive performance^27^, we next analyzed cortical network activity across different sleep/wake states in the adult ECoG recordings (Extended Data Fig. 5). Changes were more pronounced in *mut/mut* animals, with increased power in the theta frequency range (5-9 Hz) during slow-wave sleep (SWS) and decreased theta oscillation frequency during wakefulness. Moreover, mid-gamma (50-90 Hz) power during wakefulness was increased in both *mut/mut* and *+/mut* mice, showing a gene-dose-dependent effect. Motivated by our previous observations of subtle changes in hippocampal interneuron immunoreactivity and alterations in hippocampal-dependent behaviors, we conducted an in-depth analysis of hippocampal network activity. Using in vivo silicon probe local field potential (LFP) recordings, we found layer-specific alterations of hippocampal network activity in adult *CD1.Scn2a* animals (Fig. 4a-d). Mid-gamma frequency showed a consistent decrease of 5.4 Hz [4.5 6.0] Hz on average across all layers in *mut/mut* mice (Fig. 4e). In *+/mut* mice, both mid-gamma frequency and power were specifically decreased in layers containing or adjacent to cell bodies, namely stratum pyramidale, stratum radiatum, and granule cell layer (Fig. 4d, e). The modulation index for mid-gamma oscillation amplitude by theta phase showed a trend toward a decrease in the middle molecular layer of the dentate gyrus in *mut/mut* mice, accompanied by an increased modulation index for high-gamma oscillations (Fig. 4f, g). Of particular note, the hippocampal layers with the most pronounced changes corresponded to areas receiving input from the medial entorhinal cortex (MEC), which is also a pivotal contributor to mid-gamma oscillations^28,29^. Furthermore, the theta phase shift of the CA1 LFP depth profile^30^, a measure of the relative contribution of afferent CA3 and MEC theta activity, was higher in *mut/mut* animals (Fig. 4h, i). We also examined sharp-wave ripple (SWRs) oscillations, which play a crucial role in memory consolidation and replay of hippocampal-dependent memories^31^. Sharp-wave ripple peak frequencies trended toward a decrease in *mut/mut* and *+/mut* mice. However, there were no differences in ripple incidence or power between genotypes (Fig. 4j, k). The median incidence observed during the recordings was 0.1 [0.1 0.2] Hz.

**Fig. 4.**
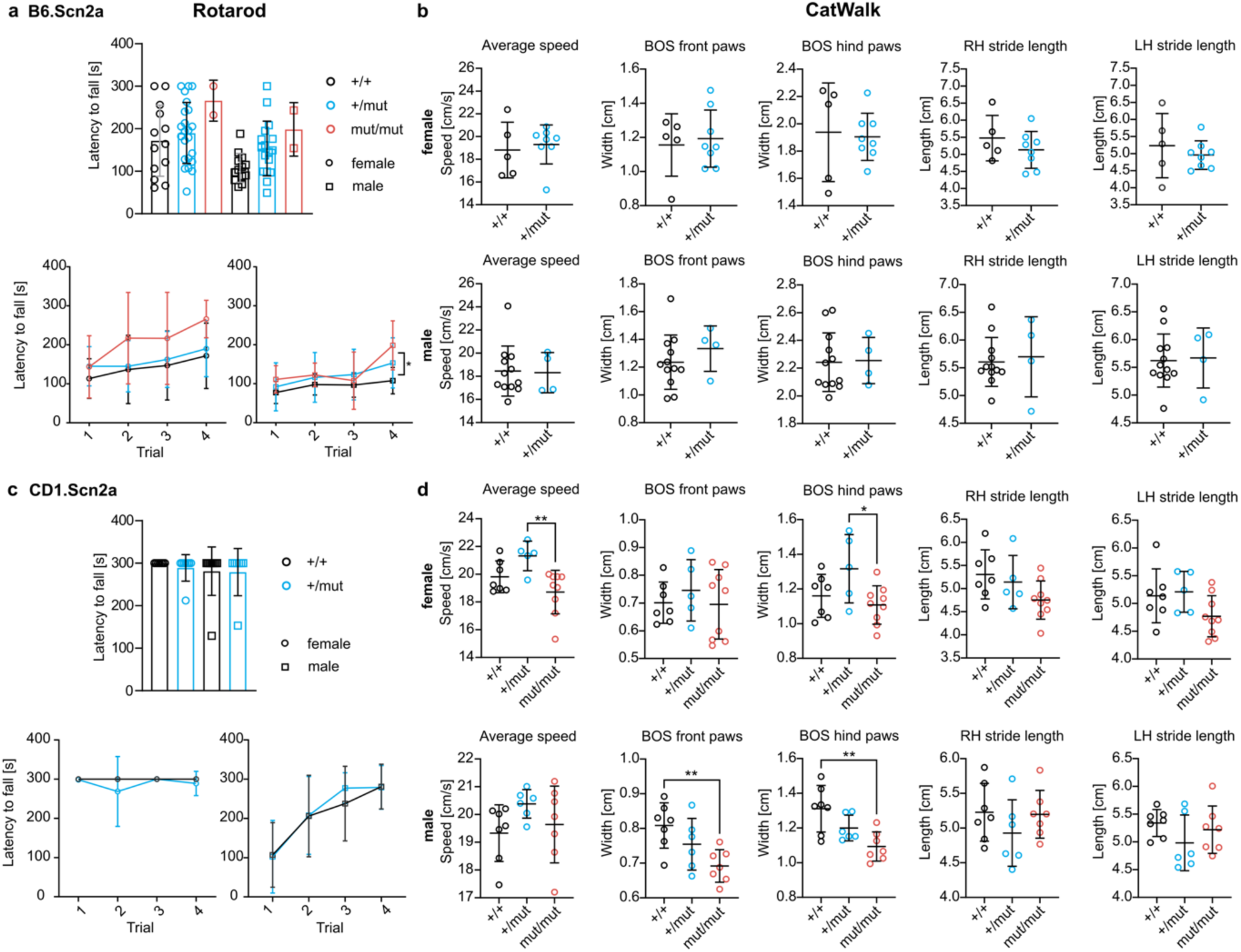
| Altered hippocampal in vivo network activity in adult *CD1.Scn2a* p.A263V mice. **a**, Silicon probe track in the dorsal hippocampus (coronal hippocampal section, cell nuclei stained with DAPI, DiI staining for the probe track, scale bar 500 µm). White box: Magnified inset with hippocampal layers marked (SO – stratum oriens, SP – stratum pyramidale, SR – stratum radiatum, SLM – stratum lacunosum moleculare, OM – outer molecular layer of the dentate gyrus (DG), MM – middle molecular layer of the DG, GL – granular cell layer of the DG, HL – hilus). The inset on the left depicts a head-fixed setup with a mouse freely moving on a floating platform. **b**, Representative local field potential (LFP) depth profile from the CA1-DG axis during the running period. **c**, Representative band-limited power (BLP) plots for theta (5-9 Hz), mid-gamma (50-90 Hz), and high-gamma (90-130 Hz) frequency bands for a *mut/mut* animal. Faint lines indicate BLP for separate running epochs, and bold lines represent the mean of all epochs. **d**, BLP analysis of theta (left) and mid-gamma (right) frequency bands. Faint lines indicate the mean of all running epochs for each animal, and bold lines indicate the mean of each genotype. **e**, Background-subtracted mid-gamma peak frequency (left) and normalized power (right) across hippocampal layers. **f**, Theta modulation index of mid-gamma amplitude. Left: Across all layers. Faint lines indicate modulation for each animal, and bold lines represent the mean for each genotype. Right: Mean modulation index of mid-gamma oscillations in the dentate gyrus middle molecular layer. **g**, Theta modulation index of high gamma amplitude. Left: Across all layers. Faint lines indicate modulation for each animal, and bold lines show the mean for each genotype. Right: Mean modulation index of high gamma oscillations in the dentate gyrus middle molecular layer. **h**, Theta phase shift. Faint lines indicate a theta phase shift for each animal, and bold lines show the mean for each genotype. **i**, Mean theta phase shift in stratum lacunosum moleculare (left) and middle molecular layer (left). **j**, Probability density function plot of sharp-wave ripple (SWR) peak frequency. **k**, SWR peak power (left) and frequency (right) for each animal. Sample size: *n =* 25 (*+/+* = 9, *+/mut* = 7 [5 for theta phase shift and modulation index], *mut/mut* = 9). Kruskal-Wallis test with Dunn’s correction for multiple comparisons was used for all analyses (\**P* < 0.05, \*\**P* < 0.01, \*\*\**P* < 0.001).

The observed changes in mid-gamma oscillations, theta-gamma coupling, and sharp-wave ripples in the hippocampus may affect information processing and memory formation. These oscillatory patterns are crucial for precisely timing neuronal firing and activity coordination across brain regions^29^. Our findings demonstrate remarkably subtle but persistent cortical and hippocampal network activity changes in *Scn2a* p.A263V mice given the severe seizure phenotype. The alterations were more pronounced in *mut/mut* mice, aligning with a potential gene-dose effect and seizures observed in adult *mut/mut* CD1.Scn2a animals.

**Extended Data Fig. 5.**
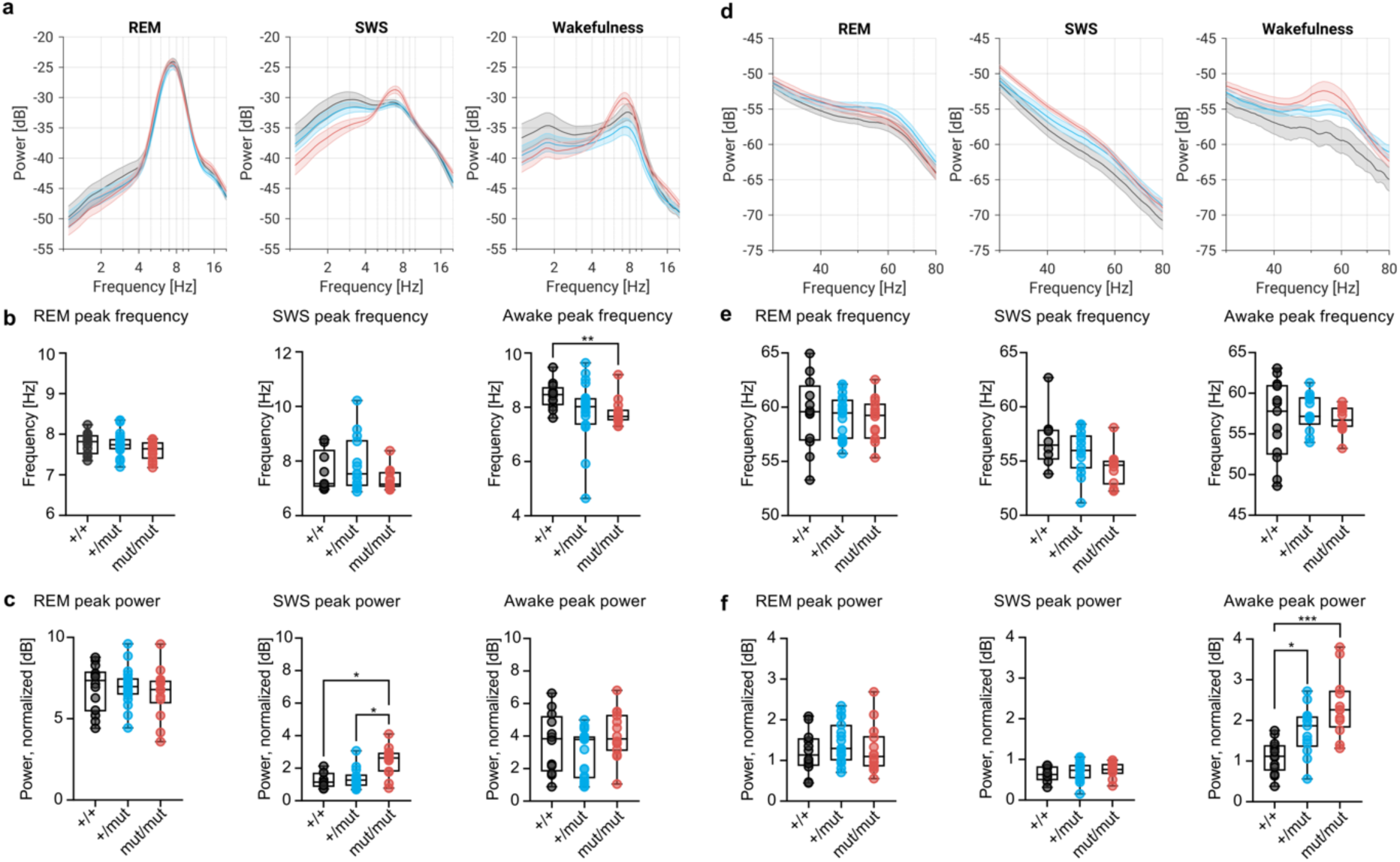
| Altered cortical interictal activity in adult *B6.Scn2a* and *CD1.Scn2a* p.A263V mice. For all panels, data are presented for rapid eye movement (REM) sleep (left), slow-wave sleep (SWS) (middle), and wakefulness (right) epochs of the electrocorticogram (ECoG) recording. **a**, Power spectral density (PSD) plots illustrating theta frequency peaks for different genotypes. **b**, Peak theta frequency. **c**, Peak theta power. **d**, PSD plots illustrating mid-gamma frequency peaks for different genotypes. **e**, Peak mid-gamma frequency. **f**, Peak mid-gamma power. PSD data were z-score normalized within each sleep/wake state, and peak frequencies and powers were calculated from background-subtracted PSD data. Data was pooled for *B6.Scn2a* (*n =* 22, *+/+* = 7, *+/mut* = 11, *mut/mut* = 2) and *CD1.Scn2a* (*n =* 28, *+/+* = 7, *+/mut* = 9, *mut/mut* = 12) animals. Kruskal-Wallis with Dunn’s correction for multiple comparisons was used for all analyses (\**P* < 0.05, \*\**P* < 0.01, \*\*\**P* < 0.001).

### *Scn2a* p.A263V variant causes seizures and altered hippocampal network activity in neonates

Given that the *Scn2a* p.A263V variant is associated with early-onset DEEs in patients, we next investigated its effects during early postnatal development, when Na_v_1.2 channels are crucial for action potential generation^32^. We first performed c-Fos immunostaining in P7 *B6.Scn2a* animals (Fig. 5a) to assess the overall neuronal activity in neonatal mice. Quantification revealed a significant increase in c-Fos-positive cells in all hippocampal regions of both *+/mut* and *mut/mut* animals compared to wildtype counterparts (Fig. 5b). In the MEC of *mut/mut* animals, deep layers (IV-VI) showed more c-Fos-positive cells than superficial ones (I-III), suggesting that network hyperexcitability might originate in the hippocampus at this age because deep MEC layers receive the hippocampal output.

**Fig. 5.**
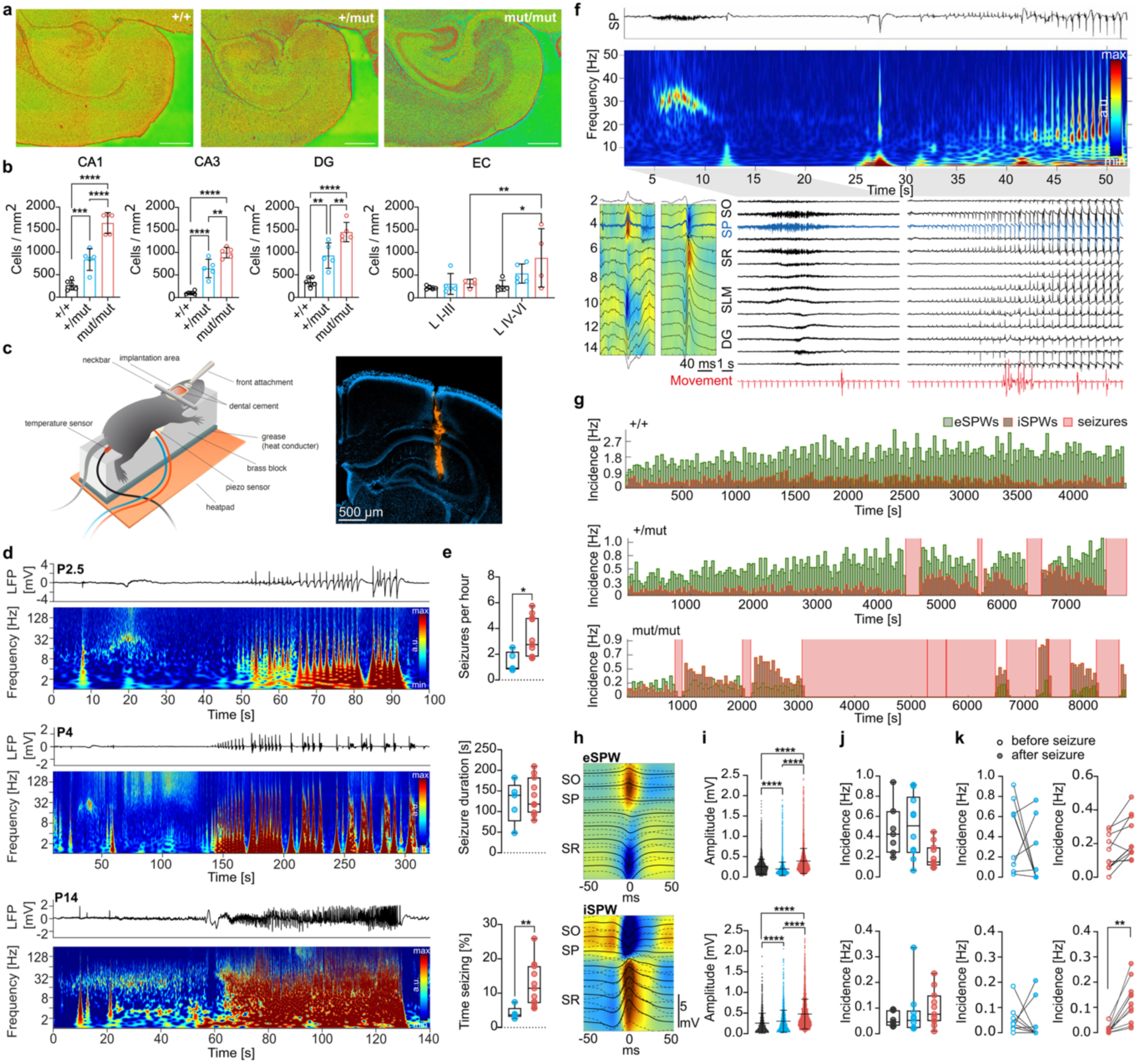
| Seizures and altered network activity in *+/mut* and *mut/mut B6.Scn2a* neonates. **a,** Representative images of c-Fos staining in hippocampal sagittal sections from P7 *B6.Scn2a +/+* (left), *+/mut* (middle), and *mut/mut* (right) animals. Scale bar 500 µm. **b,** c-Fos-positive cell counts in hippocampal CA1, CA3, DG (*n =* 16, *+/+* = 6, *+/mut* = 5, *mut/mut* = 5; one-way ANOVA with Tukey’s post-hoc test, \**P* < 0.05, \*\**P* < 0.01, \*\*\**P* < 0.001, \*\*\*\**P* < 0.0001) and superficial (L I-III) and deep (L IV-VI) layers of the entorhinal cortex (*n =* 14, *+/+* = 5, *+/mut* = 5, *mut/mut* = 4; two-way ANOVA with Šídák’s multiple comparisons test, \**P* < 0.05, \*\**P* < 0.01). **c, Left**: Schematic setup representation for acute in vivo local field potential (LFP) recordings. **Right**: Silicon probe track in dorsal hippocampus of *mut/mut* animal aged P6 (coronal hippocampal section, cell nuclei stained with DAPI, DiI staining for the probe track, scale bar 500 µm). **d,** Representative spontaneous seizures (raw LFP trace of a channel in stratum radiatum and respective spectrogram of a seizure episode) in a *B6.Scn2a mut/mut* mice aged P2.5 (top), P4 (middle), and P14 (bottom). **e,** Seizure characteristics in *B6.Scn2a +/mut* (blue) and *mut/mut* (red) mice aged P6-P7 (*n =* 21, *+/mut* = 10 (5 with seizures), *mut/mut* = 11; Mann-Whitney U test, \**P* < 0.05). Top: Average number of seizures per hour. Middle: Mean duration of a seizure episode. Bottom: Percentage of recording time during which an animal had seizures. **f,** Hippocampal network activity and seizure in *mut/mut* mouse aged P7. Top: Raw LFP trace of a channel in stratum pyramidale (SP) and the respective spectrogram. Magnified areas below are indicated with gray boxes. Bottom: Network activity across hippocampal layers (SO – stratum oriens, SR -stratum radiatum, SLM - stratum lacunosum moleculare, DG -dentate gyrus), including current source density (CSD) profile (current sinks in blue, current sources in red) with overlaid LFP traces of a representative early sharp wave (eSPW) and an inverted sharp wave (iSPW) (left), LFP traces of low gamma oscillations preceding the seizure (middle) and ictal activity (right). Red traces indicate animal movement. **g,** Example of incidence of eSPWs (green) and iSPWs (orange) throughout the recording for *+/+* (top), *+/mut* (middle), and *mut/mut* (bottom) animals aged P6-P7. Seizure periods are shaded in pink. **h,** Mean CSD profile of eSPWs (top) and iSPWs (bottom) with overlaid LFP traces for *B6.Scn2a* mice aged P6-P7. **i,** Amplitude of all detected SPWs (top) and iSPWs (bottom) per genotype. One-way ANOVA with Tukey’s post-hoc test, ****P < 0.0001. **j,** Mean incidence of SPWs (top) and iSPWs (bottom) (*n =* 28, *+/+* = 7, *+/mut* = 10, *mut/mut* = 11; Kruskal-Wallis test with Dunn’s correction for multiple comparisons). **k,** Incidence of SPWs (top) and iSPWs (bottom) before and after seizure for *+/mut* (left) and *mut/mut* (right) animals (*n =* 21, *+/mut* = 10, *mut/mut* = 11; Two-way ANOVA with Šídák’s multiple comparisons test, \*\**P* < 0.01).

Accordingly, acute hippocampal in vivo LFP recordings in locally anesthetized head-fixed neonatal *B6.Scn2a* mice (P2.5-P14) revealed spontaneous seizures in *mut/mut* animals as early as P2.5, evident in both genotypes by P6 (Fig. 5c, d). At P6-P7, 100% of *mut/mut* and 50% of *+/mut* mice had seizures during the 1-2h-long recording periods. Seizure frequency was higher for *mut/mut* mice (2.7 [1.8 4.8] seizures/hour) than for *+/mut* mice (0.9 [0.8 2.2] seizures/hour). Moreover, the total seizing time was increased in *mut/mut* animals, reaching 26% of the recording time. The seizure duration of approximately two minutes was similar for both genotypes (Fig. 5e). Between the seizures, we observed network activity typical of the developing murine hippocampus. However, we also noted the emergence of unusual, several second-long low gamma oscillations (25-35 Hz) stereotypically preceding seizures already at P2.5 (Fig. 5f,d, Extended Data Fig. 6a,b). The CSD depth profile indicated a CA3 origin of these gamma oscillations as the amplitudes were maximal in stratum oriens and stratum radiatum (Fig. 5f). Detailed examination of network activity revealed alterations in early hippocampal sharp waves (eSPWs) (Fig. 5g,h). In neonates, eSPWs are well-described events crucial to hippocampal circuit maturation^33–35^. In addition to regular eSPWs, we also detected inverted sharp waves (iSPWs). These events are characterized by an *inverted* sink and source pattern in the CSD profile, featuring a prominent sink in stratum oriens, which indicates contralateral CA3 may drive them^36^. To investigate the origin of iSPWs, we performed high-frequency stimulation of the ventral hippocampal commissure (VHC), which carries association fibers from contralateral CA3 (Extended Data Fig. 6g-i). CSD analysis of the post-HFS evoked response revealed a prominent sink in stratum oriens, matching the pattern observed during spontaneous iSPWs. In *+/+* animals aged P5-P7 (n = 4), the iSPWs/eSPWs incidence ratio increased from 0.02 [0.004 0.27] to 0.48 [0.15 0.88] in the 30 minutes following VHC stimulation, supporting our hypothesis that iSPWs are driven by contralateral CA3 inputs. Both eSPW and iSPW amplitudes were significantly increased in *mut/mut* and *+/mut* compared to *+/+* mice (Fig. 5h, i). Of special note here, while the overall incidence of eSPWs and iSPWs was similar across genotypes throughout the recording, we observed a marked increase in iSPW occurrence following seizure onset in *mut/mut* mice (Fig. 5j, k). This postictal increase in iSPWs implicates the CA3 region as a potential seizure pacemaker.

To assess whether these findings were line-specific, we performed similar recordings in *CD1.Scn2a* neonates. As in *B6.Scn2a* mutants, we observed gamma oscillations preceding seizures (Extended Data Fig. 6a,b). While the overall pattern of results was comparable, with seizures observed in *+/mut* and *mut/mut* animals, we noted some line-specific differences (Extended Data Fig. 6a-f). At P6-7 the seizure phenotype was more severe in CD1 mice; that is, 100% of *mut/mut* and 69% of *+/mut* mice had seizures, with a median frequency of 4.5 [2.7 8.3] seizures/hour for *mut/mut* and 6.5 [3.0 15.6] seizures/hour for *+/mut* animals. Seizures lasted for about seven minutes on average in *mut/mut* animals compared to two minutes in *+/mut*, while *mut/mut* mice had seizures up to 78% of the recording time. Conversely, interictal network activity seemed less affected in *CD1.Scn2a* mice. No difference was observed in iSPW occurrence before and after seizures, although the amplitude differences were preserved.

As sodium channel blockers, including phenytoin, are effective in the treatment of patients with GoF variants, including in patients with *SCN2A* p.A263V, we tested whether the hippocampal seizures present in every homozygous mutant could be attenuated by daily postnatal phenytoin treatment from P1. When recorded at P6-P9 one day following the last treatment (i.e., at the phenytoin trough serum levels of 13±6 mg/dl; n = 4 (pooled samples from 5 neonates)), no phenytoin-treated *mut/mut* neonate showed hippocampal seizures (n = 5, *P* = 0.0079 (Fisher’s exact test)). In contrast, all vehicle-treated *mut/mut* neonates (n = 4) had seizures during the 90 to 120-minute recording periods.

Taken together, these results demonstrate that the *Scn2a* p.A263V mutation leads to profound alterations in spontaneous in vivo neonatal hippocampal network activity and the emergence of unusual gamma oscillations and seizures during early postnatal development, which likely originate in the hippocampal CA3 region and can be prevented by phenytoin.

**Extended Data Fig. 6.**
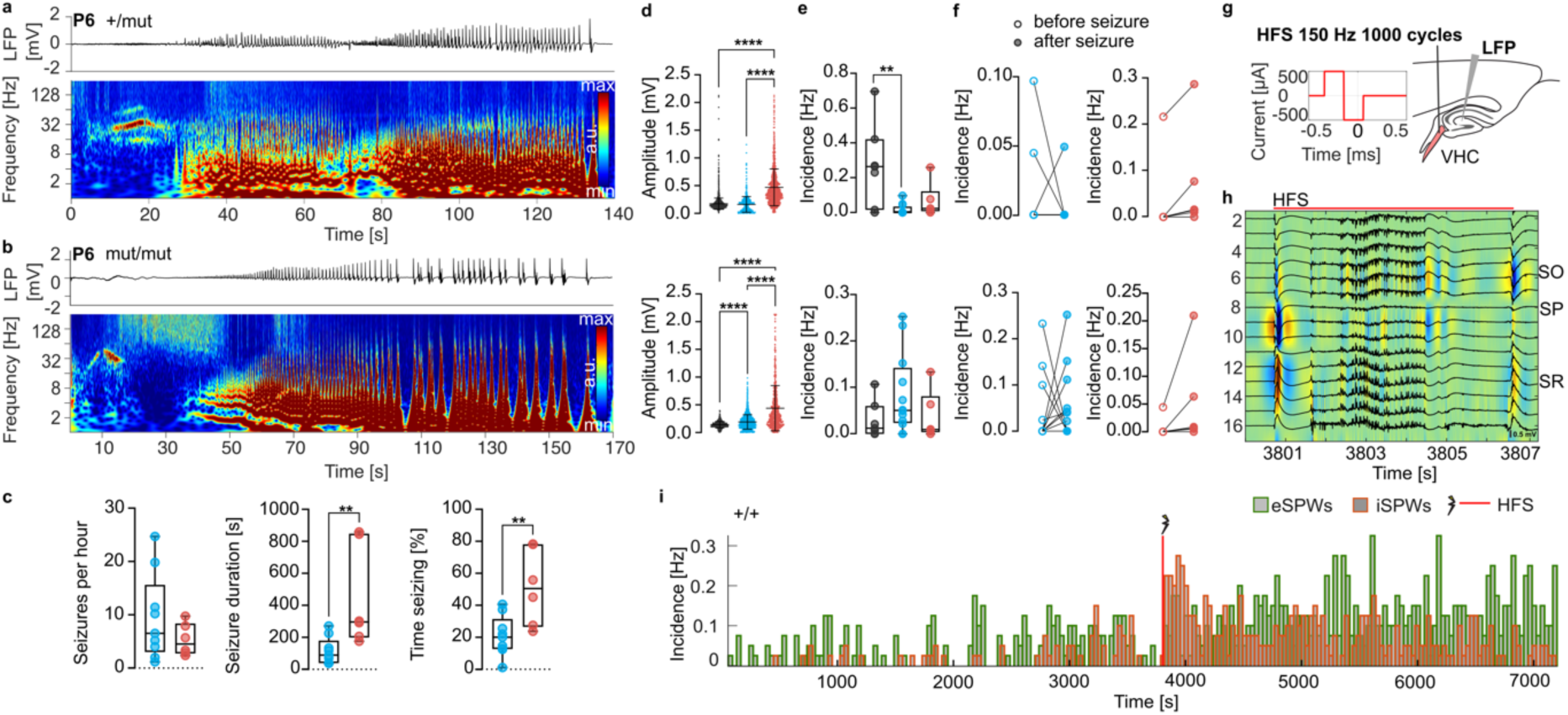
| Seizures and altered network activity in *+/mut* and *mut/mut CD1.Scn2a* neonates. **a,** Representative spontaneous seizure (raw LFP trace of a channel in stratum radiatum and respective spectrogram of a seizure episode) in *CD1.Scn2a +/mut* mouse aged P6. **b,** Representative spontaneous seizure (raw LFP trace of a channel in stratum radiatum and respective spectrogram of a seizure episode) in *CD1.Scn2a mut/mut* mouse aged P6. **c,** Seizure characteristics in *CD1.Scn2a +/mut* (blue) and *mut/mut* (red) mice aged P6-P9 (*n =* 19, *+/mut* = 13 (9 with seizures), *mut/mut* = 6; Mann-Whitney U test, *P < 0.05). Left: Average number of seizures per hour. Middle: Mean duration of a seizure episode. Right: Percentage of recording time during which the animal had seizures. **d,** Amplitude of all detected eSPWs (top) and iSPWs (bottom) per genotype. One-way ANOVA with Tukey’s post-hoc test, ****P < 0.0001. **e,** Mean incidence of eSPWs (top) and iSPWs (bottom) (*n =* 25, *+/+* = 7, *+/mut* = 12, *mut/mut* = 6; Kruskal-Wallis test with Dunn’s correction for multiple comparisons). **f,** Incidence of eSPWs (top) and iSPWs (bottom) before and after seizure for *+/mut* (left) and *mut/mut* (right) animals (*n =* 17, *+/mut* = 11, *mut/mut* = 6; Two-way ANOVA with Šídák’s multiple comparisons test). **g,** Schematic diagram showing the high-frequency stimulation (HFS) (1000 cycles of 150 Hz, 500 µA, 0.2 ms biphasic stimulus) applied to the ventral hippocampal commissure (VHC). **h,** CSD profile with overlaid LFP traces from the hippocampus during VHC stimulation showing a prominent sink in stratum oriens. **i,** Incidence of eSPWs (green) and iSPWs (orange) one hour before and one hour after VHC stimulation (vertical dotted line) in a P5 +/+ mouse.

### The hippocampus acts as a pacemaker for ongoing early network activity in *Scn2a* p.A263V mice

To further investigate the early postnatal mechanisms of epileptogenesis in the *B6.Scn2a* model, we quantified the ongoing network activity in the cortico-hippocampal formation of *+/mut* and *+/+* mice. We focused on P3, the earliest age at which we reliably detected hippocampal seizures in vivo. Our previous data showed that in WT mice, this activity reflects early network oscillations (ENOs), whose frequency and pattern in vivo and in situ are very similar at P3^37^. Therefore, we conducted large-scale Ca^2+^ imaging in acute slices labeled with Oregon Green BAPTA-1 AM (OGB-1 AM), which facilitated access to the hippocampus while maintaining network dynamics comparable to the in vivo state (Fig. 6a). The ENO-induced fluorescence changes were extracted from raw videos using principal component analysis (see Materials and methods). Spatial analyses of ENO-induced Ca^2+^ signals at high temporal (50 frames/s) resolution showed that each Ca^2+^ signal represented a wave of activity originating in a specific cortical or hippocampal area (pacemaker region) and spreading throughout the tissue (Fig. 6a, b). Notably, the distribution of pacemaker regions differed significantly between the genotypes (Yates chi-square, P=0.01). In *+/+* mice, 62.6% of waves originated in the cortex, whereas 56.6% of waves were of hippocampal origin in *+/mut* mice. Higher pacemaking activity in *+/mut* mice was observed across all hippocampal areas, except the subiculum. The difference between *+/+* and *+/mut* mice was especially prominent in CA3 (Fig. 6c; P=0.048, Wilcoxon Rank test), but the significance was lost after correcting for multiple comparisons (Fig. 6d).

**Fig. 6.**
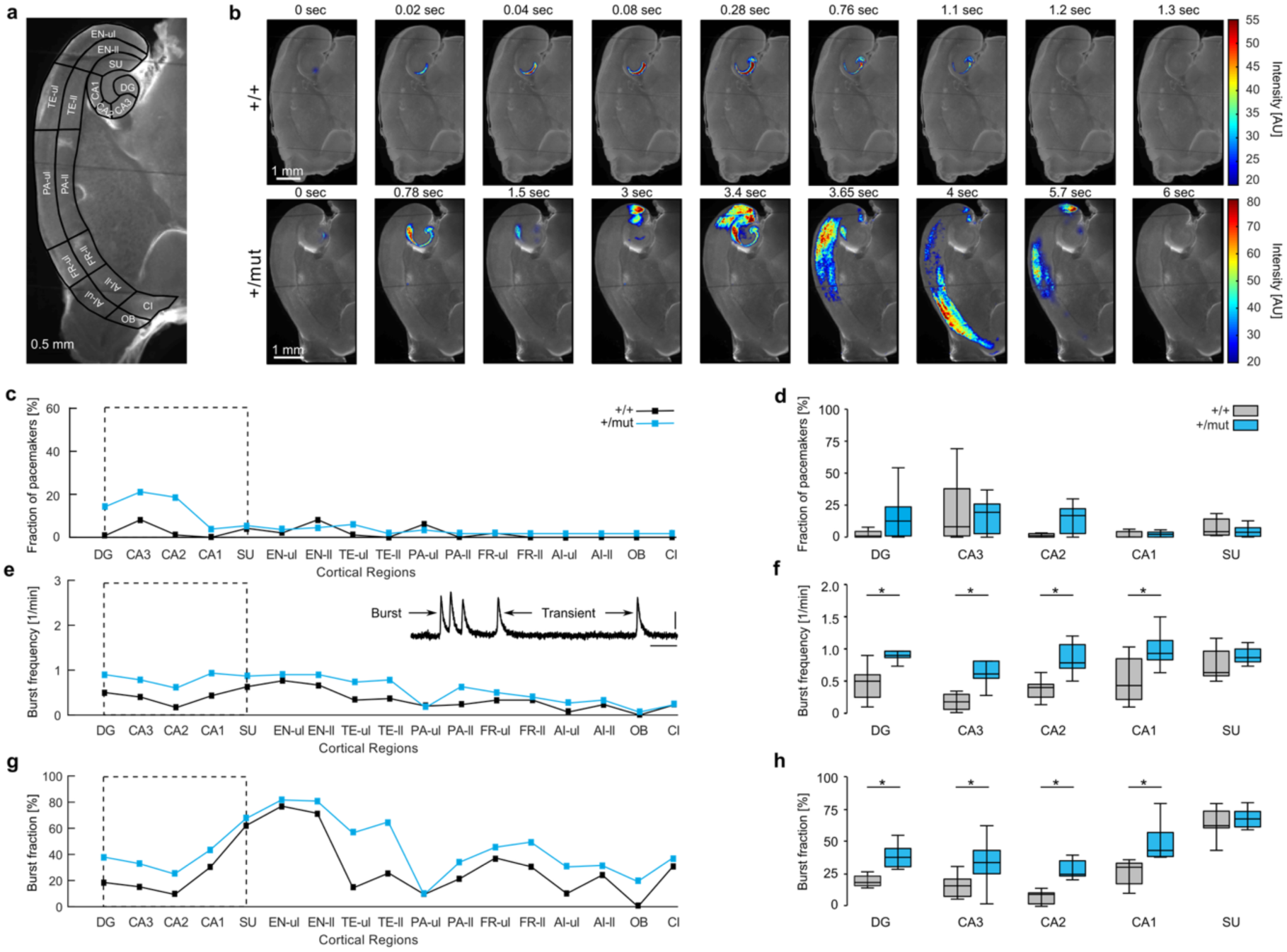
| Altered ongoing spontaneous activity in the cortico-hippocampal formation of the neonatal *B6.Scn2a +/mut* mice. **a,** A 500-µm-thick horizontal slice of the P3 mouse brain stained with the membrane-permeant fluorescent Ca^2+^ indicator Oregon Green™ 488 BAPTA-1 AM (OGB-1). Dash lines delineate different brain areas (DG: Hippocampal dentate gyrus; CA1; CA2, CA3: Areas 1, 2 and 3 of the Ammon’s horn; SU: Subiculum; EN: Entorhinal, TE: Temporal, PA: Parietal, FR: Frontal, AI: Agranular insular, OB: Orbital and CI: Cingulate cortices; ul stands for upper and ll for lower cortical layers). **b,** Representative waves of activity visualized as increases in the fluorescence intensity of OGB-1 (AU: Arbitrary units) in *+/+* (upper row) and *+/mut* (lower row) mice. **c,** Median (across mice) values illustrating the fractions of wave pacemakers detected (as described in Materials and methods) in different brain areas mentioned above (here and below n= 7 *+/+* and 9 *+/mut* mice). The data for the hippocampal regions (framed part of the graph) are detailed in d. **d,** Box plots illustrating the fractions of wave pacemakers measured in *+/+* (gray) and *+/mut* (blue) mice (Mann-Whitney test with Bonferroni correction for multiple comparisons, P > 0.05 for all comparisons). **e-h,** Median (per mouse) burst (defined as described in Materials and methods) frequencies (e, f) and fractions of bursts among all recorded spontaneous Ca^2+^ signals (g, h), illustrated similarly as described for wave pacemakers in (c, d). The insert in e shows the ΔF/F trace recorded in the CA3 area of the hippocampus. Scale bars: 6 s and 2% ΔF/F f,h, Mixed-design analysis of variances followed by the simple main effect on the group (*+/+* vs. *+/mut*) test, \**P* < 0.05 for all comparisons.

Analysis of mean region-specific Ca^2+^ signals from the cortex and hippocampus revealed no differences in the overall frequency of Ca^2+^ signals between genotypes (including single transients and bursts), their amplitude, or the inter-event and intraburst intervals. However, the burst frequency and the fraction of bursts among Ca^2+^ signals were significantly higher in *+/mut* compared to *+/+* mice across all hippocampal areas except the subiculum. The enhanced burstiness of the predominant pacemaker area (hippocampal network) in the *+/mut* mice aligned well with a higher fraction of waves propagating over larger areas (P < 0.05, bootstrap-based estimation for 1-41, 57-75, and 90-100 percentiles).

In summary, at P3, the presence of the *Scn2a* p.A263V variant alters endogenous network activity in the cortico-hippocampal formation by increasing network burstiness, shifting pacemaking activity toward the hippocampus, and enlarging the size of propagating waves. These changes support the generation of infrequent seizure-like activity, observed through electrical recordings in vivo.

### Intrinsic excitability of hippocampal CA3 and CA1 regions transiently increases in *Scn2a* p.A263V mice during the neonatal period

Given our in vivo LFP recordings and in situ Ca^2+^ imaging results implicating the hippocampus, particularly the CA3 region, as a potential driver of synchronized activity during early development in *Scn2a* p.A263V mutant animals, we analyzed cellular excitability in CA3 and CA1 pyramidal neurons. We focused on two developmental time points in *B6.Scn2a* mice: Neonatal (P10-14) and juvenile (P24-30) (Fig. 7). These time points correspond to periods before and after the developmental switch at the AIS from Na_v_1.2 to Na_v_1.6 as the primary channel responsible for action potential (AP) generation^32^. During the early postnatal period (P10-14), pyramidal neurons in both CA1 and CA3 regions of *mut/mut* and *+/mut* animals exhibited increased excitability compared to their *+/+* counterparts (Fig. 7a, g). This increased excitability was evident from the significantly altered input-output curves, with CA1 neurons showing increased AP firing in response to depolarizing current injections (Fig. 7h). CA3 neurons displayed a similar trend (Fig. 7b) and increased maximal firing rates for both *+/mut* and *mut/mut* animals (Supplementary Table 1). Both CA3 and CA1 neurons in *Scn2a* p.A263V animals demonstrated a lower AP threshold, aligning with their increased excitability. At the same time, resting membrane potentials remained unchanged across all genotypes (Fig. 7c). Other intrinsic properties showed region-specific alterations. In CA3, rheobase decreased in *+/mut* and *mut/mut* animals, while input resistance, AP half-width, and AP peak amplitude were affected only in *+/mut* animals. CA1 neurons, in contrast, exclusively showed changes in AP half-width, which decreased in *mut/mut* animals. To further characterize the effects of the mutation on Na_v_1.2 function, we examined the biophysical properties of sodium currents in CA1 pyramidal neurons using the nucleated patch method. We found no alterations in steady-state activation, inactivation, time of recovery from inactivation, or current density, suggesting that the observed changes in excitability were not directly linked to these sodium current properties (Supplementary Table 1). In contrast to the P10-14 age group, *+/mut* and *mut/mut* AP firing normalized in CA3 and CA1 pyramidal neurons by P24-30 (Fig. 7d, e, j, k). There were no significant differences in AP threshold or other active or passive properties between genotypes in either region at this later stage (Fig. 7f, l). Because of the high mortality of *mut/mut B6.Scn2a* animals, we recorded from only two CA3 pyramidal neurons at this age, an insufficient number for statistical comparisons. These results suggest that the *Scn2a* p.A263V mutation leads to transient hyperexcitability in both *+/mut* and *mut/mut* CA3 and CA1 pyramidal neurons during early postnatal development, which normalizes by P24-30.

**Fig. 7.**
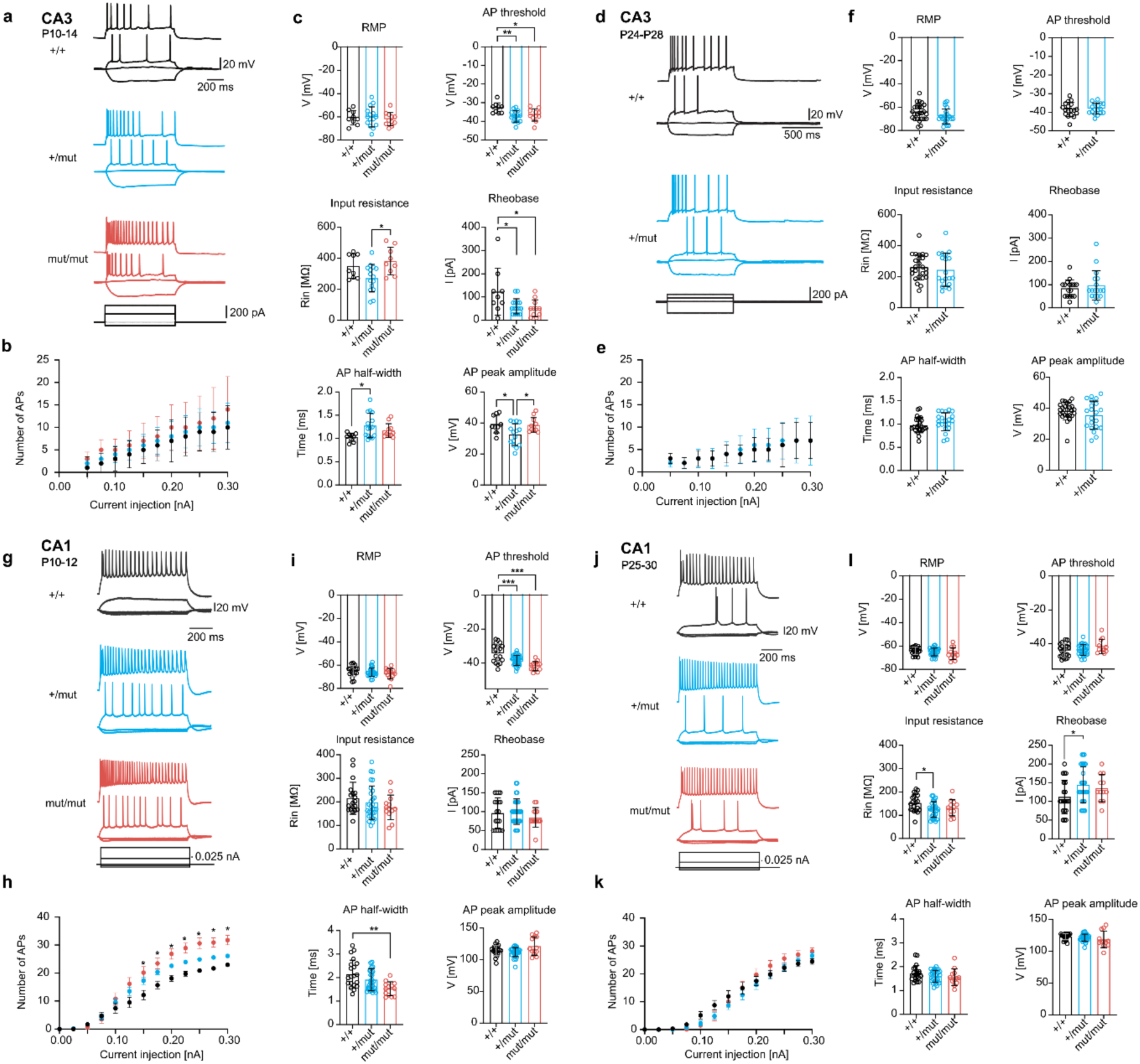
| Transient hyperexcitability in hippocampal pyramidal neurons of *B6.Scn2a* p.A263V mice. Intrinsic properties of CA3 and CA1 pyramidal neurons were assessed at two developmental stages: P10-P14 (a-c, g-i) and P24-P30 (d-f, j-l). **a-c**, Intrinsic properties of CA3 pyramidal neurons at P10-P14. **a**, Representative voltage traces from different genotypes. **b**, Input-output curve showing the number of action potentials (APs) evoked by increasing current injections (mean ± SEM). **c**, Summary of intrinsic neuronal properties: Resting membrane potential (RMP), AP threshold, input resistance, rheobase, AP half-width, and AP peak amplitude. **d-f**, Intrinsic properties of CA3 pyramidal neurons at P24-P28. **d**, Representative voltage traces from different genotypes. **e**, Input-output curve showing the number of APs evoked by increasing current injections (mean ± SEM). **f,** Summary of intrinsic neuronal properties, as in c. **g-i**, Intrinsic properties of CA1 pyramidal neurons at P10-P12. **g**, Representative voltage traces from different genotypes. **h**, Input-output curve showing the number of APs evoked by increasing current injections (mean ± SEM). **i,** Summary of intrinsic neuronal properties, as in c. **j-l**, Intrinsic properties of CA1 pyramidal neurons at P25-P30. **j,** Representative voltage traces from different genotypes. **k**, Input-output curve showing the number of APs evoked by increasing current injections (mean ± SEM). **l**, Summary of intrinsic neuronal properties, as in c. Sample sizes (number of cells per genotype) are provided in Supplementary Table 1. Statistical tests: One-way ANOVA with Bonferroni’s (CA3, *+/+*, *+/mut*, *mut/mut*) or Dunnett’s (CA1, *+/+*, *+/mut*, *mut/mut*) post-hoc test, ANOVA on ranks with Dunn’s post-hoc test (CA1, *+/+*, *+/mut*, *mut/mut*), unpaired t-test (CA3, *+/+*, *+/mut*), or Mann-Whitney test (CA1, *+/+*, *+/mut*); \**P* < 0.05, \*\**P* < 0.01, \*\*\**P* < 0.001.

**Supplementary Table 1.**
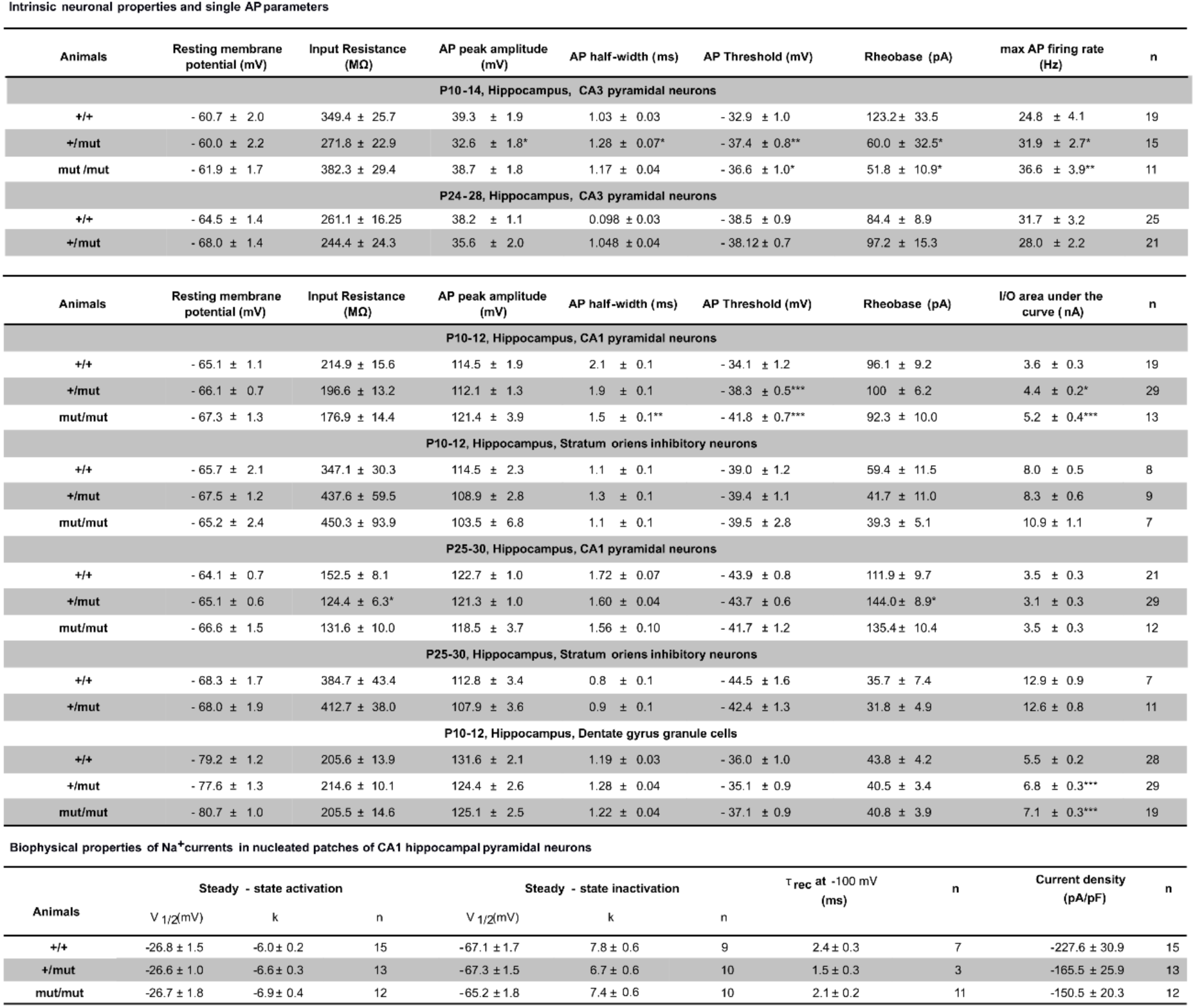
| Summary of intrinsic neuronal and biophysical properties of sodium currents of hippocampal pyramidal neurons. AP = action potential; I/O = input-output curve. Data is presented as mean ± SEM; n, number of recorded neurons; \**P* < 0.05; \*\**P* < 0.01; \*\*\**P* < 0.001. Statistical analyses: For CA3, one-way ANOVA with Bonferroni’s post-hoc test (+/+, +/mut, mut/mut) or unpaired t-test (+/+, +/mut). For CA1, one-way ANOVA with Dunnett’s post-hoc test (+/+, +/mut, mut/mut), ANOVA on ranks with Dunn’s post-hoc test (+/+, +/mut, mut/mut), or Mann-Whitney test (+/+, +/mut).

### Hippocampal hyperexcitability in *B6.Scn2a* mice causes gene dose-dependent accelerated maturation of the hippocampal network

We employed single-cell RNA-sequencing (scRNA-seq) to systematically characterize molecular and cellular mechanisms underlying neuronal hyperexcitability and epileptogenesis across different *B6.Scn2a* mouse line genotypes (+/+, *+/mut*, and *mut/mut*) during critical stages of hippocampal network development. We selected the ages of P4, P7, and P14, which capture the emergence of hippocampal seizure activity (P4), the establishment of hippocampal network hyperactivity and strong seizure phenotype (P7), and the potential onset of homeostatic normalization towards a decrease in neuronal hyperexcitability (P14). Single cells from the dorsal hippocampi at P4, P7, and P14 were collected and processed using the 10x Genomics protocol for scRNA-seq. After preprocessing the data (Suppl. Data Fig. 1-3), cell types and their subtypes were manually annotated based on the known marker genes^38–46^, identifying ten excitatory (Ex) and 14 inhibitory (In) neuronal subtypes (Fig. 8a, Suppl. Fig. 4a-c). In total, 71,428 neurons, from 138,610 cells, were sampled from 39 male *B6.Scn2a* mice – 17,632 neurons at P4 (4 +/+, 4 *+/mut*, and 2 *mut/mut*), 33,837 neurons at P7 (9 +/+, 4 *+/mut*, and 3 *mut/mut*), and 19,959 neurons at P14 (5 +/+, 3 *+/mut*, and 5 *mut/mut*) (Suppl. Table 2). The *Scn2a* transcript was pan-neuronally detected throughout all postnatal ages and genotypes (Fig. 8b). To investigate the extent to which the *Scn2a* p.A263V variant perturbed the transcriptome of hippocampal neurons, we first calculated the overall gene expression distance (as a proxy for the magnitude of gene expression change^47^) between wildtype and mutant neurons across maturational trajectory (Fig. 8c). The overall gene-expression distances demonstrated a clear gene-dose-dependent effect, with *mut/mut* neurons showing larger transcriptional differences than *+/mut* at both P7 and P14 (Fig. 8c). Moreover, this effect was most pronounced at P7, when the gene expression divergence peaked, suggesting that this stage represents a critical point for the gene dose-dependent changes. By P14, the differences had partially resolved, particularly in *+/mut* mice, although *mut/mut* neurons continued to display more pronounced divergence (Fig. 8c).

**Fig. 8.**
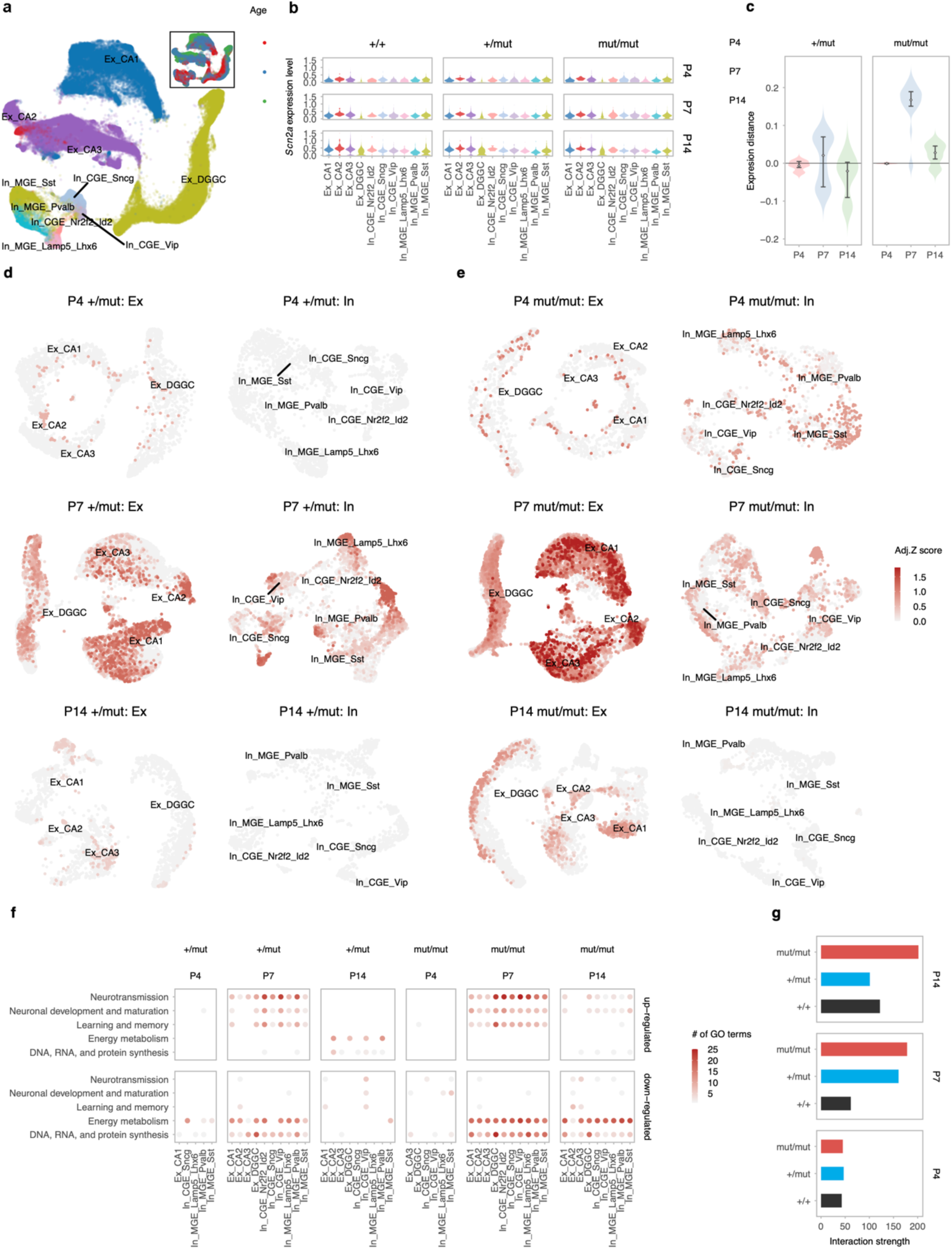
| *Scn2a* gene dose-dependent accelerated maturation of hippocampal networks identified by scRNA-seq. **a,** Violin plots showing the expression level of *Scn2a* across all neuronal subtypes (medium-resolution annotations) in +/+, +/*mut*, and *mut*/*mut* mice at P4, P7, and P14. **b,** UMAP representation of analyzed neurons, colored by medium-resolution annotations; inset shows ages. **c,** Violin plots showing inter-sample gene expression distances between mutant and wildtype neurons at P4, P7, and P14. **d,** Graph-based analysis of expression changes in excitatory and inhibitory neurons in +/*mut* mice at P4, P7, and P14. UMAP plots display the extent of gene expression changes between +/*mut* and +/+ mice, colored by adjusted significance levels. **e,** Same as **d** for *mut*/*mut* versus +/+ mice. **f,** Dot plots showing the numbers of significantly enriched GO Biological Process terms (Benjamini-Hochberg-adjusted *P* value < 0.05) related to specific topics in the top up- and down-regulated genes across neuronal subtypes (medium-resolution annotations) between mutant and wildtype mice at P4, P7, and P14, colored by the number of significantly enriched GO terms. **g,** Bar plots showing the overall interaction strength of inferred cell-cell communication networks in neurons from mutant and wildtype mice at P4, P7, and P14.

Further analysis of gene expression changes using a graph-based approach^47^ revealed widespread transcriptional alterations across multiple neuronal subtypes at P7 in both excitatory (Ex) and inhibitory (In) neurons (Fig. 8d,e). Remarkably, between postnatal days P4 and P7, we observed substantial transcriptome-wide alterations in neuronal gene expression profiles of *B6.Scn2a* mice (Fig. 8d,e), suggesting that the early neural network hyperactivity detected at P2.5 did not immediately trigger comprehensive molecular remodeling. These subtle transcriptional changes at P4 may be early molecular triggers that prime seizure-supporting neuronal network maturation by P7. By P14, the substantial reduction in gene expression changes (Fig. 8d,e) likely reflected the normalization of somatic hyperexcitability observed previously (Fig. 7, Supplementary Table 1). To characterize these developmental dynamics systematically, we calculated gene expression distances for each neuronal subtype in *+/mut* and *mut/mut* mice across P4-P7-P14. The result confirmed a massive and transient transcriptional shift at P7 (Fig. 8c-e, Suppl. Fig. 4). The gene expression landscape varied specifically between genotypes: in *+/mut* mice, dentate gyrus granule cells and parvalbumin-positive interneurons exhibited the most pronounced expression changes, while these changes were additionally observed in CA1 neurons of *mut/mut* mice (Suppl. Fig. 4d,e). To evaluate whether the *Scn2a* p.A263V variant influenced hippocampal neuronal composition during network maturation, we employed two complementary analytical approaches: Compositional data analysis (CoDA)^47^ and graph-based cell density difference estimation (Suppl. Fig. 5). Whereas no statistically significant global changes in neuronal subtype composition were detected across genotypes, we observed a prominent trend toward an increased magnitude of changes in *mut/mut* mice. Notably, these alterations were not characterized by a uniform neuronal decrease but rather by nuanced and differential changes across various neuronal subtypes. This pattern suggests potential subtle perturbations in neuronal differentiation programs induced by seizure-related molecular mechanisms.

To elucidate the molecular pathways perturbed by the *Scn2a* p.A263V variant, we performed gene set enrichment analysis (GSEA) on highly expressed genes, focusing on Gene Ontology (GO) Biological Process terms. Mirroring the gene-dose-dependent transcriptomic changes, the number of enriched GO terms peaked at P7 and subsequently declined by P14 (Extended Data Fig. 7). Of note, *mut/mut* mice maintained a higher number of enriched GO Biological Process terms at P14, with more affected neuronal subtypes compared to *+/mut*, suggesting persistent molecular consequences of the mutation. Systematic categorization of GO terms revealed significant enrichment in developmental, maturation, and neurotransmission pathways at P7 (Fig. 8f, Extended Data Fig. 8). This underscores P7 as a pivotal period of neuronal hyperexcitability and accelerated maturation. The developmental impact was transient in *+/mut* mice but remained pronounced in *mut/mut* mice, indicating gene dosage-mediated molecular programming. The *Scn2a* p.A263V variant profoundly disrupted fundamental cellular processes. Energy metabolism, DNA, RNA, and protein synthesis pathways were transiently downregulated at P7 in *+/mut* mice and persistently altered at P7 and P14 in *mut/mut* mice (Fig. 8f, Suppl. Fig. 6). These perturbations suggest a fundamental disruption of neuronal homeostatic mechanisms that may contribute to the premature mortality observed in *mut/mut* mice. GO terms associated with learning and memory demonstrated widespread upregulation by P7 in both *+/mut* and *mut/mut* mice (Fig. 8f, Suppl. Fig. 7). This early molecular signature potentially foreshadows the cognitive and neuronal network changes subsequently detected in the *Scn2a* lines (Figs. 3,4; Extended Data Fig. 5), linking early network hyperactivity to long-term neurodevelopmental consequences.

To investigate network reorganization during hippocampal maturation, we conducted a cell-cell interaction analysis using CellChat. A striking observation emerged: *+/mut* mice exhibited accelerated neuronal interaction dynamics, reaching peak neuron-neuron interaction strength one week earlier than +/+ mice—specifically at P7 instead of P14 (Fig. 8g, Suppl. Fig. 8). This finding suggests a precocious neuronal maturation trajectory in *+/mut* mice. The interaction strength dynamics revealed nuanced genotype-specific patterns. In *+/mut* mice, neuron-neuron interaction strength stabilized between P7 and P14, whereas *mut/mut* mice displayed a continued increase in interaction strength (Fig. 8g, Suppl. Fig. 8). These progressive dynamics potentially indicate a gene-dosage-dependent mechanism of persistent neuronal network reorganization characterized by ongoing acceleration of neuronal maturation and hyperactive network development.

To further dissect the temporal dynamics of hippocampal neuron development, we applied RNA velocity-based pseudotime inference across excitatory and inhibitory neuronal subtypes at P4, P7, and P14. UMAP embeddings revealed coherent maturation trajectories across DG, dorsal CA3, and CA1 excitatory neurons, as well as *Pvalb*, *Sst*, and *Vip* inhibitory populations (Extended Data Fig. 9a–b). Compared to +/+ controls, both +/*mut* and *mut*/*mut* neurons exhibited accelerated pseudotemporal progression, with shifts detectable as early as P4 in many subtypes. However, these early changes were not statistically significant in DG (+/*mut*), *Pvalb* (both genotypes), and *Vip* (+/*mut*). The divergence peaked at P7 and was widespread across genotypes and subtypes (Extended Data Fig. 9c–f; Extended Data Fig. 10a–h). By P14, pseudotime distributions in +/*mut* mice showed reversal relative to wildtype in DG, CA1, and *Vip* neurons, indicating potential overcompensation. A milder but significant reversal was also observed in dorsal CA3. In contrast, *Pvalb* and *Sst* neurons in +/*mut* mice showed no significant changes at this stage. *Mut*/*mut* neurons, by comparison, consistently maintained or further exaggerated their advanced maturation profiles. These findings, supported by ECDF analyses and two-sided Kolmogorov–Smirnov tests, revealed dynamic and cell-type-specific effects of *Scn2a* gene dosage on neuronal maturation, marked by early acceleration and, in some cases, a reversal of trajectory in heterozygous mice at later stages.

Collectively, our scRNA-seq analyses show that the Scn2a p.A263V variant triggers a gene-dose-dependent acceleration of neuronal transcriptomic maturation, with a pronounced molecular signature emerging at P7—a pivotal developmental window characterized by heightened neuronal hyperexcitability. The mutation’s profound impact manifests through a comprehensive disruption of fundamental cellular processes, including neurotransmission and energy metabolism, identifying P7 as a critical nexus for epileptogenic network transformation in our GOF *Scn2a* disease models.

**Extended Data Fig. 7.**
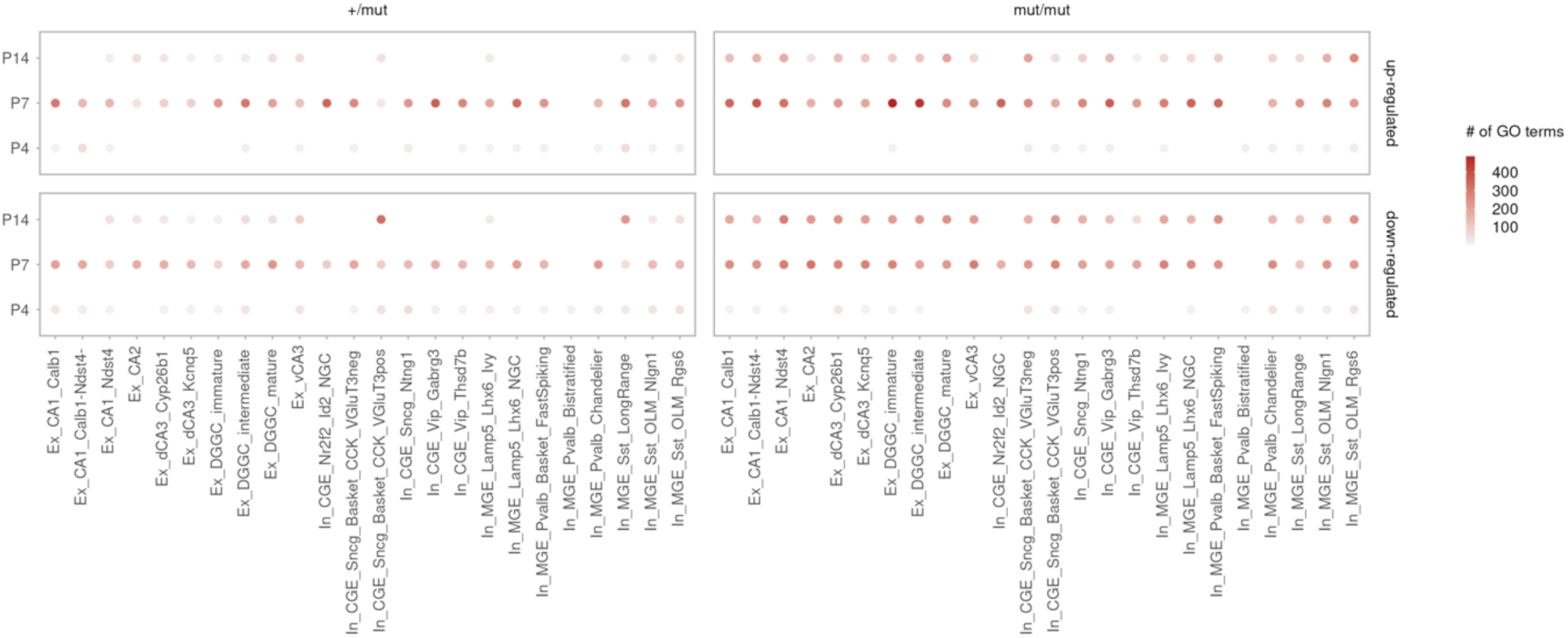
| Changes in GO Biological Processes in *Scn2a* mutant mice. Dot plots showing the numbers of significantly enriched GO Biological Process terms (Benjamini-Hochberg-adjusted *P* value < 0.05) related to specific topics in the top up- and downregulated genes across neuronal subtypes (high-resolution annotations) between mutant and wildtype mice at P4, P7, and P14, colored by the number of significantly enriched GO terms.

**Extended Data Fig. 8.**
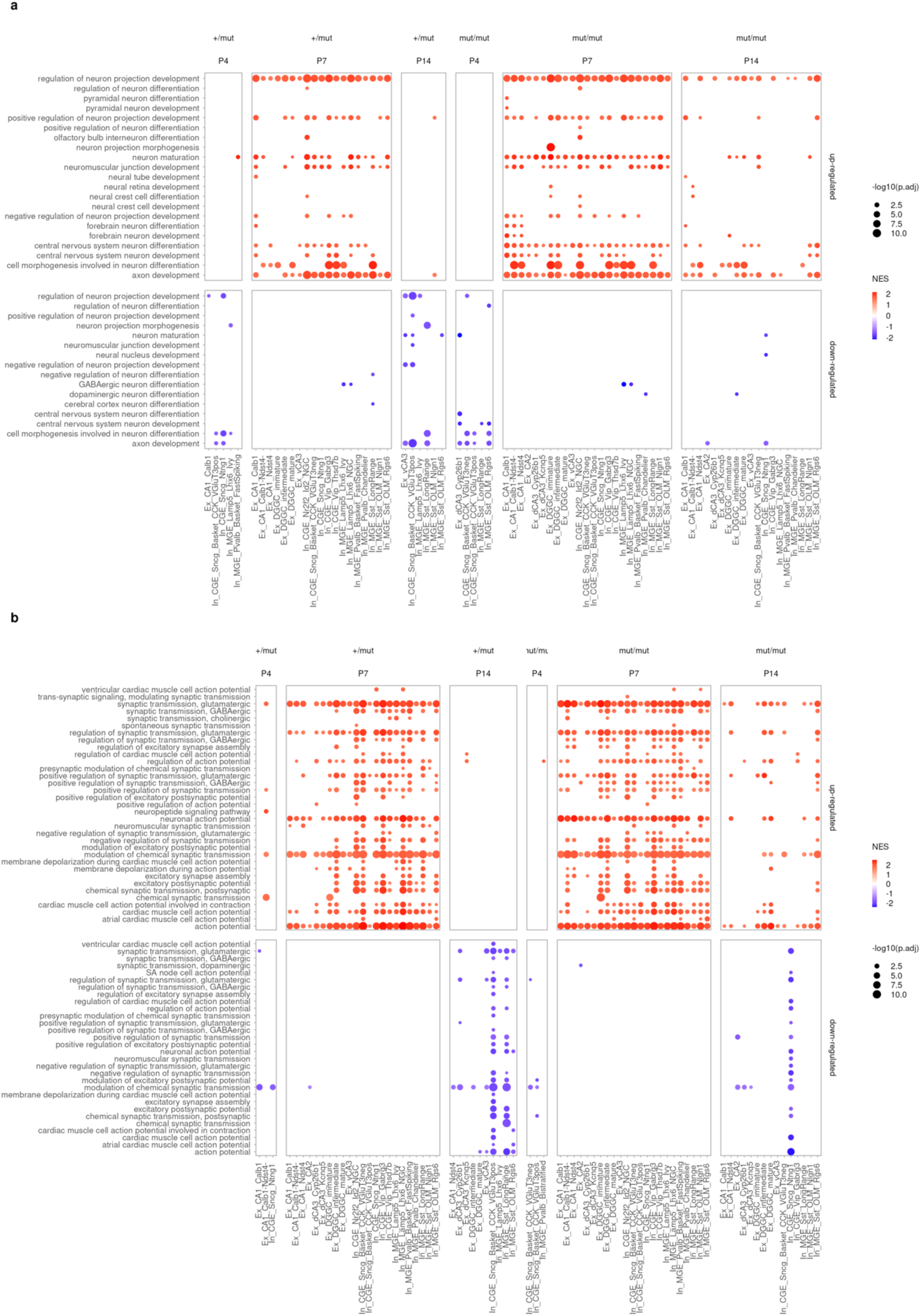
| Changes in GO Biological Processes related to neuronal development and neurotransmission in *Scn2a* mutant mice. **a**, Dot plots showing significantly enriched GO Biological Process (BP) terms associated with neuronal development and maturation in the top up- and down-regulated genes across neuronal subtypes (high-resolution annotations) between mutant and wildtype mice at P4, P7, and P14, colored by normalized enrichment score (NES) and sized by -log10(BH-adjusted *P* value). **b,** Same as **a** for GO BP terms associated with neurotransmission.

**Extended Data Fig. 9.**
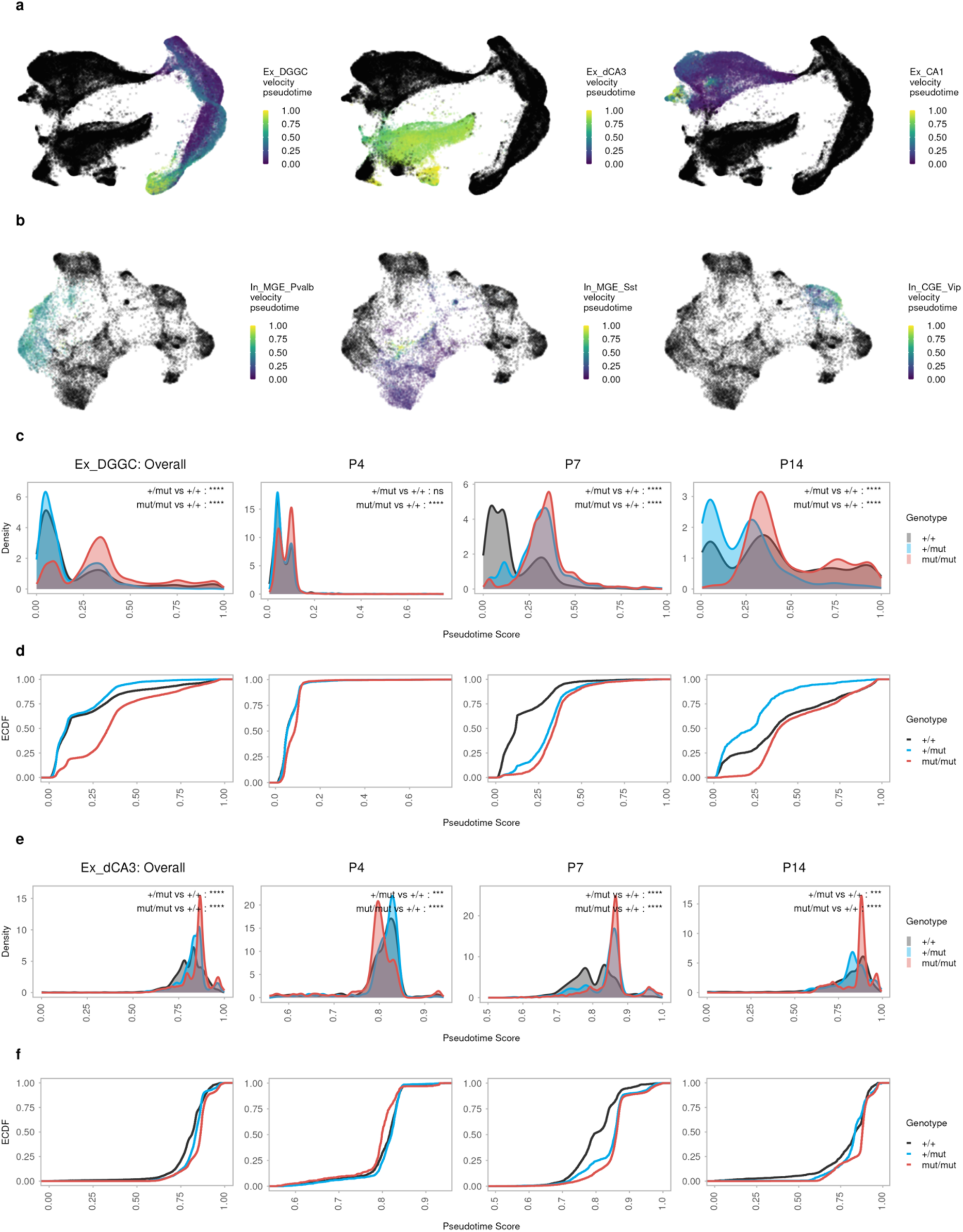
| Gene dose-dependent alterations in mutant mice in pseudotemporal distribution of hippocampal neurons. **a**, UMAP representation of all measured excitatory neurons, colored by inferred velocity pseudotime in DG, dorsal CA3 and CA1 excitatory neurons (from left to right). **b**, Same as a for measured inhibitory neurons, colored by inferred velocity pseudotime in *Pvalb*, *Sst* and *Vip* inhibitory neurons (from left to right). **c**, Density plots showing the distribution of DG excitatory neurons along the pseudotime between genotypes in all samples and separately in P4, P7, and P14 samples. Significance was assessed using the two-sided Kolmogorov–Smirnov (KS) test. *P* values are indicated as follows: ns (*P* ≥ 0.05), * (*P* < 0.05), ** (*P* < 0.01), *** (*P* < 0.001), **** (*P* < 0.0001). **d**, Corresponding empirical cumulative distribution function (ECDF) plots showing the same pseudotime distributions as in c, across genotypes and developmental stages. **e**, Same as c for dorsal CA3 excitatory neurons. **f**, Same as d for dorsal CA3 excitatory neurons.

**Extended Data Fig. 10.**
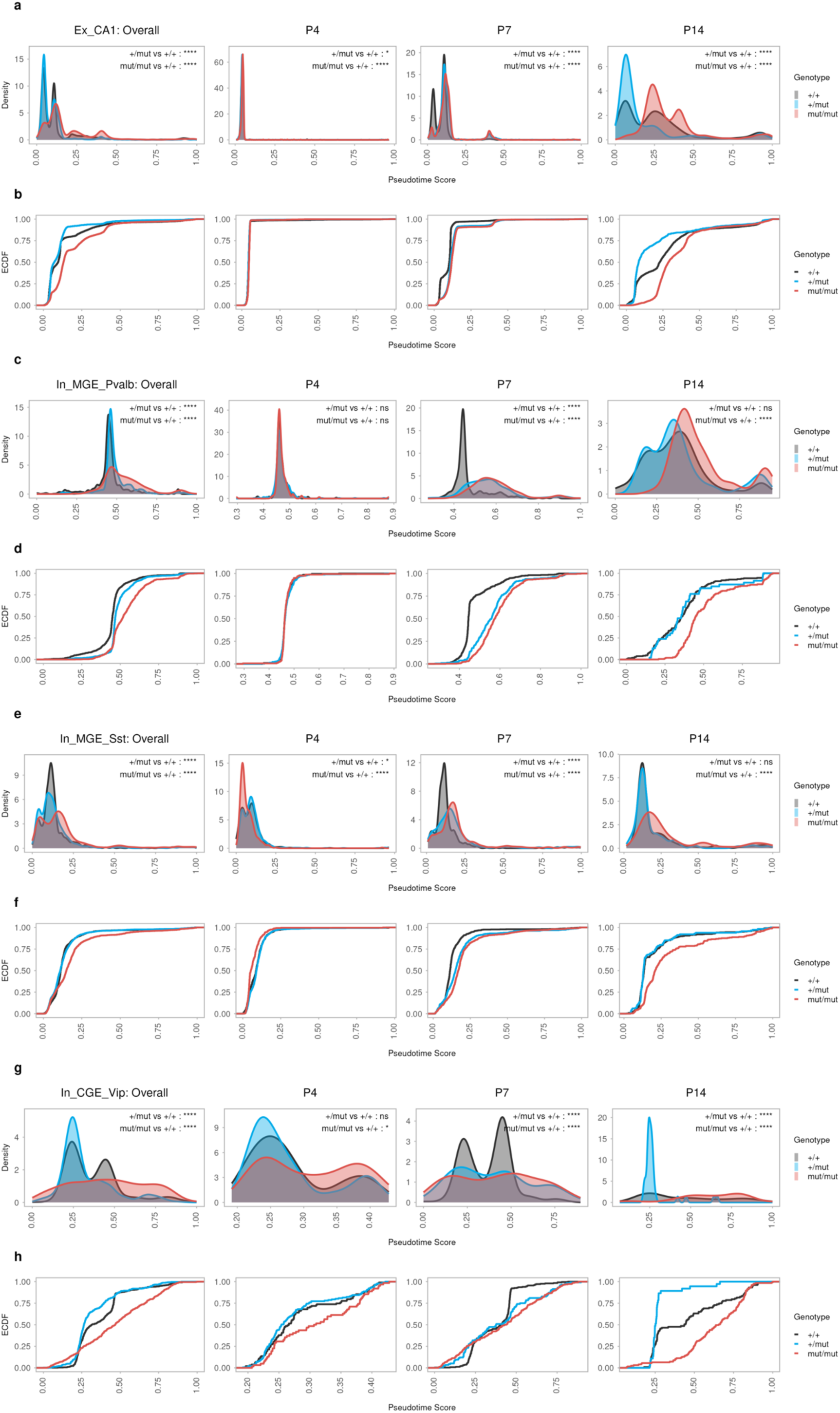
| Neuron type-specific pseudotemporal distribution changes in CA1 excitatory neurons and inhibitory neurons. **a,** Density plots showing the distribution of CA1 excitatory neurons along the pseudotime between genotypes in all samples and separately in P4, P7, and P14 samples. Significance was assessed using the two-sided Kolmogorov–Smirnov (KS) test. *P* values are indicated as follows: ns (*P* ≥ 0.05), * (*P* < 0.05), ** (*P* < 0.01), *** (*P* < 0.001), **** (*P* < 0.0001). **b,** Corresponding ECDF plots showing the same pseudotime distributions as in **a**, across genotypes and developmental stages. **c,** Same as **a** for *Pvalb* inhibitory neurons. **d,** Same as **b** for *Pvalb* inhibitory neurons. **e,** Same as **a** for *Sst-*inhibitory neurons. **f,** Same as **b** for *Sst* inhibitory neurons. **g,** Same as **a** for *Vip* inhibitory neurons. **h,** Same as **b** for *Vip* inhibitory neurons.

## Discussion

*SCN2A* GOF variants remain largely unexamined at the molecular, cellular, and network levels. While the increased excitability of excitatory cortical neurons has been implicated^48,49^, the complete time course and underlying mechanisms of epileptogenesis are yet to be determined. To address this knowledge gap, we generated mouse lines carrying a pathogenic *SCN2A* variant linked to neonatal seizures in children. Our *Scn2a* p.A263V mouse lines replicate key phenotypic features observed in patients with *SCN2A*-linked DEEs and mice harboring GOF *Scn2a* variants. These features include early postnatal seizure onset, increased mortality, and responsiveness to sodium channel blockers^48,50^, demonstrating construct, face, and predictive validities. Mirroring the variable phenotype expressivity seen in patients^50^, our mouse lines exhibit gene dose and genetic background-dependent phenotypes. Homozygous animals display more severe seizures, increased mortality, and subtle behavioral changes, while heterozygous mice show milder symptoms and a self-limiting epilepsy phenotype.

In early postnatal stages, when Na_v_1.2 channels are the predominant voltage-gated sodium channels responsible for action potential generation, the *Scn2a* p.A263V GOF variant is linked to increased intrinsic neuronal excitability and elevated c-Fos levels in all hippocampal subregions. By showing that somatic excitability in mutant glutamatergic neurons normalizes at about four weeks of age, we establish a link between the known^19^ Na_v_1.2-to-Na_v_1.6 switch at the AIS during the third postnatal week and changes in action potential generation. To investigate how the increased cellular excitability is linked to network excitability, we employed a comprehensive strategy focused on the early postnatal hippocampus. In situ calcium imaging in cortico-hippocampal slices revealed an increased incidence and frequency of hippocampal calcium bursts in three-day-old heterozygous mice, indicating elevated network excitability of the neonatal hippocampus. The increase in c-Fos staining at P7 in all hippocampal subregions and in the deep MEC layers receiving the hippocampal output also suggests primarily hippocampal hyperexcitability at this age. This finding was supported by in vivo hippocampal depth recordings from awake mouse pups as early as P2.5, which revealed spontaneous electrographic hippocampal seizures that most often did not have a behavioral correlate, akin to electroclinical uncoupling in human neonates^51^.

CA1 LFP depth profile analysis showed beta/low-gamma frequency oscillations at seizure onset in the stratum radiatum and oriens layers receiving CA3 input, implicating CA3 in seizure initiation. The CA3 region contributes to early hippocampal synchrony through the action of GABAergic hub neurons orchestrating network synchronization in the developing hippocampus^52^. The CA3 region of the adult hippocampus is primed to generate synchronous network activity at gamma frequencies due to the combination of extensive recurrent connectivity and perisomatic feedback inhibition^53–55^. The latter is not assumed to mature before the second postnatal week, when CA1 ripple oscillations appear^56^. While short (<200 ms, 25 Hz) gamma bursts in CA1 stratum radiatum of wildtype rats were detected as early as P2^57,58^, we observed spontaneous CA3-driven early gamma oscillations lasting several seconds in P2.5 *Scn2a* GOF mice, suggesting premature maturation of CA3 circuits. Previous studies showed a strong age dependence of kainate-induced activity in the isolated intact neonatal rat hippocampus, where kainate-evoked early gamma oscillations and ictal-like activity did not emerge before the end of the first postnatal week^59,60^. Interictal spikes induced by contralateral kainate infusion^61^ resembled iSPWs in our neonatal recordings (current sink in stratum oriens), with their incidence increasing after the first seizure (Fig. 5). Notably, HFS of the VHC in P5-7 control mice resulted in a more pronounced increase in the occurrence of CA1 iSPWs rather than eSPWs, lasting for more than 30 min (Extended Data Fig. 6), which suggests that bilateral stimulation of CA3 preferentially excites the basal dendrites of CA1 pyramidal cells in stratum oriens. These dendrites not only receive more excitatory input from contralateral than from ipsilateral CA3 pyramidal neurons but also preferentially target CA1 cells with axon-carrying dendrites (AcD cells)^36^. Excitatory input to AcD branches can evade perisomatic inhibition and trigger action potentials in the attached axon^62^. The AIS of AcD cells also contains fewer inhibitory synapses and shows limited homeostatic plasticity^63^ compared to non-AcD cells. Therefore, in our *Scn2a* mice, AcD cells may contribute to network synchronization as observed during (CA3-driven) ripple oscillations, when they exhibit preferential firing^64^.

The abnormal CA3-driven network patterns prompted us to systematically characterize the molecular mechanisms driven by early neuronal hyperexcitability during postnatal development using scRNA-seq. These experiments revealed a notable age and gene-dose-dependent gene expression change within glutamatergic and GABAergic neurons across all hippocampal subfields. At P4, only minor expression shifts were detectable, even though the mice had already experienced electrographic seizures. Thus, the significant expression shifts at P7 likely reflect network hyperactivity-induced changes. By the second postnatal week, the divergence between self-limiting heterozygous and more severe homozygous phenotypes becomes apparent through transient versus persistent gene expression shifts, changes in pathways related to development, maturation, and energy metabolism, and distinct PV^+^ interneuron maturation trajectories. These early transcriptional differences are reflected in morphological, behavioral, and electrophysiological differences observed between the two conditions in adulthood.

Our findings demonstrate that the neonatal hippocampal seizures associated with the *Scn2a* p.A263V variant induce persistent molecular and network-level alterations despite limited gross morphological changes. The reduction in juvenile PNNs and adult PV expression in hippocampal subfields, particularly CA3, suggests impairment of inhibitory microcircuits. PV+ interneurons are critical for generating gamma oscillations and maintaining excitatory-inhibitory balance^65^, while PNNs stabilize synaptic connections during critical periods^66^. The observed decreases in WFA+ PNNs at P21 and PV immunoreactivity in adults likely reflect activity-dependent remodeling of inhibitory networks, consistent with the altered theta-gamma coupling and reduced mid-gamma oscillation frequency in hippocampal recordings. These network changes persisted into adulthood despite normalization of CA3 pyramidal cell excitability by P24-30, indicating seizure-induced maladaptive plasticity during early developmental windows. Of note, the behavioral consequences remained relatively modest except in *B6.Scn2a* mut/mut mice. This dissociation between substantial neonatal network-level perturbations and preserved behavioral output underscores the remarkable resilience of developing hippocampal networks and suggests a high therapeutic potential for early intervention. The detection of electrographic seizures as early as P2.5, before the developmental switch from Na_v_1.2 to Na_v_1.6 dominance at the AIS, indicates the need for early intervention in SCN2A DEEs. Our scRNA-seq data show accelerated transcriptomic maturation at P7, which coincides with peak seizure burden, suggesting that hyperactivity drives the observed molecular reprogramming. While postnatal phenytoin administration prevented acute seizures, the ideal therapeutic window likely precedes seizure onset. In humans, in-utero treatment would be required because Na_v_1.2 is expressed as early as in the second trimester (Suppl. Fig. 9)^67^. In current clinical practice, sodium channel blockers or ASO treatment are typically initiated following diagnosis (weeks to months after birth), likely missing critical periods when aberrant activity patterns develop^68^. Our findings align well with the growing recognition that neurodevelopmental disorders such as *SCN2A* DEEs require early intervention to mitigate this abnormal excitability during circuit formation for optimal outcomes^69^. The temporally restricted nature of the CA3 hyperexcitability and its driving role in network changes make such early treatments especially promising for SCN2A GOF.

## Methods

### Animals

This study used two mouse lines carrying the Scn2a p.A263V mutation: The *Scn2a* mutant line on a C57BL/6J background^21^ (*B6.Scn2a*) and a newly generated *CD1.Scn2a* line created by backcrossing onto a CD1 background for at least ten generations. Mice were housed in type II long plastic cages under standard conditions (21 ± 2°C, 50% relative humidity) with food (ssniff Spezialitäten GmbH, Soest, Germany) and water available ad lib. Housing included individually ventilated cages or cabinets (SCANBUR) with an inverted 12:12 dark:light cycle (lights on at 10 p.m.). All animal care and experimental procedures were conducted following institutional guidelines, the German Animal Welfare Act, and the European Communities Council Directive 2010/63/EU. The experiments were approved by the Ministry for Health and Consumer Protection (Hamburg, Germany) and the North Rhine-Westphalia State Agency for Nature, Environment, and Consumer Protection (LANUV NRW, Germany).

### Immunohistochemistry

Animals were deeply anesthetized with an intraperitoneal injection of anesthetics (12% ketamine, 8% xylazine in physiological saline) and transcardially perfused with phosphate-buffered saline (PBS) followed by 4% paraformaldehyde (PFA, Roti® Histofix, Roth, Germany). Brains were post-fixed overnight at 4°C in 4% PFA. For GFAP, WFA, PV, NPY, and c-Fos staining, brains were transferred to 20% sucrose in PBS for cryoprotection. After 2 days, brains were frozen in pre-cooled (−80°C) 2-methyl-butane (isopentane, Roth, Germany) and stored at −80°C until further processing. Coronal sections were cut at 20 μm thickness using a cryostat (Leica, Germany). All stainings followed a general protocol of washing, blocking, primary antibody incubation, secondary antibody incubation, and final washing steps, with specific variations noted below. Washing and blocking steps varied as follows: For GFAP, sections were washed three times for 5 min using PBS, then blocked with 5% normal goat serum (NGS, Vector, USA), 0.3% TritonX-100 (Sigma, USA) and 0.2% bovine serum albumin (BSA, Sigma, USA) for 1-2 hours at room temperature. For WFA and NPY, sections were washed three times for 10 min in 0.1% PBST (0.1% Triton X-100 in PBS) on a shaker at room temperature and then blocked with 5% normal goat serum (NGS) in 1x PBS for 1 hour at room temperature. For PV, sections were washed three times for 5 min in PBS, once for 10 min in PBST, and once for 5 min in PBS on a shaker at room temperature and then blocked with 5% NGS in PBST for 1 hour at room temperature. Primary antibodies were incubated overnight at 4°C: Rabbit polyclonal anti-GFAP (1:500, Dako #Z033429, USA), biotinylated WFA (1:500, Sigma #L1516, USA), mouse monoclonal anti-parvalbumin (1:200, Abcam, UK), and rabbit polyclonal anti-NPY (1:1000, Immunostar, USA). Secondary antibodies were incubated for 2 hours at room temperature: Alexa546 goat anti-rabbit (1:500, Molecular Probes, Netherlands) for GFAP, Alexa546 streptavidin (1:500, MP#S11225, Netherlands) for WFA, Alexa546 goat anti-mouse (1:500, Molecular Probes, Netherlands) for PV, and Alexa546 goat anti-rabbit (1:500, Molecular Probes, Netherlands) for NPY. The final washing steps were as follows: For GFAP, sections were washed in PBS (3 times, 5-10 min each). For WFA and NPY, sections were washed three times for 10 min in PBST and three times for 10 min in PBS. For PV, sections were washed five times for 5 min in PBS. All sections were mounted with DAPI Fluoromount-G (Southern Biotech, USA). For c-Fos staining, sections were washed with PBS and incubated in 0.3% H_2_O_2_ solution for 30 min to inhibit endogenous peroxidases. Sections were then washed 3 times for 15 min and then blocked with 5% normal goat serum and 0.2% TritonX-100 in PBS for 1 hour at room temperature. Blocked sections were incubated with polyclonal rabbit anti-c-Fos (1:1000, Calbiochem, Germany) overnight at 4°C. After washing three times for 15 min in PBS, sections were incubated with biotinylated goat anti-rabbit (1:200, Vector, USA) for 2 hours at room temperature. Sections were then incubated for 2 hours with the preformed avidin-biotin complex using VECTASTAIN® ABC-Elite Kit (Vector, USA) prepared according to manufacturer’s instructions, washed 4 times for 5 min in PBS and subsequently incubated in DAB substrate solution (SIGMA FAST™ 3,3’-Diaminobenzidine tablets, Sigma, USA) until staining became visible. To stop the reaction, slices were washed 4 times in water. Sections were dehydrated through an ethanol series (70%, 90%, 98%, twice 100% ethanol and twice Roti® Histol (Roth, Germany) and coverslipped with Roti® Histokitt (Roth, Germany).

Images were acquired on a Zeiss LSM 700 laser scanning confocal microscope with Zen 2012 SP5 FP3 black edition software, using either a Zeiss Fluar ×5/0.25 objective (0.5 zoom, pixel size: 2.5 × 2.5 µm2) or a Zeiss Plan-Apochromat ×20/0.8 objective (1.0 zoom, pixel size: 0.3126 × 0.3126 µm2). Image analysis and fluorescent signal quantification were performed using ImageJ 1.49v software (National Institutes of Health, USA). For c-Fos quantification, stereological analyses were performed using Stereo Investigator Software Version 10.01 (MicroBrightField) on an AxioImager microscope. Experimenters performed the analysis blinded to the experimental conditions and the examined hemisphere. Sections cut at distances between 3 and 4 mm caudally to the rostral pole of the forebrain were analyzed. The hippocampal region was subdivided, and the areas of the whole hippocampus, pyramidal cell layers CA1 and CA3, and granular cell layer in the dentate gyrus were measured separately. The entorhinal cortex was divided into superficial layers (L1-L23) and deep layers (L43-L6). Objects within dissector squares at 100-μm distances, with a dissector depth of 12 μm, were counted using a 20x magnification objective. Three sections were evaluated per animal and staining. The background was removed for representative images, and the contrast was linearly adjusted.

### Behavioral tests

All behavioral tests were conducted on adult mice older than 8 weeks. Both C57BL/6J (*B6.Scn2a*) and CD1 (*CD1.Scn2a*) lines, and both males and females, were used for each test. Exceptions were as follows: The Morris Water Maze and Contextual Fear Conditioning tests were performed only on B6 mice; the Passive Avoidance task was conducted only on CD1 mice; and the One-Trial Spatial Learning test was performed only on male B6 mice. The Open Field (OF) test was used to assess anxiety and exploratory behavior. Mice were placed in one corner of an illuminated (100 lux) white box (50 x 50 x 40 cm) and allowed to roam freely for 15 min. Each animal was tracked and recorded using EthoVision XT 14 (Noldus, Netherlands). The arena was subdivided into a border (5 cm from the edge) and a center area (20 x 20 cm square in the middle). Each animal was placed on the lower left side, with the head facing the corner. The testing period began after 3 seconds of movement recognition within the arena. Analysis parameters included total distance moved [m], distance moved per time slot [m], velocity [m/s], time in center, and time in border [%].

The Y-Maze test was used to assess working memory. The apparatus consisted of three equally sized arms (34 x 5 x 30 cm) made of gray plastic. Animals were placed in the center of the arena and allowed to freely explore until they reached 24 transitions, or for a maximum of 15 min. Each entry into a new arm was considered a transition, and an alternation was recorded when the two other arms had been visited beforehand. For a proper turn, an animal had to enter one of the three arms and transition to another one with its entire body. The test was conducted on two consecutive days, and the average was used for analysis. Parameters analyzed included percent alternations [%] and average transition time [s].

The contextual fear conditioning (CFC) test investigated the ability of mice to learn and remember an association after an aversive experience. Animals were conditioned in a dark chamber with black-white striped walls (Biological Research Apparatus, Model: 46590-PB; 40500-001, Ugo Basil S.R.L.), the dimensions of which were 18 x 18.5 x 26 cm. The chamber was illuminated at 8 lux. The test consisted of a 120-second baseline followed by 120 seconds with three 1-second shocks (0.45 μA) at 120, 160, and 200 seconds. After 24 hours, the animals were placed in the same environment for 240 seconds without shocks. Freezing behavior (defined as no movement for at least one second) was analyzed using EthoVision XT 14. The cumulative duration [%] of freezing on the test trial was used for further analysis.

The passive avoidance test was used to assess associative learning in the CD1 line due to the altered performance of this line in CFC^70,71^. The arena consisted of a dark chamber (18 x 18.5 x 26 cm, 30-45 lux) with a grid floor and a light chamber (18 x 18.5 x 26 cm, 1245-1285 lux) with an even floor surface. On day one, mice were placed into the light chamber with the nose facing the wall opposite the door. The door between the chambers was initially closed. After 1 second of movement recognition, the trial started. After 30 seconds, the door was manually opened. When the mouse entered the dark chamber with its whole body, the door was closed immediately, and a 2-second shock (0.5 μA) was applied. The trial ended 20 seconds after the shock. On day two, the mice were placed in the arena as before, but the door remained open for the entire 300-second trial. The latency to enter the dark chamber on the second day was analyzed.

The one-trial spatial learning test assessed spatial learning under minimally anxiogenic conditions^72^. The apparatus was a square arena (84 × 84 cm) with transparent Plexiglas walls, elevated 82 cm from the ground. Three small compartments were located in three corners. The arena was indirectly illuminated (65 lux) and surrounded by dark curtains. Male mice explored the apparatus for 20 min, with three unfamiliar female mice placed in one compartment and the other two containing only nesting material. Small holes in the compartment walls facilitated odor dispersion. A recall trial was conducted 24 hours later using an identical apparatus without female mice. The time spent near the target compartment versus other compartments was measured using Ethovision XT 14. The female position was randomized between trials to avoid side bias.

The Morris Water Maze test was used to assess spatial learning and memory, following the standard hidden-platform acquisition and retention version of the Morris Water Maze^73^. The arena consisted of a circular pool (145 cm in diameter, 40 cm walls above water surface area, water temperature: 22.0°C ± 1.0°C). Non-toxic white paint was used to stain the water opaque. Mice underwent a 2-day pretraining protocol for habituation, followed by a 5-day acquisition phase with four daily trials. Probe trials were conducted on days 3, 4, 5 of acquisition, and 2 days following acquisition.

The accelerated RotaRod test was used to assess motor learning. Animals walked on a turning, corrugated rod (3.2 cm diameter, Jones & Roberts, TSE Systems). Six trials were performed for each animal. Trials 1 and 2 were conducted at a constant speed of 4 rpm for 3 min. For subsequent trials, the speed was increased from 4 rpm to 40 rpm within 4 min, with a maximum trial time of 5 min. Animals were retested after 24 hours at increasing speed. Performance was measured by latency to fall. Mice were allowed one passive rotation before being gently removed from the rotarod. The trial was then considered complete.

The CatWalk XT system (Noldus) was used for automated gait analysis. Mice ran on an enclosed walkway on a glass plate. Run duration was set to 0.5–20 s, with a maximum speed variation of 60% and a minimum of 10 consecutive steps. At least eight runs per mouse were collected. Runs were classified with the CatWalk XT 10 software, and footprints were automatically detected and manually corrected by visual inspection. The parameters analyzed included running speed, stand (duration of paw contact with the glass plate), stride length, step cycle, base of support (BOS; average width between front or hind paws), step sequence, and regularity index (percentage of normal step sequence patterns relative to total paw placements).

### In situ *Ca^2^*^+^ *Imaging*

Horizontal brain slices (thickness 500 µm) were prepared from P3 mice as described previously^37,74^. For consistency, we only used slices located 1.5-2 mm higher than the bottom of the brain from each animal. For labeling with Oregon Green™ 488 BAPTA-1 AM (OGB-1, Invitrogen, CA, USA), the slices were incubated at 36° C for 10-15 min in the standard extracellular solution containing (in mM): 125 NaCl, 4.5 KCl, 2 CaCl_2_, 1.25 NaH_2_PO_4_, 26 NaHCO_3_, 20 glucose (pH 7.4 when bubbled continuously with 95% O_2_ and 5% CO_2_) and 15 µM OGB-1^74^. Imaging of spontaneous network activity was performed at 32-34 °C, 512 x 276-pixel resolution, and 20 ms/frame using an MVX10 Research Macro Zoom Microscope (Olympus, Tokyo, Japan) equipped with an LED source for excitation (Thorlabs, central wavelength 470 nm), the Zyla 4.2 sCMOS camera (Andor Technology) and Andor Solis software for image acquisition. For each slice, six 5-minute-long consecutive videos were recorded.

For analyses, each video frame was first reduced in height and width by averaging 2×2 pixels. Then, fluorescence changes were extracted from videos using principal component analysis (PCA)^79^. To do so, the videos were reduced in dimension to the first 200 principal components, which were inspected by eye to separate components representing neuronal activity (unequivocal increases in fluorescence intensity) from artifacts (occasional movement of small bubbles, slice jittering, etc.). The refined data set was generated using only components representing neuronal activity. Regions of interest (ROIs; illustrated in Fig. 6a) were drawn manually in ImageJ based on *Facundo Valverde’s Golgi atlas of the postnatal mouse brain* (Springer, Wien, New York 1998) and imported into MATLAB (R2018b) for further analyses. Fluorescence traces for given ROIs were calculated by averaging the intensity of pixels inside each ROI in all video frames. A robust linear regression algorithm was used to detect the baseline fluorescence F_0_^80^. ΔF/F traces were calculated as (F-F_0_)/F_0_. Ongoing spontaneous events were detected for a given ΔF/F trace, and their peak amplitude was calculated using MATLAB’s peak detection function. Depending on their kinetic properties, detected events were classified as either a single transient or a burst (illustrated in Fig. 6e, insert). Events decaying to less than 10% of their peak amplitude before the beginning of the next event were classified as transients. All other events were considered bursts. All data recorded from one slice were pooled to calculate the frequency of bursts and their fraction among all events, and the median was taken as a representative value. The refined data was binarized for pacemaker analysis using manually set threshold values; each binary video was subdivided into shorter consecutive frames containing only one *no activity-activity-no activity* set of transitions, called an active window. The new active window started after at least 1 second without any activity. The second frame of the active window was projected onto the ROI map shown in Fig. 6a to determine the anatomical location of a pacemaker. The number of pacemakers detected in this ROI was divided by the total number detected in the entire slice to calculate the fraction of pacemakers located in a given anatomical region. For statistical analyses, the mixed design analysis of variance was used to model 2 individually measured dependent values, namely the frequency and the fraction of bursts. In the mixed design model, the brain regions were defined as a within-subject factor, and two independent groups (wildtype and mutant) were defined as a between-subject factor. For the fraction of pacemakers, data followed a non-normal distribution, and statistical comparison was done using the non-parametric Mann-Whitney Test with Bonferroni correction for multiple comparisons. JASP software (https://jasp-stats.org/) was employed for all statistical comparisons.

### In vitro Electrophysiology

All experiments were performed on *Scn2a*^p.A263V^ mice pups (postnatal days 10-15 or 23-30). Coronal slices (300 µm) were cut using a microtome (HM650 V; Microm International GmbH, Walldorf, Germany) incubated for 30 min at 34°C in a holding chamber. Slices were then allowed to recover for 1h at room temperature (R.T.; 19 – 23°C) in standard aCSF containing (in mM): 125 NaCl, 3.5 KCl, 1.25 NaH_2_PO_4_, 26 NaHCO_3_, 2.6 CaCl_2_, 1.3 MgCl_2_, and 15 glucose and equilibrated with 5% CO_2_: 95% O_2_. Slices were transferred to a recording chamber, where they were superfused (≥2.5 mL/min) with standard aCSF at 33-35°C. Cells within the pyramidal layer were visualized with infrared oblique illumination optics and a water immersion objective (60x, 0.9 NA, Olympus). Current-clamp whole-cell recordings from the soma of hippocampal CA3 pyramidal neurons were performed using a BVC-700 amplifier (Dagan Corporation, Minneapolis, Minnesota, USA). Data were filtered at 10 kHz and sampled at 50 kHz with a Digidata 1440 interface controlled by pClamp Software (Molecular Devices, Union City, CA). Patch-pipettes (2-6 MΩ) were filled with a solution containing 115 mM K-gluconate, 20 mM KCl, 10 mM Na phosphocreatine, 10 mM HEPES, 2 mM Mg-ATP, 0.3 mM Na-GTP (pH:7.3; osmolality 295 mOsm) complemented with 50 μM Alexa Fluor 594 and ∼0.1%–0.3% biocytin. The liquid-junction potential was calculated as −14 mV and subtracted from all voltages. Passive properties were determined at a membrane potential of –74 mV according to standard protocols using 800-ms-long subthreshold hyperpolarizing and depolarizing current injections. Active properties were determined from suprathreshold long current injections. The first action potential within 10 ms of the onset of the current injection was selected to analyze action potential properties. The action potential threshold was determined using the first peak in the second derivative^62^. The action potential peak was the peak voltage reached, and the amplitude was determined as the difference from the AP threshold.

The current clamp and nucleated patch clamp recordings in CA1 neurons were performed in the horizontal brain slices (350-400 µm) as previously described^75^. Cells were hold at –70 mV. The liquid junction potentials were not corrected. The intrinsic neuronal properties and single action potential parameters were analyzed as previously described^75^. The input resistance was determined by the slope of a linear regression fit to corresponding steady-state voltage responses plotted versus a series of current injections ranging from –110 to –10 pA with 10 pA increments. Only events with a voltage peak amplitude surpassing 0 mV were regarded as action potentials. The first evoked action potential was used to determine single action potential parameters. The peak amplitude of an action potential was measured from the baseline to the peak. The half-width of an action potential was determined at 50% of the peak amplitude. The threshold of an action potential was determined as the voltage at which the first derivative dV/dt reached 50 V/s.

### Electrocorticogram (ECoG) Telemetry Recordings

For ECoG recordings, adult mice of both sexes were implanted with telemetry transponders (PhysioTel ETA-F10, Data Sciences International). For analgesia, mice received 0.025 mg/kg buprenorphine (i.p., TEMGESIC, Indivior, UK Limited) and 5.0 mg/kg carprofen (s.c., Norbrook Laboratories Limited, Ireland) 30 min prior to surgery. Mice were anesthetized with 0.5–4% isoflurane in 100% oxygen and kept under 0.8–1.5% isoflurane anesthesia throughout the surgery. Body temperature was maintained at 36.5°C using a homeothermic heating pad (Stoelting, USA). Mice were placed into a stereotaxic device (Kopf Instruments, CA), and a midline skin incision was made above the skull. The skull was cleaned by treating it shortly with 10% H_2_O_2_, rinsed with 0.9% NaCl solution, and dried. A bonding agent (OptiBond, Kerr Dental, Germany) was applied to the skull surface. Using a dental drill, a craniotomy for the recording wire was drilled above the S1 cortex and hippocampus (AP –1.8; ML +1.8) and adjusted using bregma-lambda distance. For the reference wire, another craniotomy was performed above the cerebellum. The tips of the EEG leads were placed epidurally and fixed with dental cement (Tetric EvoFlow, Ivoclar, Liechtenstein). Transponders were implanted subcutaneously between the neck and abdomen, with the wires placed just below the nuchal fold and fixed with dental cement. The skin incision was then closed using cyanoacrylate skin glue (GLUture, Zoetis, USA). After the surgery, animals were kept on an electric heating pad and allowed to recover for 5 days. During that time, animals were closely monitored, scored, and received additional analgesics via the drinking water if needed. After the recordings, animals were sacrificed via transcardial perfusion, and the brain was explanted for immunohistochemistry. Animal cages were placed on receiver boards to detect the ECoG signal and movement activity. Synchronized video recordings were acquired simultaneously. Data were digitally stored using Ponemah (DSI, MN) and the Media Recorder (Noldus, Wageningen, The Netherlands). Animals were recorded for an average of 1 week, with some recordings extending up to 6 weeks. Custom MATLAB scripts (MathWorks, MA) were used to visualize EEG and generate multitaper spectrograms of the whitened signal. Seizures were manually identified, and their phases were delineated based on these visualizations. The resulting seizure annotations were then used to quantify the seizure characteristics. For the analysis of interictal activity, periods of rapid eye movement (REM) sleep, slow-wave sleep (SWS), and wakefulness were manually defined based on characteristic EEG frequency composition as visualized in the spectrograms. The concatenated EEG signal for each state was z-scored, and power spectral density (PSD) was calculated using custom MATLAB scripts. The background was subtracted using the fitting oscillations & one over f (FOOOF) method^76^.

### In vivo *LFP Recordings in Neonates*

Mouse pups of both sexes aged P2.5-P14 (*B6.Scn2a*) and P3-P12 (*CD1.Scn2a*), still nursed by mothers at the time of experimentation, were used for the recordings. Animals were injected with buprenorphine (s.c., 0.0125 mg/kg b.w., TEMGESIC, Indivior UK Limited, UK) 30 min prior to surgery. P2.5-P4 pups were anesthetized using cold anesthesia, and P5–P14 pups were anesthetized using 1.5–2% isoflurane in 100% oxygen. A midline incision was followed by the removal of the skin covering the top of the skull. The incision edges were treated with the local anesthetic (1 % bupivacaine, Bucain, Puren Pharma, Germany), and the periosteum was then denatured and briefly treated with 10% H_2_O_2_. The skull was rinsed with 0.9% NaCl solution and dried. A bonding agent (OptiBond, Kerr Dental, Germany) was applied to the skull surface. For a common ground electrode, an approximately 0.8-mm-wide craniotomy was drilled into the skull above the cerebellum, and a silver wire was inserted epidurally and fixed with dental cement (Tetric EvoFlow, ivoclar, Liechtenstein). A metal bar (ca. 2 mm in diameter) was attached with dental cement to the neck of the mouse, and the animal was subsequently head-fixed in the custom-made stereotactic apparatus. A constant temperature of 33 to 34° C was maintained to mimic nesting temperature using a homeothermic heating pad (Stoelting, USA). For the hippocampal recordings, burr holes were placed at 1.1-1.9 mm posterior to bregma and 1.3-1.6 mm lateral to midline, adjusted to age and bregma-lambda distance. Isoflurane anesthesia was discontinued after surgery. Linear 16-site silicon probes with a 50-µm distance between recording electrodes (A1×16-3mm-50-703, NeuroNexus, USA) or 32-site probes with a 25-μm distance between recording sites (A1×32-6mm-25-703-A32) were inserted in either of the hemispheres or bilaterally. The silicon probes were connected to a unity gain buffer amplifier (HS-18 or HS-36 headstage, Neuralynx, Inc., USA) mounted to the stereotaxic instrument (Stoelting, USA). Data were digitally filtered (0.5 – 8000 Hz bandpass) and digitized as 16-bit integers with a sampling rate of 32 kHz using the Digital Lynx 4SX acquisition system (Neuralynx Inc, USA). Breathing rate and movement during the recordings were measured using a piezoelectric sensor placed under the animal’s thorax and recorded in parallel with the same sampling rate. Data acquisition started 30 min after probe insertion. Before insertion, the silicon-probe shank was marked with the fluorescent dye DiI (Molecular Probes, Netherlands) to verify electrode positions post hoc in hippocampal coronal slices. The ventral hippocampal commissure (VHC) was stimulated using an MCS Stimulus Generator (STG4002, Multi-Channel Systems, Germany). Tungsten stereotrodes (WE3ST30.1A3, Science Products, USA) were placed 0.4 mm posterior to bregma and 0.2 mm lateral to midline, contralateral to the silicon probe in the dorsal hippocampus. Treatment with phenytoin or vehicle was administered by subcutaneous injection starting at P1 in *B6.Scn2a* (n = 3, 2×30 mg/kg BW/d in water) and *CD1.Scn2a* mice (n = 2, 10 mg/kg BW/d in 10% cyclodextrin).

All in vivo data were analyzed and visualized using MATLAB (Mathworks, MA) or Neuroscope (http://neuroscope.sourceforge.net/)^77^. Local field potentials (LFPs) were low-pass filtered and downsampled to 1.25 kHz from raw traces. Seizures were manually identified and marked based on LFP visualizations. Multitaper spectrograms of original and whitened signals were computed using Chronux (http://www.chronux.org) or wavelets (Gabor family) for additional visualization. For sharp-wave (SPW) analysis, current source densities (CSDs) were calculated as the second spatial derivative of 1-to 200-Hz–filtered LFPs. SPWs were automatically detected on CSD; the events were then clustered using UMAP^78^, resulting in three categories: SPWs (sink below pyramidal layer, source above), inverted SPWs (sink above pyramidal layer, source below), and unidentified events. The clustered dataset was manually curated, and SPW properties were then computed on this curated dataset using custom MATLAB scripts.

### In vivo *LFP Recordings in Adults*

For adult in vivo LFP recordings, adult mice of both sexes were implanted with magnetic head plates (Model 13, Neurotar, Finland). For analgesia, mice received 0.025 mg/kg buprenorphine (i.p., TEMGESIC, Indivior, UK Limited) and 5.0 mg/kg Carprofen (s.c., Norbrook Laboratories Limited, Ireland) 30 min prior to surgery. Mice were anesthetized with 0.5–4% isoflurane in 100% oxygen and kept under 0.8–1.5% isoflurane anesthesia throughout the surgery. Body temperature was maintained at 36.5°C using a homeothermic heating pad (Stoelting, USA). Mice were placed into a stereotaxic device (Kopf Instruments, CA), and a midline skin incision was made above the skull. The skull was cleaned briefly with 10% H_2_O_2_, rinsed with 0.9% NaCl solution, and dried. A bonding agent (OptiBond, Kerr Dental, Germany) was applied to the skull surface. Bregma-lambda distance and coordinates for future craniotomy above the dorsal hippocampus (AP –2; ML +1.5) were marked with a surgical marker. Using a dental drill, a craniotomy for the reference screw was made above the cerebellum. The screw, attached to the head plate with an insulated short wire, was then placed deep enough to touch the cerebrospinal fluid. The head plate was fixed to the animal’s skull with dental cement (Tetric EvoFlow, ivoclar, Liechtenstein). The skin around the head plate was then closed using a cyanoacrylate skin glue (GLUture, Zoetis, USA), and the top of the skull was covered with silicone (Kwik-Cast, WPI, USA). After the surgery, animals were kept on an electric heating pad and allowed to recover for at least 5 days before the recording. One day before the recording, a craniotomy for the silicon probe was performed using the marked coordinates under isoflurane anesthesia, following the same protocol as during the initial surgery. After the recordings, animals were sacrificed via transcardial perfusion, and the brain was explanted to verify the probe location.

All the recordings were done in Mobile HomeCage (MHC, Neurotar, USA), with the mouse freely moving in a floating cage on the air table. Before the first recording, mice were habituated for at least four days in the MHC. On the first day, mice were allowed to explore the cage freely. For each day that followed, they were head-fixed for 5, 15, and 30 min. Linear 32-site silicon probes were used with a 25-or 50 µm distance between the recording sites (A1×32-5mm-25-177-A32, A1×32-5mm-50-177-A32, NeuroNexus, USA). For probe insertion, mice were head-fixed using the magnetic head plate. The probe was inserted using the motorized Multi-Probe Micromanipulator System (NewScale MPM, USA) at 200 µm/min speed and 10 µm/min from 1 mm depth. The final depth was 1.6-1.8 mm for 25 µm probes and 2.0-2.4 mm for 50 µm probes. For optimal recording quality, silicone oil was applied to the craniotomy after probe insertion, and a minimum 30-minute period was observed before starting the recording. The silicon probes were connected to a unity gain buffer amplifier (HS-36 headstage, Neuralynx, Inc., USA). Data were digitally filtered (0.5 – 8000 Hz bandpass) and digitized as 16-bit integers with a sampling rate of 32 kHz using the Digital Lynx 4SX acquisition system (Neuralynx Inc, USA). Mouse position was recorded simultaneously and synchronized with the LFP data using MHC magnetic tracking. Each recording lasted about 1.5h. After recording, the probe was removed, and the skull surface was protected with Kwik-Cast. For each animal, 2 to 4 recordings were done, with at least one day passing between the recordings. Before insertion, the silicon probe shank was marked with the fluorescent dye DiI (Molecular Probes, Netherlands), DiO, or DiD (Biotium, USA) to verify electrode positions post hoc in hippocampal coronal slices.

All analyses of hippocampal LFP were made using custom-written MATLAB (MathWorks, MA) scripts. Periods of immobility and movements were classified by thresholding using the delta band (1–3 Hz) and theta (3–7 Hz)/delta ratios and mouse running speed data. The threshold was manually set for each recording. Because we found no significant differences between movement and immobility periods, only data calculated on movement periods for all the analyzed parameters (except sharp-wave ripple characteristics, only analyzed during immobility) are presented in Fig. 8. Delta and theta bands were computed as the absolute value of the Hilbert transform of the bandpass-filtered LFP and smoothed with a 10-sec quadratic kernel.

Band-limited power (BLP) analysis of the different strata in the dorsal hippocampus was performed for the following frequency bands: delta, 0.5–4 Hz; theta, 5–9 Hz; sigma, 11–16 Hz; beta, 16–29 Hz, low-gamma, 30–50 Hz; mid-gamma, 50–90 Hz; high-gamma, 90–130 Hz; ripple, 120–250 Hz; high-frequency oscillations (HFO), 250–600 Hz; and MUA, 700–3000 Hz. The original LFP at 32 kHz was downsampled to 1280 Hz (except for the MUA band), bandpass filtered, rectified, and resampled to 250 Hz. Hippocampal strata identification was assessed based on stereotaxic and electrophysiological markers such as the laminar distribution of the BLPs during movement and immobility, theta-gamma coupling as quantified by the modulation index (MI)^79^, ripple amplitude, theta phase shifts^80^, spiking activity, and post-hoc histologically verified. For a channel in the middle of each layer, a concatenated 1280-Hz signal for each state (immobility and running) was z-scored, and power spectral density (PSD) was calculated using custom MATLAB scripts. To analyze the difference in hippocampal oscillations and account for the 1/f component of the power spectrum, the background was subtracted using the fitting oscillations and one over f (FOOOF) method^76^. The theta phase shift was calculated between SP and GL for each movement period, and subsequent analyses of those periods were concatenated. For the MI laminar distribution, the recording site with the highest theta amplitude in SLM was selected as a reference to extract the theta phases. The tested amplitude-modulated frequency bands were low-, mid-, high-gamma, and HFO. Ripples were detected by bandpass filtering the CA1 SP recording channel at 100–220 Hz and thresholding at 5 SD of the absolute value of the Hilbert transform, with 2 SD threshold for ripple onset and offset, 15 ms interevent interval before merging, and events discarded based on the comparison with common noise and presence of the similar oscillations in SLM. Following detection, the frequency and duration of individual ripple events were calculated from time-resolved generalized Morse wavelets spectrograms of ±30 ms peri-threshold crossing windows.

### Statistics

Statistical analysis was performed using GraphPad Prism (versions 9-10, GraphPad). The significance level was set at P < 0.05. Parametric data are presented as mean ± standard deviation (SD), with error bars in figures representing SD. Nonparametric data are presented as median [first quartile, third quartile] and visualized using box and whisker plots. In these plots, the box represents the interquartile range (IQR) with the median line, while the whiskers extend to the minimum and maximum values within the range. No randomization method was used for this study. For c-Fos stereological analyses, investigators were blinded to the genotypes. Exclusion criteria for in vivo electrophysiology included bad ground connection, excessive 50 Hz or movement noise, and incorrect probe location. Sample sizes were determined based on previous experience and publications; all n values and specific statistical tests are indicated in the figure legends. Briefly, for normally distributed data, two-tailed Student’s t-test, paired t-test (two groups, independent or paired measures respectively), or one-way ANOVA with either Tukey’s, Bonferroni’s or Dunnett’s post-hoc tests (three groups) were used for normally distributed data. Nonparametric data were analyzed using the Mann-Whitney U test to compare medians between two groups, or the Kruskal-Wallis test with post hoc Dunn’s multiple comparisons was used for three groups. For repeated measurements, a two-way repeated-measures ANOVA or mixed-design ANOVA was performed, followed by Šídák’s multiple-comparisons test or simple main effect on group post hoc test as appropriate. For survival analysis, the Log-rank (Mantel-Cox) test was employed. The Wilcoxon signed-rank test was used to compare the observed results with a chance-level performance.

### Preparation of single-cell suspensions for scRNA-seq

Mice at the respective age (postnatal day = P4 or P7 or P14) and of appropriate genotypes were sacrificed by decapitation (pups at the age of P14 were decapitated under brief isoflurane anesthesia), the brain was rapidly removed and immediately transferred into ice-cold oxygenized (bubbled with carbogen: 95% O2, 5% CO2) artificial cerebrospinal fluid (aCSF: 125mM NaCl, 25mM NaHCO3, 1.25mM NaH2PO4, 2.5mM KCl, 0.05mM CaCl2, 6mM MgCl2, 2.5mM glucose, 50mM sucrose, 3mM kynureinic acid) for ages P4 and P7^46,81^. At the age of P14 NMDG-aCSF (96mM NMDG, 20mM HEPES, 10mM MgSO4, 0.5mM CaCl2, 25mM glucose, 5mM Na-Ascorbate, 3mM Sa-Pyruvate, 12mM N-Acetylcysteine, 3mM myo-inositol, 0.01 mM taurine, 2mM thiouria, 13.2mM trehalose, 3mM kynureinic acid, pH 7.3-7.4 adjusted with 1N HCl) ^40,82,83^, was used to prevent interneuron loss which begins to become substantial around P14 with standard aCSF. At ages P7 and P14, the brains were trimmed on a cooled metal block to contain the hippocampus, and 300-µm-thick slices were prepared on a sectioning vibratome (Leica VT1000S) in a cooled and carbogen-bubbled chamber containing the respective cutting solution (aCSF or NMDG-aCSF). With Dumont No.7 forceps, the region of interest was microdissected from the brain slices on a cooled metal block in the respective aCSF under a stereomicroscope to obtain dorsal CA3 and adjacent regions (distal part of the upper leaf of dentate granule cell layer, complete CA3, complete CA2 and proximal half of CA1). This strategy was used to obtain comparable cell numbers of the main excitatory neuron populations from the dorsal hippocampus (DGGC, CA3, CA1), without depleting any particular cell type. For age P4, the hippocampus was quickly dissected, and approximately 0.5mm-thick slices were prepared from the dorsal half under the dissecting stereomicroscope with a scalpel. Single-cell suspensions were prepared with the Papain dissociation system (Worthington) according to the manufacturer’s instructions. This procedure was performed in the presence of RNase inhibitor and blockers of synaptic transmission and neuronal activity in all solutions (except for the lower layer of ovomucoid inhibitor solution as a bottom layer for the final discontinuous gradient centrifugation) (RNasin: 400U/mL, 1 µM TTX, 20 µM D-AP5, 10 µM CNQX). Four samples containing different genotypes were processed per preparation day.

### 10x Genomics Single-cell Library Preparation and Sequencing

Cell capture, barcoding, reverse transcription, cDNA amplification, and library construction were performed according to manufacturer’s instruction (10X genomics, Chromium Next GEM Single Cell 3’ Reagent Kit v3.1 (Dual Index), whereas the first seven samples from age P7 were prepared with the V2 chemistry: Chromium single Cell 3’ Library & Gel Bead Kit v2) at the West German Genome Center (WGG), Cologne. Quality and molecular concentrations of the libraries were assessed on a tape station (Agilent Technologies), and sequencing was performed on an Illumina HiSeq 4000. Sample de-multiplexing, barcode processing, and single-cell gene UMI (unique molecular identifier) count matrix were performed with and generated by Cell Ranger Suite (10X Genomics, using versions 2.2.0 - 7.0.0) with murine reference genome (GRCm38/mm10).

### Single-cell RNA Sequencing Data Preprocessing

Raw feature-barcode matrices generated by Cell Ranger were preprocessed using the CRMetrics v0.3.0 pipeline^84^ for quality control. Briefly, CellBender v0.3.0^85^) was applied with appropriate --expected-cells and --total-droplets parameters based on the distribution of total unique molecular identifier (UMI) count per droplet for each sample to remove ambient RNA. The filtered count matrices were then processed with Scrublet v0.2.3 ^86^ and DoubletDetection v4.2^87^ with default parameters to detect doublets. Cells identified as doublets by either method were removed. Additionally, cells with fewer than 1,000 total UMI counts or a mitochondrial gene fraction exceeding 10% were excluded.

### Sample Integration, Clustering and Cell Type Annotation

Count matrices from all samples were integrated using Conos v1.5.0^88^. First, sample processing was performed using pagoda2 v1.0.11^89^ via the basicP2proc() function with default parameters, except for setting min.transcripts.per.cell = 1 and n.odgenes = 2000, to normalize expression values per sample. A joint cell-cell connectivity graph was then constructed using the buildGraph() function with score.component.variance = TRUE to establish weighted inter-sample cell-to-cell links. A two-dimensional UMAP embedding of the joint graph was generated using the embedGraph() function in Conos, employing functions from the uwot v0.1.16 package^90^, as implemented in leidenAlg v1.1.4^91^., was applied using the findCommunities() function in Conos to determine joint clusters across samples with a resolution parameter set to 1.

Clusters were annotated based on the expression levels of known cell-type-specific marker genes curated from the literature^38–46,92–95^ (the marker gene list will be uploaded as a supplementary file). After annotating major cell types, excitatory and inhibitory neurons were separately subsetted and re-clustered for annotation at medium and high resolution.

### Differential analysis for wildtype vs mutant conditions

Most differential analyses were performed using Cacoa v0.4.0 (https://github.com/kharchenkolab/cacoa) package for cross-condition analysis (see below). The detailed description of the methods for the package are described elsewhere^96^. Note that the tools from the package have been implemented originally^97^, and further successfully implemented by the developers at full-scale of the toolkit^47,98–100^.

### Cluster-based Expression Distance Analysis

Cluster-based expression distances between genotypes were estimated at both medium- and high-resolution annotations using the estimateExpressionShiftMagnitudes() function from the Cacoa package. The function was run with the meta parameter set to account for technical variation between different experimental batches and 10x Genomics sequencing chemistry versions, and top.n.genes = 1000 to focus on the top 1,000 most highly expressed genes.

Pseudo-bulk expression profiles were generated for each cell type by aggregating gene expression across cells within each sample. Expression shift magnitudes were calculated by computing correlation distances between samples of different genotypes. To remove technical artifacts, batch and sequencing chemistry effects were regressed out using a generalized linear model. Residuals from this model replaced the raw correlation distance values, ensuring that expression shifts reflect biological variation rather than technical confounding factors. Statistical significance was assessed via 1,000 random permutations of genotype labels within each cell type, generating a background distribution of expression shifts. Resulting *P* values were corrected for multiple testing using the Benjamini-Hochberg (BH) method.

### Graph-based Expression Shift Analysis

Graph-based expression shifts between genotypes were estimated using the estimateClusterFreeExpressionShifts() function in Cacoa with gene.selection = “expression” to prioritize highly expressed genes. This method was applied to the joint graph constructed from Conos, enabling expression shift calculations independent of predefined clusters.

For each cell, gene expression values were smoothed over its local graph-based neighborhood, and pairwise correlation distances were computed to quantify shifts between samples of different genotypes. Statistical significance was assessed via 500 random permutations of genotype labels, generating a null distribution of expression shifts. The final shift magnitudes were further processed using a median filter to align estimates with statistical significance measures.

### Cluster-based Compositional Data Analysis

Compositional differences in cell-type abundances at both medium- and high-resolution annotations between genotypes were assessed using the estimateCellLoadings() function in Cacoa with default parameters. Compositional Data Analysis (CoDA) was applied to evaluate shifts in cell-type proportions while addressing the inherent dependencies of compositional data.

Cell-type proportions were first transformed using the isometric log-ratio (ILR) transformation, enabling statistically valid comparisons of relative abundances. Linear discriminant analysis (LDA) was applied to extract compositional loadings and identify the most discriminative axes separating genotypes. Statistical significance was assessed through permutation testing with 1,000 bootstraps, generating a null distribution of cell-type loadings. Resulting *P* values were corrected using the BH method.

### Graph-based Differential Cell Density Estimation

Graph-based cell density estimation was performed using the estimateCellDensity() function in Cacoa with the method parameter set as “graph”. This method smooths cell density across the joint graph constructed by Conos. For each sample, a binary indicator matrix was first created, marking whether a given cell belonged to a specific sample. This matrix was then smoothed over the graph using a heat kernel filter with a decay parameter (beta = 30), enabling density estimates to account for the local neighborhood structure of the graph. The resulting sample-wise density estimates were normalized by the total number of cells in each sample to ensure comparability across different conditions.

To assess differential cell density between genotypes, the estimateDiffCellDensity() function was applied with type = “wilcox” to the estimated density matrix. This method calculates the Wilcoxon rank-sum statistic to compare density distributions between the reference and target groups at each graph node. The test outputs a signed Z-score, where positive values indicate increased cell density in the target genotype relative to the reference, and negative values indicate depletion. To evaluate statistical significance, permutation-based adjustments were performed (n.permutations = 400) by randomly shuffling sample labels and recalculating density differences, yielding an empirical background distribution. Finally, multiple testing correction was applied using the BH method to control the false discovery rate.

### Identification of Differentially Expressed Genes

Differential expression (DE) analysis was conducted at both medium- and high-resolution annotations using the estimateDEPerCellType() function in Cacoa. The Wald test implemented in DESeq2 v1.38.3 package^101^ (test = ‘DESeq2.Wald’) was used to identify differentially expressed genes between genotypes.

For each cell type, pseudo-bulk count matrices were constructed by aggregating gene expression values across cells within each sample. To ensure reliable DE detection, only cell types with at least 10 cells per sample (min.cell.count = 10) and genes expressed in at least 5% of cells within each cell type (min.cell.frac = 0.05) were included in the analysis. Independent filtering was enabled (independent.filtering = TRUE) to optimize statistical power by removing genes with low mean expression before multiple testing correction. Resulting *P* values were corrected using the BH method. Additionally, Z-scores were computed based on raw and adjusted *P* values to rank genes by their effect sizes.

### Gene Set Enrichment Analysis

Gene set enrichment analysis (GSEA) was performed using the estimateOntology() function with type = “GSEA” in Cacoa, leveraging the clusterProfiler v4.6.2^102^), DOSE v.3.24.2^103^ and fgsea v.1.24.0^104^ packages for functional enrichment analysis. This method ranks all genes tested by DESeq2 and tests whether predefined gene sets are significantly enriched at either end of the ranked list.

First, genes were ranked by their signed test statistic from the DE analysis, and the top 500 highly expressed genes (n.top.genes = 500) were selected for enrichment analysis. GSEA was conducted using the fgsea algorithm, accessed via the GSEA_internal() function from the DOSE package. This method applies a weighted Kolmogorov-Smirnov-like test to measure whether a specific gene set is overrepresented at the extremes of the ranked list. Functional annotation was performed using GO Biological Process terms, with gene sets retrieved from the org.Mm.eg.db v3.16.0 database. Statistical significance was assessed using the fgsea permutation-based approach, in which gene labels were randomly shuffled to generate an empirical background distribution of enrichment scores. Resulting *P* values were corrected using the BH method.

### Cell-cell Communication Inference

Cell–cell communications between different cell types, annotated at high resolution, were separately inferred for samples from different ages and genotypes using the CellChat v2.1.2 package^105^. CellChat models intercellular communication by integrating gene expression data with a curated ligand-receptor interaction database that includes known ligand-receptor interactions and cofactors. For each age group and genotype, joint normalized expression matrices were generated and used to construct CellChat objects. Standard preprocessing steps were applied following the CellChat pipeline with default parameters to ensure accurate inference. The CellChatDB.mouse database was used as the reference for ligand-receptor interactions, incorporating all available interaction categories. To maintain statistical robustness, interactions involving fewer than 10 cells were excluded from further analysis.

To infer significant cell-cell communication networks, CellChat computed the probability of ligand-receptor interactions by integrating gene expression levels with prior knowledge of known interactions. The functions identifyOverExpressedGenes(), identifyOverExpressedInteractions(), and projectData() were sequentially applied to identify overexpressed signaling components and project the data into a lower-dimensional space for improved interpretability. Low-confidence interactions were filtered out to refine the communication probability matrix, ensuring that only biologically meaningful interactions were retained. Communication probabilities were further refined by computing pathway-level interactions using computeCommunProbPathway(), creating an aggregated view of intercellular signaling at the pathway level.

### RNA Velocity-based Pseudotime Analysis

RNA velocity inference was performed separately for each major neuronal cell type at the medium-resolution annotation. First, spliced and unspliced UMI count matrices were generated using the velocyto v0.17.17 package^106^). The joint count matrices, aggregating all samples per neuronal cell type, were subsequently processed using the velociraptor v1.8.0 R wrapper for scVelo v0.2^107^ to estimate RNA velocity using the dynamical model.

The velocity estimation process included multiple computational steps. First, count matrices were normalized by library size, and RNA velocity inference was performed using the scvelo() function in velociraptor with use.theirs = TRUE to leverage scVelo’s built-in preprocessing pipeline. The dynamical model was applied to infer transcriptional kinetics, followed by computation of a velocity transition matrix to predict future cell states. Per-cell RNA velocity pseudotime was inferred within scvelo(), providing a relative measure of cellular progression along the inferred trajectory. The resulting velocity vectors were projected onto a Conos-generated UMAP embedding using embedVelocity(), aligning velocity vectors with the original cell embeddings. The velocity field was further summarized using gridVectors(), which aggregates velocity information into a grid structure for downstream visualization. Finally, we compared the velocity pseudotime distributions of cells across major neuronal cell types using the two-sided Kolmogorov-Smirnov test and visualized the corresponding empirical cumulative distribution functions (ECDFs).

## Acknowledgments

We thank Dr. Igor Jakovcevski for his help with quantitative immunohistochemical and immunofluorescence analyses in the neonatal hippocampus. We also acknowledge the animal facilities’ support with excellent mouse care (Prof. Esther Mahabir-Brenner (ARTs team) and Dr. Maria Guschlbauer (Medical Faculty team), and excellent support for library generation and NGS sequencing from the team of the Cologne Center for Genomics (CCG). This work was supported by the DFG Research Infrastructure West German Genome Center (407493903) as part of the Next Generation Sequencing Competence Network (project 423957469). NGS analyses were carried out at the production site West German Genome Center Cologne. The work was supported by grants from the Deutsche Forschungsgemeinschaft (DFG): FOR 2715: D.I., H.L., O.G, T.K., and H.B.; SFB 1089 to H.B., S.L.M. and D.I., and SPP 2041 Computational Connectomics of the DFG to H.B.; from the iBehave Network (Ministry of Culture and Science of the State of North Rhine-Westphalia) to D.I. and H.B., from the Deutscher Akademischer Austauschdienst (DAAD, Research Grants - Doctoral Programmes in Germany, 2020/21, ID 57507871) to Y.R.; from the Novo Nordisk Foundation Hallas-Møller Ascending Investigator Grant (NNF21OC0067146) to K.K.; and from the European Union’s Horizon 2020 research and innovation program under the Marie Sklodowska-Curie grant agreement No. 101034291 (DISCOVER) to X.X.

