## supplementary Figures 1-9 for "Early postnatal CA3 hyperexcitability drives hippocampal development and epileptogenesis in SCN2A developmental and epileptic encephalopathy"

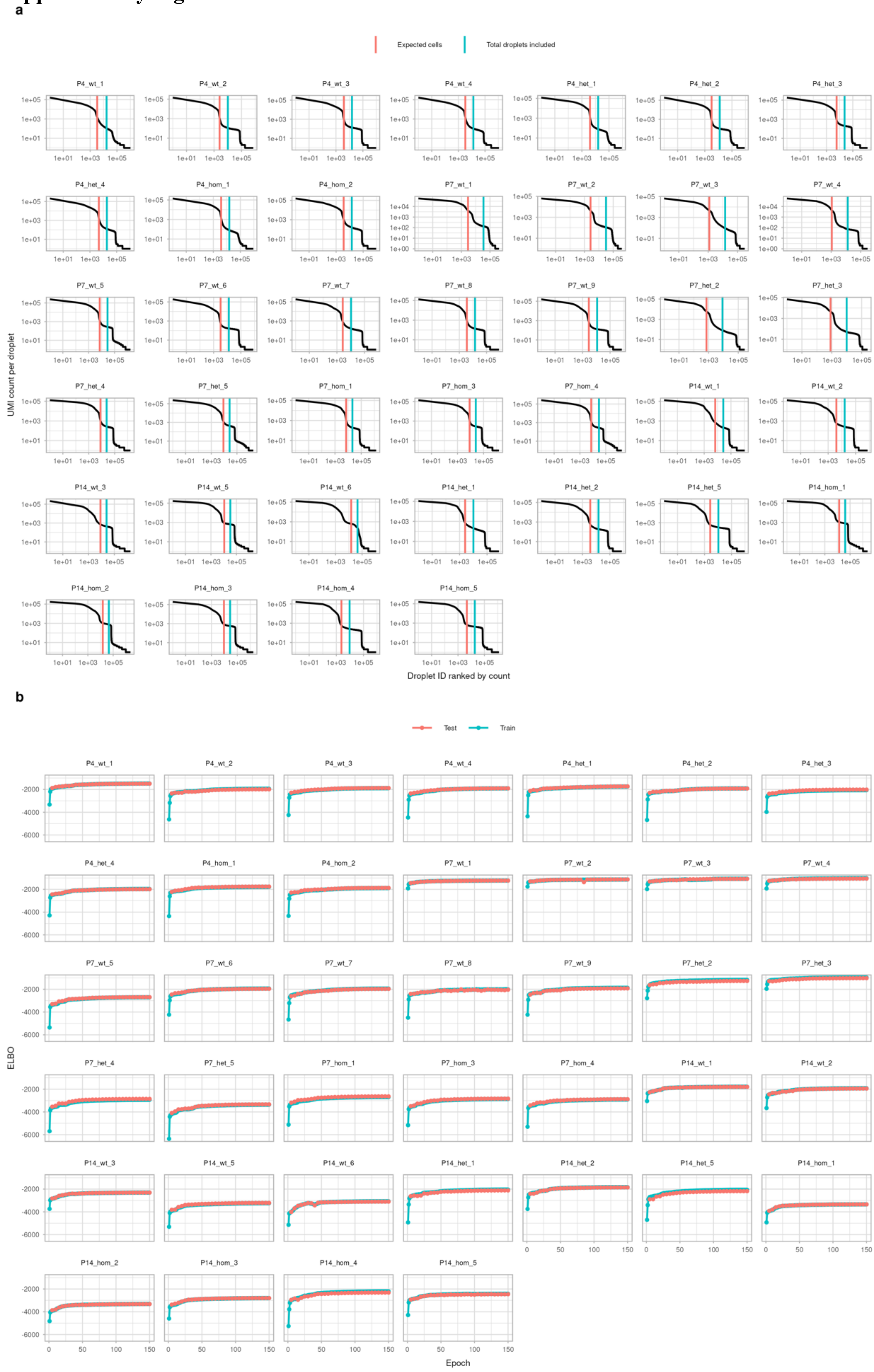

**Supplementary Fig. 1 | Ambient RNA removal using CellBender. a,** Barcode rank plots of raw 10x Genomics data from all individual samples, showing total UMI counts per droplet ranked by decreasing count. The red vertical lines indicate the expected number of recovered cells by Cell Ranger (--expected-cells), and the blue lines indicate the total number of droplets included for CellBender processing (--total-droplets). **b,** Evidence of lower-bound (ELBO) curves from CellBender training across all samples. The ELBO values for training and test datasets are shown across epochs, indicating convergence of the model during ambient RNA removal.

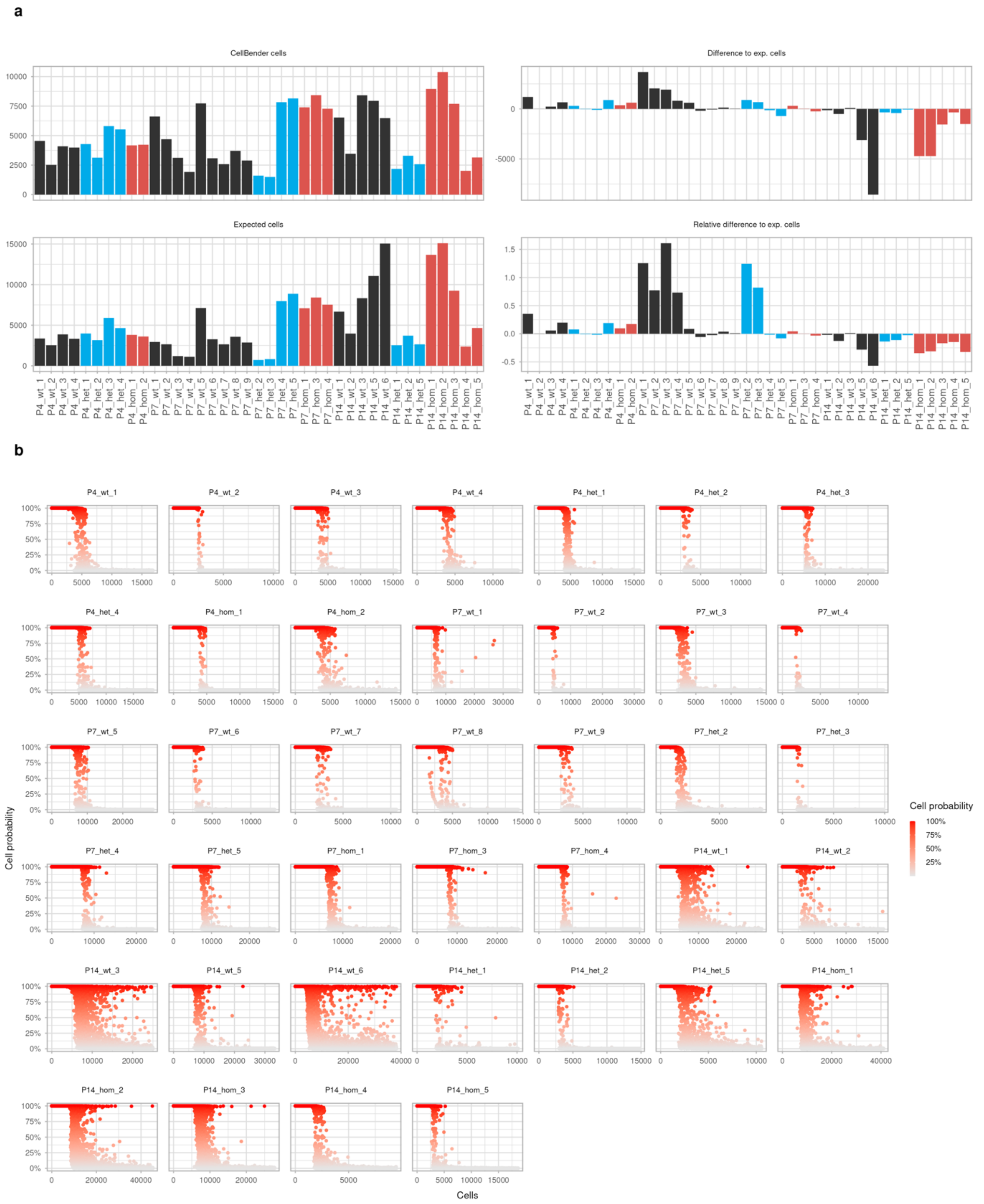


**Supplementary Fig. 2 | Summary of CellBender filtering performance across samples. a,** Summary bar plots showing the number of cells retained after CellBender filtering (top left), expected number of cells by Cell Ranger (bottom left), and absolute and relative differences between Cell Ranger-expected and CellBender-retained cell numbers (top and bottom right) for each sample. **b,** Droplet probability plots across samples as computed by CellBender, where each point represents a droplet, colored by the predicted cell probability. The plots show the separation of real cells from background droplets based on the learned model.

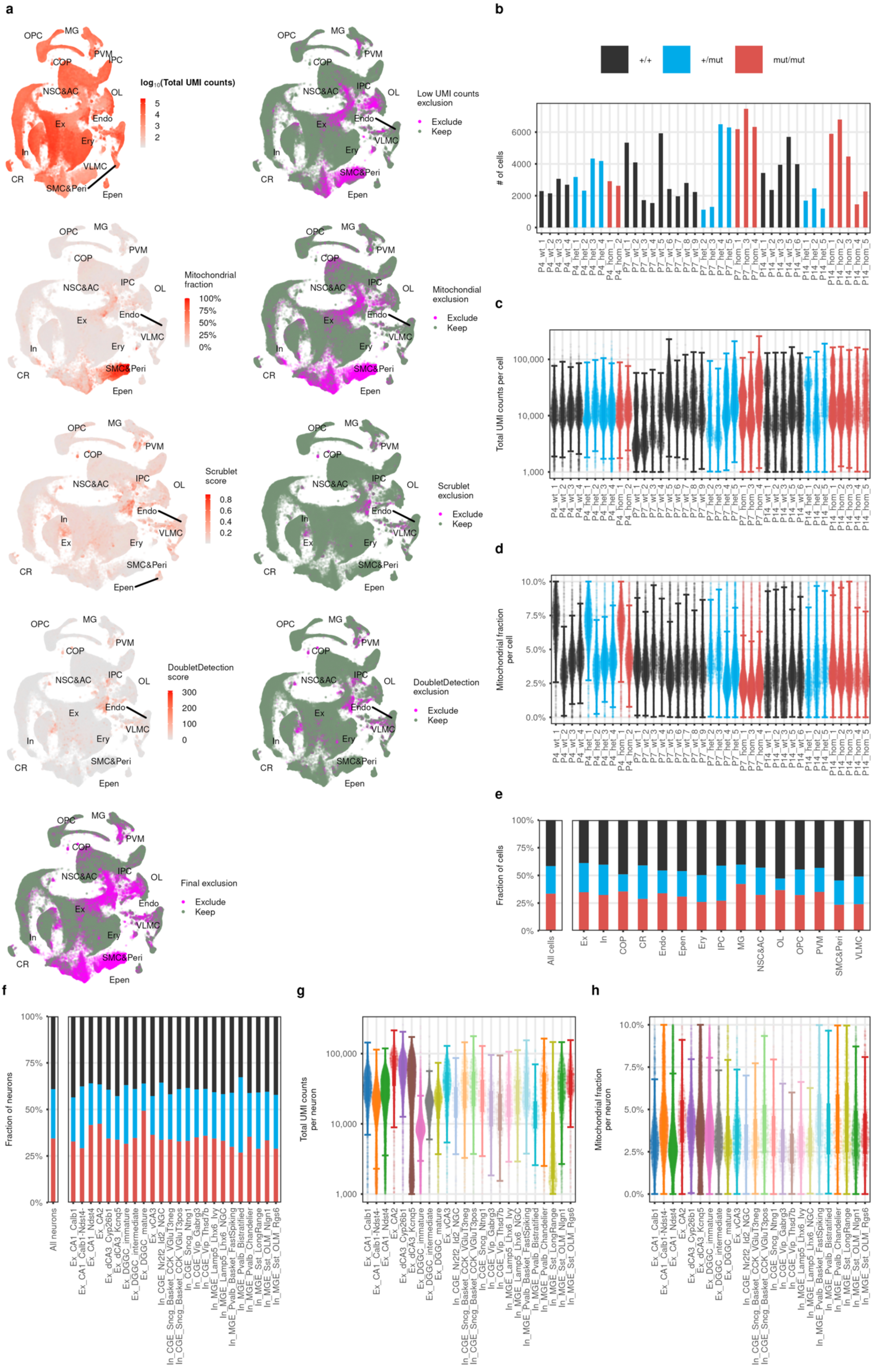
a
OPC
MG
COP
PVM
IPC
NSC&AC
OL
Endo
Ex
Ery
In
VLMC
CR
SMC&Peri
Epen
log10(Total UMI counts)
Low UMI counts exclusion
Exclude
Keep
Mitochondrial fraction
100%
75%
50%
25%
0%
Mitochondrial exclusion
Scrublet score
Scrublet exclusion
DoubletDetection score
DoubletDetection exclusion
Final exclusion
b
+/+
+/mut
mut/mut
### of cells
c
Total UMI counts per cell
d
Mitochondrial fraction per cell
e
Fraction of cells
All cells
f
Fraction of neurons
All neurons
g
Total UMI counts per neuron
h
Mitochondrial fraction per neuron

**Supplementary Fig. 3 | Overview of quality control metrics and cell filtering. a,** UMAP plots showing various quality control metrics across all cells before filtering. From top to bottom: Total UMI counts, mitochondrial gene fraction, Scrublet doublet score, and Doublet Detection score. Cells marked in pink were flagged for exclusion based on threshold criteria. **b,** Number of cells per sample after filtering, grouped by genotype. **c,** Violin plot showing total UMI counts per cell across samples. **d,** Violin plot showing mitochondrial gene fraction per cell across samples. **e,** Bar plot showing the proportion of retained cell types (low resolution) per sample, grouped by genotype. **f,** Fraction of retained neuron types (high resolution) per sample by genotype. **g,** Violin plot showing total UMI counts across retained neuron types (high resolution). **h,** Violin plot showing mitochondrial gene fraction across retained neuron types (high resolution).

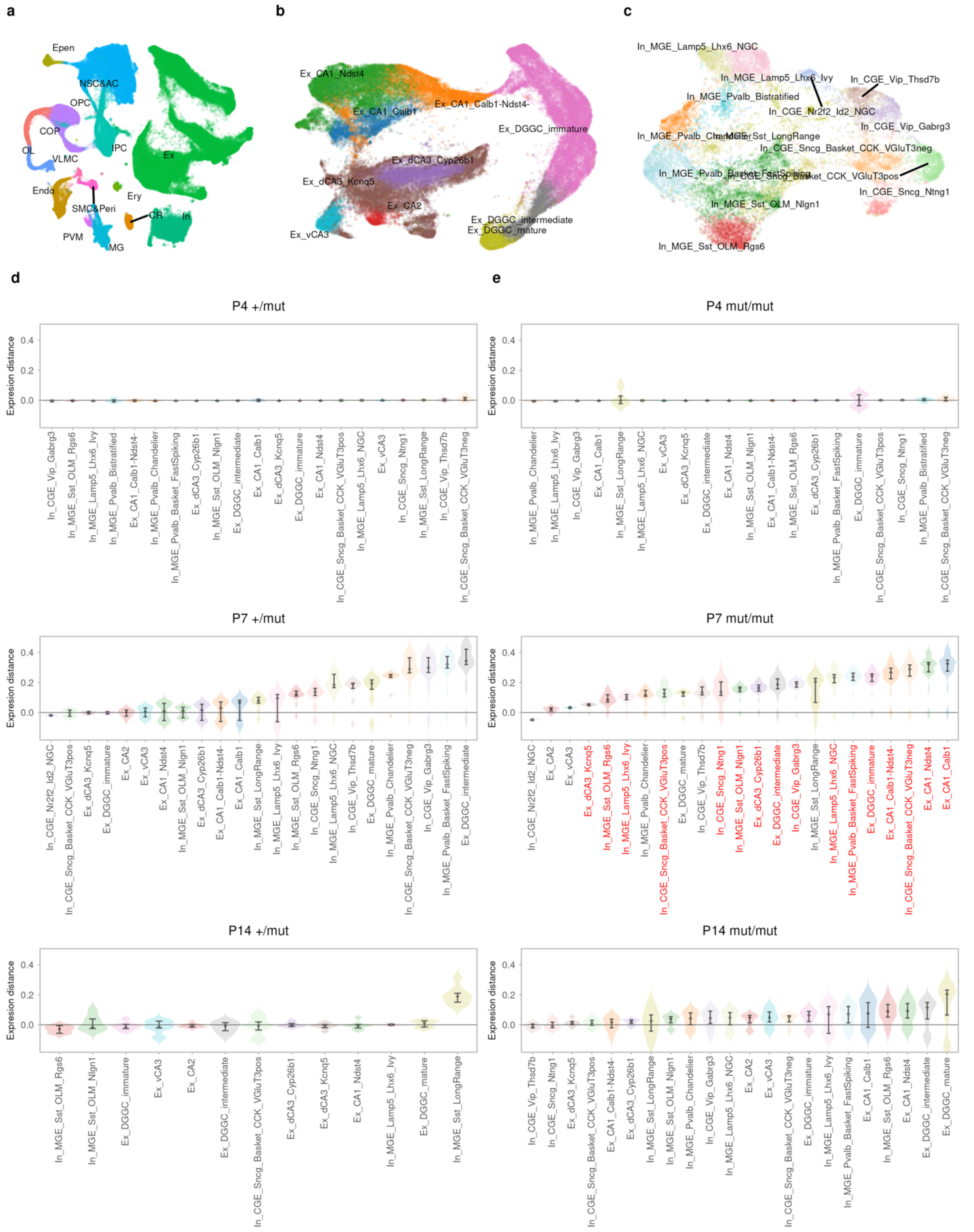


**Supplementary Fig. 4 | UMAP visualization and transcriptional changes in neuronal subtypes in *Scn2a* mutant mice.** **a,** UMAP representation of all analyzed cells, colored by low-resolution annotation. **b,** UMAP representation of analyzed excitatory neurons, colored by high-resolution annotation. **c,** UMAP representation of analyzed inhibitory neurons, colored by high-resolution annotation. **d,** Violin plots showing expression distance as a measure of the overall magnitude of transcriptional changes in neurons between +/*mut* and +/+ mice at P4, P7, and P14 for high-resolution annotations. Neuronal subtypes are ordered from smallest (left) to largest (right) mean transcriptional change. **e,** Same as **d** for *mut*/*mut* versus +/+ mice. Neuronal subtypes showing significant changes (BH-adjusted *P* value < 0.05) are marked in red.

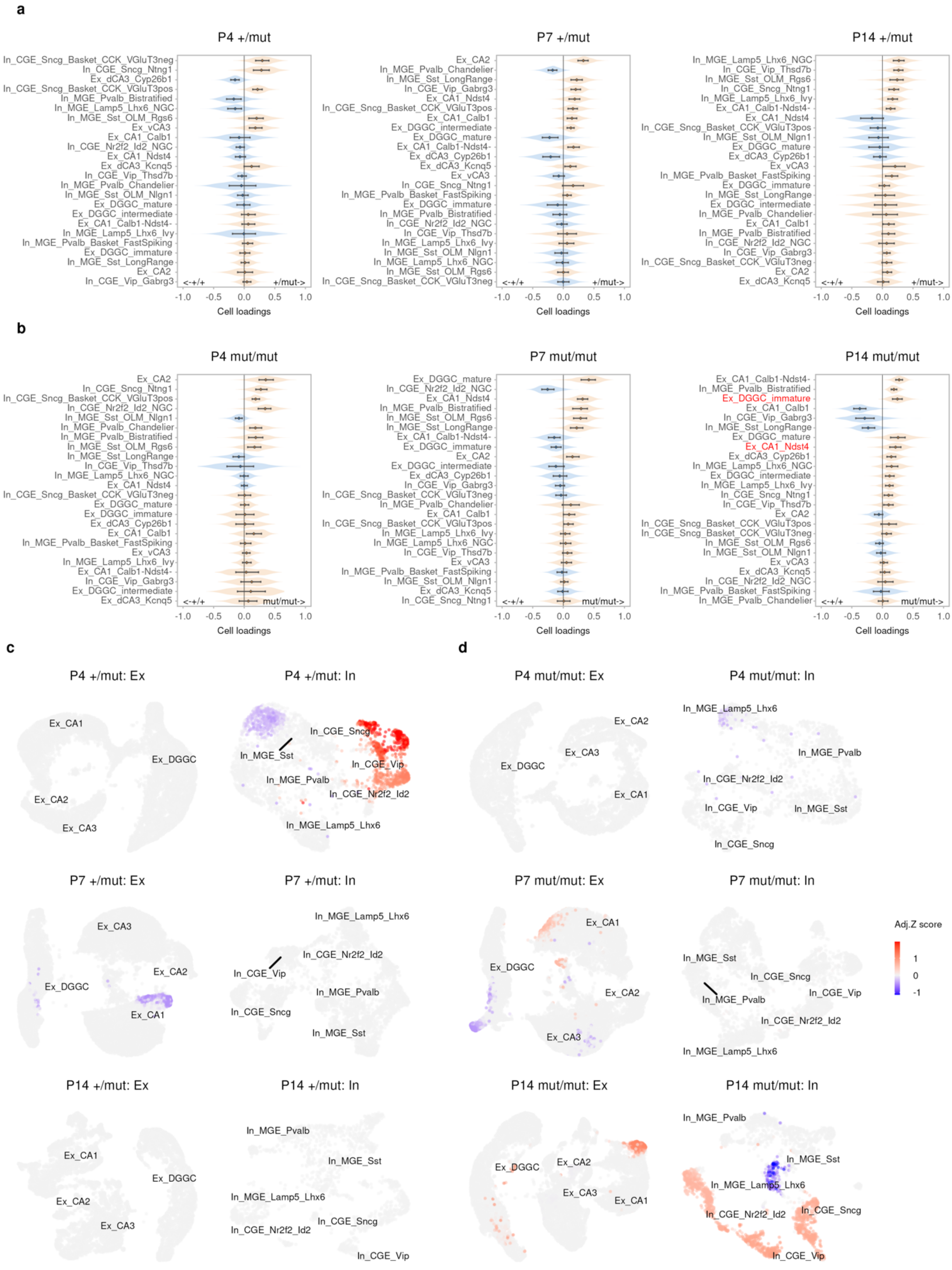


**Supplementary Fig. 5 | Neuronal compositional changes in *Scn2a* mutant mice. a,** Violin plots showing neuronal compositional changes analyzed by compositional data analysis (CoDA) between +/*mut* and +/+ mice for high-resolution annotation. Neuronal subtypes with positive cell loadings (colored in pumpkin orange) show increased abundance in +/*mut* mice, while those with negative cell loadings (colored in dodge blue) show increased abundance in +/+ mice. **b,** Same as **a** for *mut*/*mut* versus +/+ mice. Neuronal subtypes showing significant changes (BH-adjusted *P* value < 0.05) are marked in red. **c,** UMAP plots displaying changes in cellular abundance in Scn2a mutant cell types as differences of cell density on joint graphs of excitatory and inhibitory neurons between +/*mut* and +/+ mice at P4, P7, and P14, colored by adjusted significance levels. **d,** Same as **c** for *mut*/*mut* versus +/+ mice.

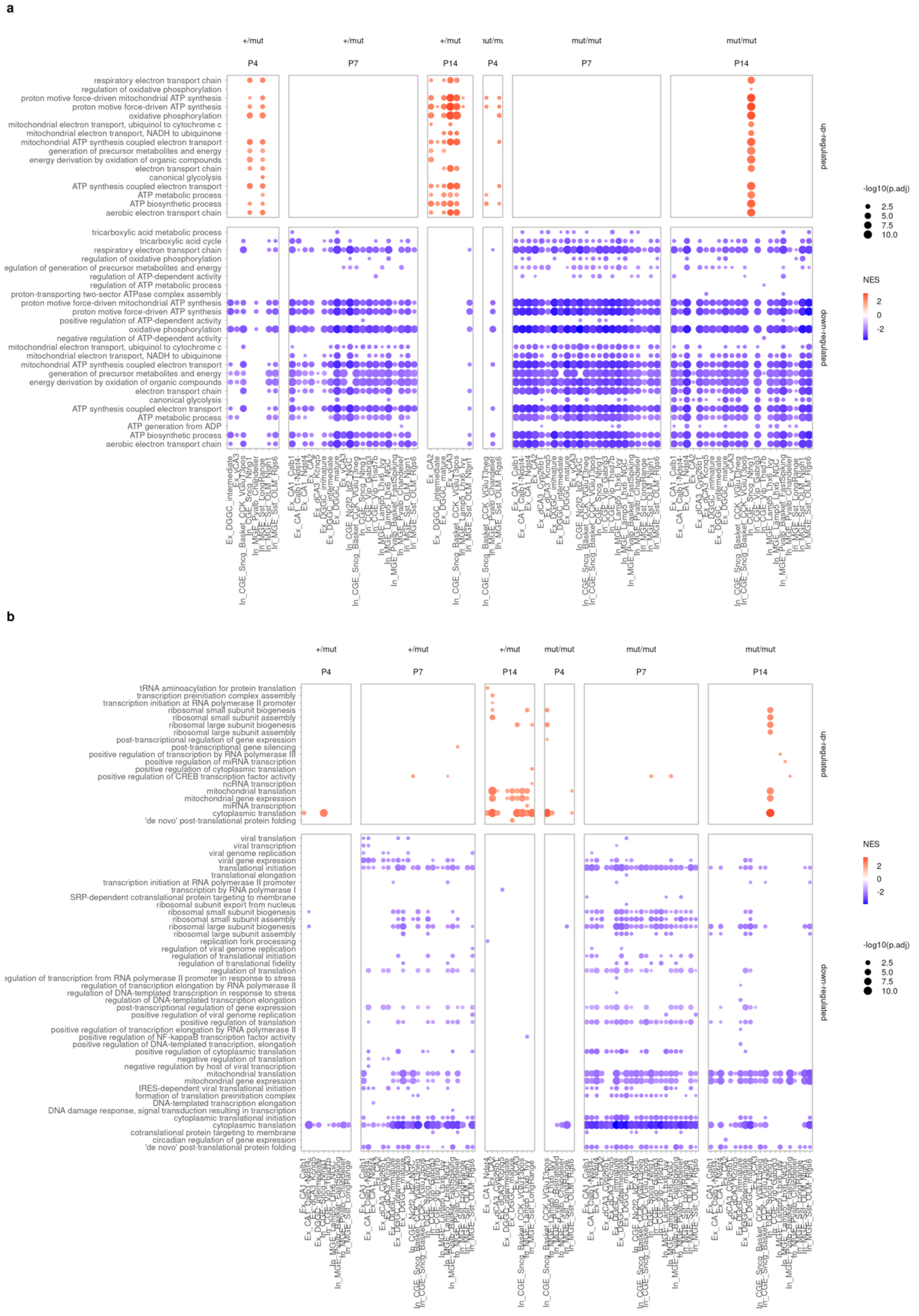
a
+/mut
mut/mut
P4
P7
P14
respiratory electron transport chain
regulation of oxidative phosphorylation
proton motive force-driven mitochondrial ATP synthesis
proton motive force-driven ATP synthesis
oxidative phosphorylation
mitochondrial electron transport, ubiquinol to cytochrome c
mitochondrial electron transport, NADH to ubiquinone
mitochondrial ATP synthesis coupled electron transport
generation of precursor metabolites and energy
energy derivation by oxidation of organic compounds
electron transport chain
canonical glycolysis
ATP synthesis coupled electron transport
ATP metabolic process
ATP biosynthetic process
aerobic electron transport chain
tricarboxylic acid metabolic process
tricarboxylic acid cycle
regulation of ATP-dependent activity
regulation of ATP metabolic process
proton-transporting two-sector ATPase complex assembly
positive regulation of ATP-dependent activity
negative regulation of ATP-dependent activity
ATP generation from ADP
up-regulated
down-regulated
-log10(p.adj)
2.5
5.0
7.5
10.0
NES
2
0
-2
b
tRNA aminoacylation for protein translation
transcription preinitiation complex assembly
transcription initiation at RNA polymerase II promoter
ribosomal small subunit biogenesis
ribosomal small subunit assembly
ribosomal large subunit biogenesis
ribosomal large subunit assembly
post-transcriptional regulation of gene expression
post-transcriptional gene silencing
positive regulation of transcription by RNA polymerase III
positive regulation of miRNA transcription
positive regulation of cytoplasmic translation
positive regulation of CREB transcription factor activity
ncRNA transcription
mitochondrial translation
mitochondrial gene expression
miRNA transcription
cytoplasmic translation
'de novo' post-translational protein folding
viral translation
viral transcription
viral genome replication
viral gene expression
translational initiation
translational elongation
transcription by RNA polymerase I
SRP-dependent cotranslational protein targeting to membrane
ribosomal subunit export from nucleus
replication fork processing
regulation of viral genome replication
regulation of translational initiation
regulation of translational fidelity
regulation of translation
regulation of transcription elongation by RNA polymerase II
regulation of DNA-templated transcription in response to stress
regulation of DNA-templated transcription elongation
positive regulation of viral genome replication
positive regulation of translation
positive regulation of transcription elongation by RNA polymerase II
positive regulation of NF-kappaB transcription factor activity
positive regulation of DNA-templated transcription, elongation
negative regulation of translation
negative regulation by host of viral transcription
IRES-dependent viral translational initiation
formation of translation preinitiation complex
DNA-templated transcription elongation
DNA damage response, signal transduction resulting in transcription
cytoplasmic translational initiation
cotranslational protein targeting to membrane
circadian regulation of gene expression
Ex_CA1_Calb1
Ex_CA1_Ndst4
Ex_CA2
Ex_CA3
Ex_dCA3_Cyp26b1
Ex_dCA3_Kcnq5
Ex_DGGC_immature
Ex_DGGC_intermediate
Ex_DGGC_mature
In_CGE_Lamp5_Lhx6_NGC
In_CGE_Sncg_Basket_CCK_VGluT3neg
In_CGE_Sncg_Basket_CCK_VGluT3pos
In_CGE_Sncg_Ntng1
In_CGE_Vip_Gabrg3
In_CGE_Vip_Htr3a
In_MGE_Pvalb_Basket_FastSpiking
In_MGE_Pvalb_Chandelier
In_MGE_Sst_LongRange
In_MGE_Sst_OLM_Ntng1
In_MGE_Sst_OLM_Rgs6

**Supplementary Fig. 6 | Changes in GO Biological Processes related to energy metabolism and DNA, RNA, and protein synthesis in Scn2a mutant mice. a,** Dot plots showing significantly enriched GO BP terms associated with energy metabolism in the top up- and down-regulated genes across neuronal subtypes (high-resolution annotations) between mutant and wildtype mice at P4, P7, and P14, colored by NES and sized by – log10(BH-adjusted *P* value). **b,** Same as **a** for GO BP terms associated with DNA, RNA, and protein synthesis.

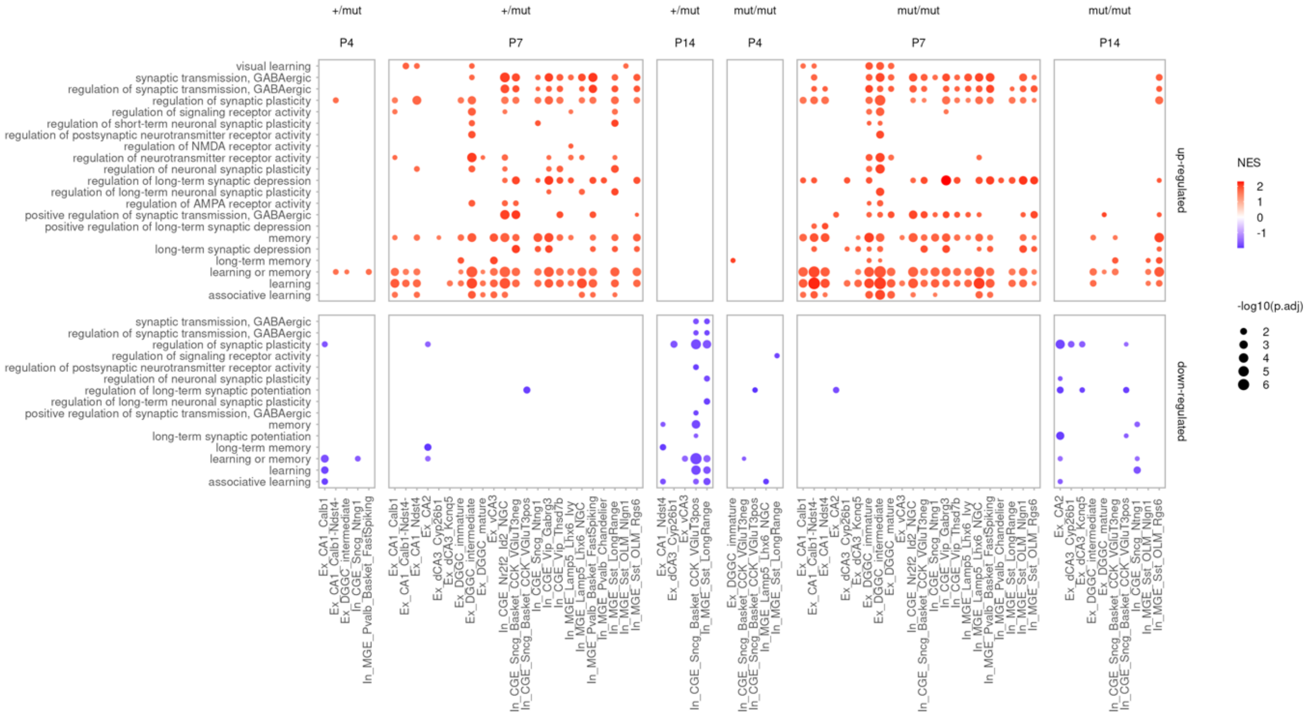


**Supplementary Fig. 7 | Changes in GO Biological Processes related to learning and memory in Scn2a mutant mice.** Dot plots showing significantly enriched GO BP terms associated with learning and memory in the top up- and down-regulated genes across neuronal subtypes (high-resolution annotations) between mutant and wildtype mice at P4, P7, and P14, colored by NES and sized by –log10(BH-adjusted *P* value).

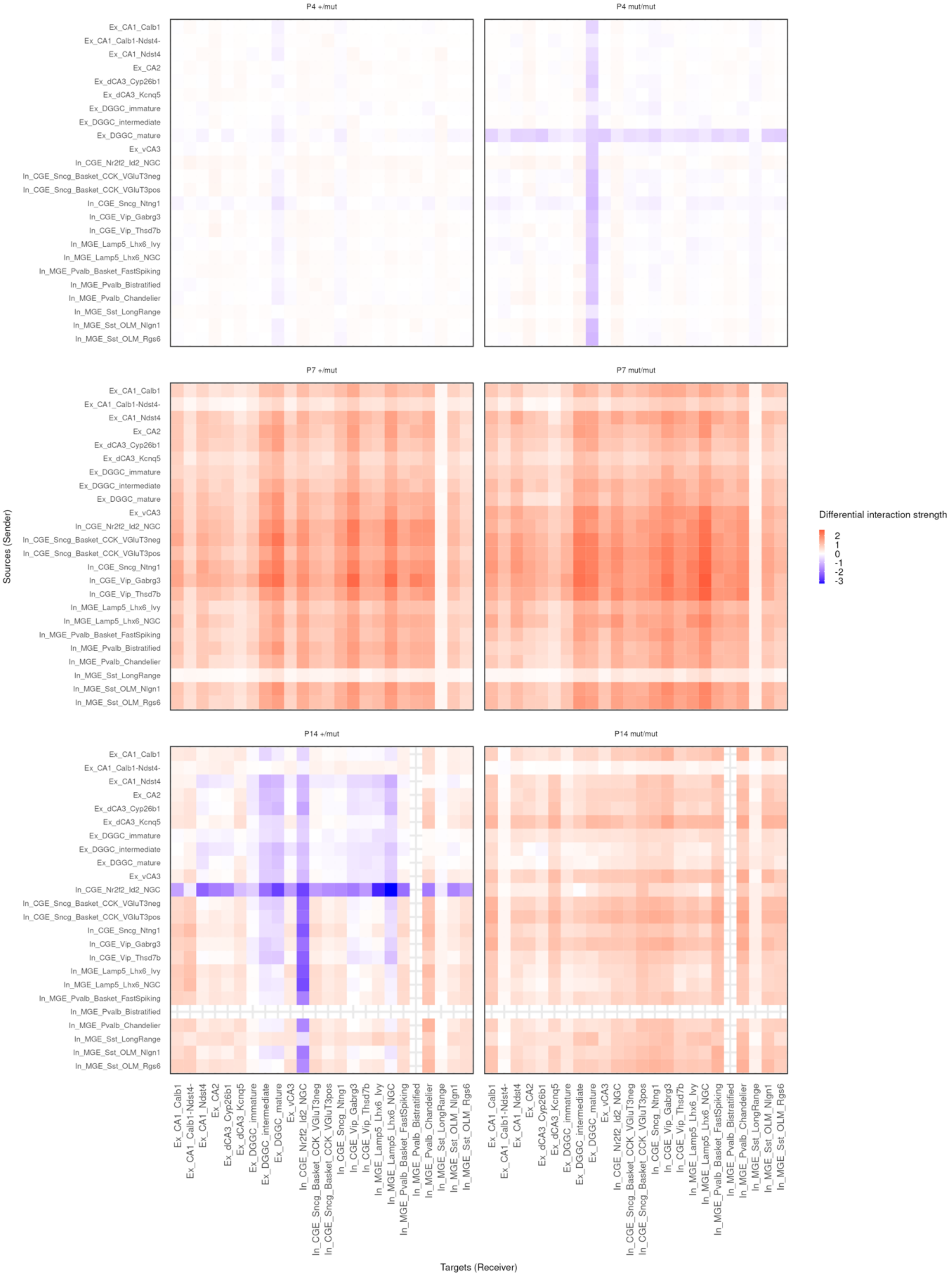


**Supplementary Fig. 8 | Differential intercellular communication networks inferred by CellChat**. Heatmaps showing differences in cell–cell communication strength between genotypes at P4, P7, and P14, as inferred by CellChat. Each matrix represents the differential interaction strength between source (sender) and target (receiver) cell types annotated at high resolution. Blue indicates weaker interactions in mutant relative to wildtype mice, and red indicates stronger interactions in mutant mice.

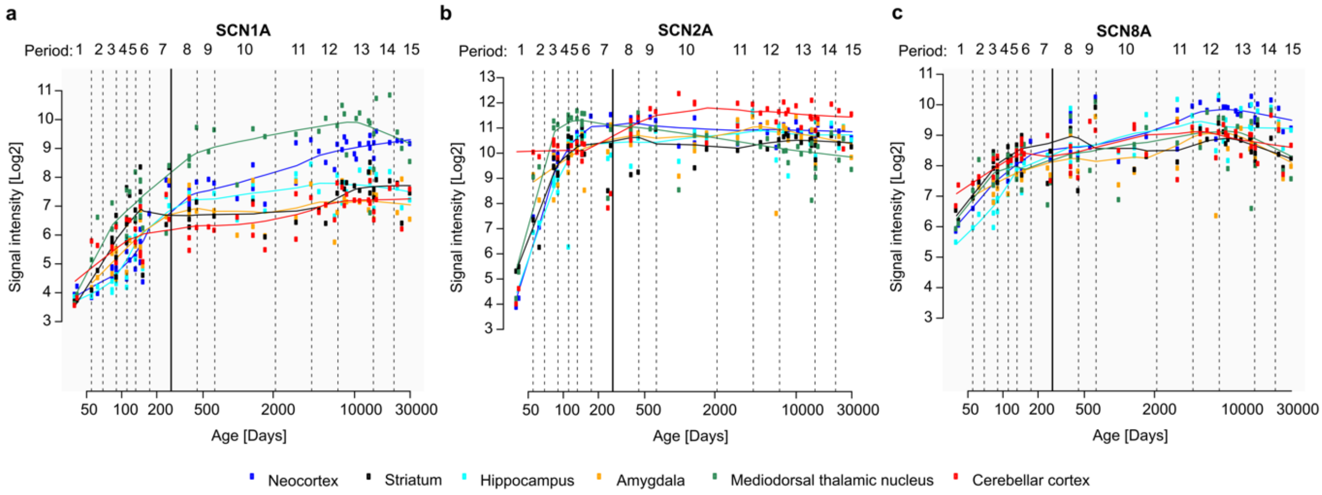


**Supplementary Fig. 9: Spatiotemporal expression of Nav subunit transcripts in the human brain.** The expression levels of the **a,** SCN1A, **b,** SCN2A, and **c,** SCN8A transcripts during human brain development were obtained from the online dataset of Kang *et al.*, 2011[1] available at http://hbatlas.org/pages/hbtd. Age is given in post-conceptional days (late embryonic development (4-8 weeks post conception, period 1), fetal development (periods 2–7), postnatal development (periods 8–12), and adulthood (periods 13–15)), the gray vertical line indicates the time point of full-term birth, expression levels are given as array signal intensity[1]. The brain regions are listed in the legend.
